# Scalable multi-group nonnegative spatial factorization for spatial genomics data with cell-type heterogeneity

**DOI:** 10.64898/2026.06.29.735224

**Authors:** Luis Chumpitaz-Diaz, Priyanka Shrestha, Barbara E Engelhardt

## Abstract

Spatial transcriptomics (ST) technologies enable the study of gene expression within the spatial context of tissues, providing insights into tissue structure, cellular interactions, and disease progression. However, existing dimension reduction methods often overlook spatial information or struggle to distinguish spatial gene patterns from those driven by cell-type differences, limiting biological interpretability by convolving differences in gene expression patterns with differences in cell-type proportions. To address these challenges, we introduce the scalable multi-group nonnegative spatial factorization (smNSF), a computationally-tractable probabilistic framework that integrates spatial coordinates and cell-type labels into a unified matrix factorization model. By using multi-group Gaussian processes (MGGPs) as priors, our model captures complex spatial variation in a cell-type specific way while enforcing nonnegativity to enhance interpretability. We develop a variational inference framework for MGGPs that supports scalable optimization and improves the numerical stability of smNSF. Across seven spatial transcriptomics datasets spanning diverse technologies and tissues, smNSF recovers sparse, interpretable spatial factors and, through its cell-type conditional posteriors, organizes them into cell-type enriched, cell-type specific, and universal spatial programs that are not apparent from marginal factors alone. Given cell-type labels in ST data, smNSF enables cell-type aware spatial decompositions and supports cell-type conditional posteriors for *in silico* exploration of relationships between spatial patterns and cellular identity.

**Author summary:** Most current analysis methods for spatial transcriptomics either ignore spatial structure or fail to separate gene-driven spatial patterns from cell-type driven differences. In this work, we develop a method that uses spatial coordinates and cell-type labels together to better uncover patterns of gene expression. Our approach, scalable multi-group nonnegative spatial factorization (smNSF), uses Gaussian processes to model spatial structure, which we extend to capture both spatial structure and cell type within a unified framework called multi-group Gaussian processes.

Applying smNSF to spatial transcriptomics datasets from mouse brain and human tissues, we find that conditioning on different cell types reveals spatial patterns that are invisible in standard analyses: some cell types suppress a pattern, others sharpen it, and some reveal structure that only emerges after conditioning. This helps us understand how cell-type specific spatial programs contribute to tissue organization.

To make this analysis tractable on the scale of modern spatial experiments, we also introduce a new computational approximation for Gaussian processes that is directly applicable in principle to other latent variable GP models. Together, these tools help disentangle biological sources of variation and support *in silico* exploration of how gene expression might change under different tissue or cell-type compositions.

## 1 Introduction

Spatially resolved transcriptomics (ST) technologies enable the measurement of gene expression while preserving spatial coordinates of those genes in tissue sections. This spatial information is crucial for understanding how gene activity varies across different tissue regions [1], how cells interact with their neighbors [2], and how multicellular structure and function emerge in health and disease [3]. Such insights have been instrumental in the study of embryonic development [4–6], tumor progression [7, 8], and immune compartmentalization [9, 10].

Dimension reduction methods such as principal component analysis (PCA), factor analysis (FA) [11], and nonnegative matrix factorization (NMF) [12] are commonly used to analyze high-dimensional single-cell RNA-sequencing (scRNA-seq) data [13–15].

These methods can also be applied to ST datasets, but they disregard spatial information.

More recently, spatially aware models such as MEFISTO [16], SpatialPCA [17], and nonnegative spatial factorization (NSF) [18] use Gaussian processes (GPs) [19] or related kernel-based priors to capture smooth spatial variation. MEFISTO uses an explicit GP formulation, while SpatialPCA uses a multivariate normal prior with spatial kernel-defined covariance, which is functionally similar to a GP. While these models recover spatial structure, their dense real-valued factors may be difficult to interpret. NSF improves interpretability by enforcing nonnegativity via exponentiated GP priors, yielding sparse, parts-based decompositions [18]; mNSF [20] extends NSF to multi-sample analysis through alignment-free sample-specific spatial correlation modeling. Other nonnegative spatial methods, including spatial LDA [21], neighborhood NMF [22], and FISHFactor [23], similarly produce interpretable spatial patterns. However, none of these methods — nonnegative or otherwise — take cell-type labels as input, limiting their ability to distinguish spatial patterns driven by gene-expression changes across space from those driven by cell-type-proportion changes within tissues.

More modern methods such as CELESTA [24] consider spatial context when inferring cell types. Meanwhile, methods such as cell2location [9], DestVI [25], and C-SIDE [26] focus on estimating cell-type composition across spatial locations using reference single-cell data and spatial information. However, these approaches are designed to recover cell-type labels, not spatially structured gene expression patterns.

A *spatial gene expression pattern*, as recovered by spatially-aware dimension reduction methods [16–18], captures changes in the expression levels smoothly across a tissue slice. Without explicit modeling and consideration of cell types, it is impossible to know when spatial changes in gene expression are due to expression changes within a cell type across the tissue, or changes in cell-type proportions across the tissue, because cell types generally express different sets of genes [27]. Deconvolving the impact of both types of changes on gene expression in spatial samples is an important task.

Clarifying the contribution of each source is essential for understanding how spatial regulation and cellular identity shape tissue architecture. For example, during embryonic development, positional information precedes cell identity: gradients of morphogens guide gene expression, differentiation, and tissue patterning before distinct cell types have formed. Methods such as DistMap [4] use spatial references to localize scRNA-seq cells in developing embryos. In tumor microenvironments, spatial organization reflects interactions between malignant and immune cells, often forming structured niches. Methods like Starfysh [7], stKeep [28], and SpaTopic [8] have uncovered spatial domains using expression patterns and graph-based decompositions.

In this work, we introduce scalable multi-group nonnegative spatial factorization (smNSF), a fast, flexible, and interpretable factorization method for spatial transcriptomics. smNSF integrates both spatial coordinates and known or inferred cell-type labels into a unified framework using multi-group Gaussian processes (MGGPs) [29]. Unlike standard GPs, MGGPs enable the modeling of structured variation across distinct groups (e.g., cell types) while sharing statistical strength across the spatial domain and across the group structure. smNSF extends the NSF framework [18] by replacing the GP priors with multi-group GP priors over spatial factors, building on the general MGGP framework of [29]. This modification allows smNSF to model cell-type specific variation in gene expression across space within the interpretable and sparse structure of nonnegative factorization. To make this concrete, we construct a new non-separable Matérn family of MGGP kernels in which group distance modulates both the spatial amplitude and the effective lengthscale; the kernel is derived from first principles via the spectral route (Bochner’s theorem) and proven positive-definite by Schoenberg’s theorem (Supplement §3). We develop a variational inference framework for MGGPs — including a new approximation we call *locally conditioned Gaussian processes* (LCGP) — to enable stable, scalable training on datasets with hundreds of thousands of cells. We demonstrate that smNSF recovers interpretable spatial factors across diverse ST platforms and enables cell-type conditional posterior predictions that distinguish spatial patterns driven by cell-type heterogeneity from those driven by spatial gradients of gene expression.

## 2 Materials and methods

### 2.1 Nonnegative spatial factorization

MEFISTO uses Gaussian processes as spatially-aware priors within factor analysis (FA) [16]. However, because FA is real-valued and not restricted to nonnegative values, the resulting MEFISTO patterns are often hard to interpret and not sparse [18].

Nonnegative spatial factorization (NSF) is a method that expands probabilistic nonnegative matrix factorization (PNMF), a probabilistic version of NMF, by using a Gaussian process as the prior instead of a univariate Gaussian prior. Restricting weights and factors to nonnegative values leads to a naturally sparse, parts-based decomposition. In NSF, this restriction provides interpretable nonnegative factors that capture small groups of genes that are co-expressed in a spatially smooth way [18].

The NSF model is as follows:

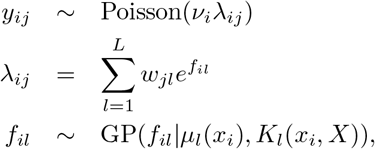

where:

- *v*_*i*_ is a per-cell normalization constant (e.g., total counts),
- *w*_*jl*_ are nonnegative gene loadings, enforced via softplus transformation,
- *f*_*il*_ are spatial latent factors drawn from a GP prior.

Here, the data consists of a cell-by-gene matrix of raw transcript counts *Y* ϵ ℝ^*N ×J*^ and spatial cell coordinates *X* ϵ ℝ^*N ×D*^ for each cell. Here, we consider 2D spatial coordinates (i.e., *D* = 2).

### 2.2 Multi-group Gaussian processes

Multi-group Gaussian processes (MGGPs) provide a family of positive-definite covariance kernel functions to perform Gaussian process regression on multi-group data [29]. The multidimensional inputs, *X* ϵ ℝ^*N ×D*^, also contain group labels, *C* = {*c*_1_, *c*_2_, …, *c*_*N*_}, and outcomes *Y* ϵ ℝ^*N*^ :

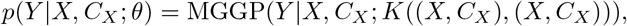

where the kernel takes both the input and group label:

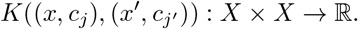

Various reductive special cases of MGGPs exist. The simplest case is separate GPs (SGP), where 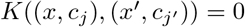, equivalent to training a separate GP per group [29]. Separate GPs per group do not allow information to be shared across groups, and SGP struggles when groups are imbalanced in size. In genomics, the SGP approach is not useful as it means training a separate NSF model per cell type, resulting in zero shared factors across cell types. The union GP (UGP) is equivalent to setting 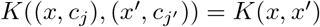, meaning group labels are ignored, which is the current baseline of the NSF model. Separable MGGPs use the covariance function as the Kronecker product of two kernels: 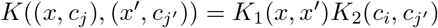.

However, these covariance functions can lead to “ridges” and discontinuities [29, 30]. For this work, we use the inseparable Matérn-3/2 MGGP kernel constructed in §2.3 (full derivation in Supplement §3).

### 2.3 Scalable multi-group nonnegative spatial factorization (smNSF)

Scalable multi-group nonnegative spatial factorization (smNSF) extends the nonnegative spatial factorization (NSF) framework [18] to incorporate both spatial coordinates and discrete group labels into a unified generative model. Here, we use cell-type labels as the group labels, allowing us to recover factors that distinguish structure in each cell type but with patterns shared across all cell types. smNSF recovers spatially structured latent factors using multi-group Gaussian processes (MGGPs) [29], enabling both (i) smooth signal sharing across similar groups and (ii) structured differences between groups.

Relative to NSF (§2.1), the input includes spatial coordinates *X*, but also a vector of group labels *C* = {*c*_1_, …, *c*_*N*_}; the MGGP prior now conditions on the pair (*x*_*i*_, *c*_*i*_). Using the MGGP, dimension reduction operates on the gene-by-count matrix, while taking into account spatial coordinates and cell-type labels (Fig. 1).

**Fig 1.**
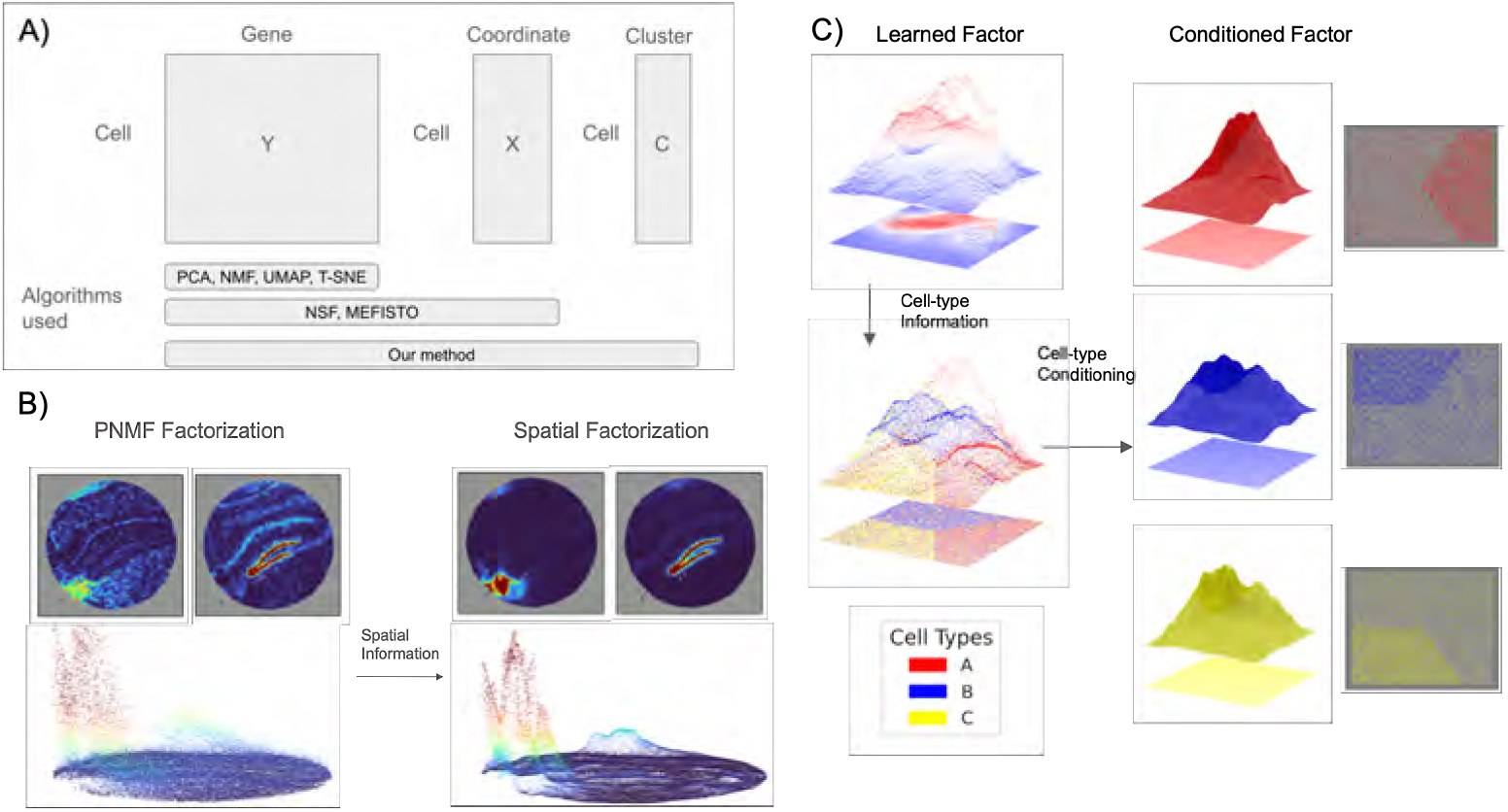
Overview of smNSF and cell-type conditional spatial factorization. A) Comparison of dimension reduction methods by their input data. Classical methods (PCA, NMF, UMAP, T-SNE) operate on the cell-by-gene matrix *Y* alone; NSF and MEFISTO additionally incorporate spatial coordinates *X*; smNSF further incorporates cell-group labels *C* (e.g., cell type, tissue region, or disease status). B) Incorporating spatial information transforms an independent cell signal into a smooth function across space: we learn a Gaussian process decomposition in which each factor is a nonnegative, continuous spatial function. The Matérn-3/2 kernel enables the model to capture sharp spatial signals without oversmoothing or compromising spatial correlation across the tissue. C) A single learned spatial factor can be conditioned on individual cell types; by incorporating cell-type information via our multi-group Gaussian process, we can compute the posterior conditioned on a given cell type, revealing how the decomposition of the tissue region would look if it were composed of that cell type alone.

#### smNSF model

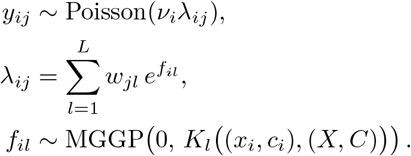

Here, *v*_*i*_ and the nonnegative loadings *w*_*jl*_ are as in §2.1; the novelty is that the latent factors *f*_*il*_ are drawn from an MGGP prior (S1 Fig) that explicitly accounts for the group label *c*_*i*_.

#### smNSF kernel

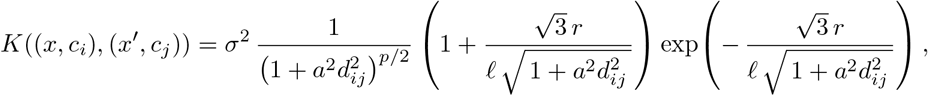

where *d*_*ij*_ encodes *group (cell-type) distances* (e.g., *d*_*ij*_=0 if *i*=*j, d*_*ij*_=1 otherwise), *σ* is the marginal scale, *a* controls the strength of difference between groups (cell types), *l* is the spatial inverse length-scale and *p* is the dimensionality of the data. Throughout this work, we use the Matérn-3/2 variant of the MGGP kernel, which allows controlled roughness at group boundaries while maintaining smoothness within groups. The closed-form expression for the Matérn-3/2 MGGP kernel is derived in Supplement §3.

### 2.4 MGGP inference

#### 2.4.1 Stochastic variational multi-group Gaussian processes

Exact inference under the NSF model is intractable: Gaussian process priors scale cubically with the number of observations, and the Poisson likelihood is non-conjugate, ruling out closed-form posteriors. We therefore adapt stochastic variational Gaussian processes (SVGP) [18, 31–34], which introduce a small set of *M* representative spatial locations called *inducing points* 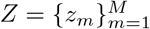 to reduce the computational burden.

Then, SVGP performs variational inference over the function values at those locations.

To extend SVGP to MGGPs, each inducing point also carries a cell-type label *c*_*m*_,forming *inducing groups* 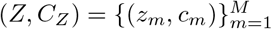 that mirror the structure of the observed group inputs; we sample *C*_*Z*_ from the empirical group distribution to ensure adequate coverage of every cell type. With this construction, we define a variational distribution *q*(*U*) over the inducing function values. The resulting evidence lower bound (ELBO) decomposes into a Poisson likelihood term, approximated by Monte Carlo and a closed-form KL divergence to the inducing prior. The full derivation is in Supplement §1 and §2. The Poisson expected log-likelihood expansion and multiplicative-update training methods are described in Supplement §11.

#### 2.4.2 Locally conditioned Gaussian processes (LCGP)

The cubic cost of inverting *K*_*ZZ*_ restricts standard SVGPs to *M* ≈ 3,000 inducing points — far below the actual cell counts in modern ST datasets. To capture both means and scales while preserving scalability, we derive a new variational approximation that maintains local covariance structure (full derivation in Supplement §5). We approximate the full-rank variational covariance via *local* neighborhood factors using an embedding *L*_*u*_ ϵ ℝ^*M ×K*^. For each neighborhood *n*(*i*):

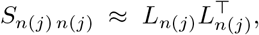

and we adopt the conditional variational structure:

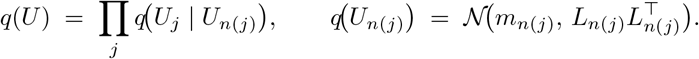

This formulation uses a full local covariance rather than independent variances, preserving cross-cell and cross-group correlations within each neighborhood (Supplement §8); the rank of each local covariance matrix is chosen by the spectral-decay criterion described in Supplement §6, giving a compact but expressive representation. Because the same conditional factorization is applied to *q*(*U*), the KL divergence decomposes into a sum of closed-form terms over neighborhoods (Supplement §5). Finally, neighbors are selected via kernel-conditioned sampling rather than Euclidean distance, producing neighborhood sets whose spatial extent remains stable across varying data densities (Supplement §7).

The resulting approach, which we term *locally conditioned Gaussian processes* (LCGPs), enables setting *M* = *N* (inducing points equal to total data points) without the (*OM* ^3^) bottleneck, while accurately capturing both the means and scales of spatial factors. We demonstrate the *M* = *N* regime on the Slide-seqV2 hippocampus (41,783 spots; Fig. 2); the same inference machinery scales to larger datasets, since the expensive local linear algebra depends on the neighborhood size *K* rather than requiring an *M* × *M* factorization, so per-step cost scales linearly rather than cubically in *M* when *K* is fixed. This is critical for smNSF: the scales encode biologically meaningful uncertainty and cell-type specific variance that would be lost under diagonal approximations.

**Fig 2.**
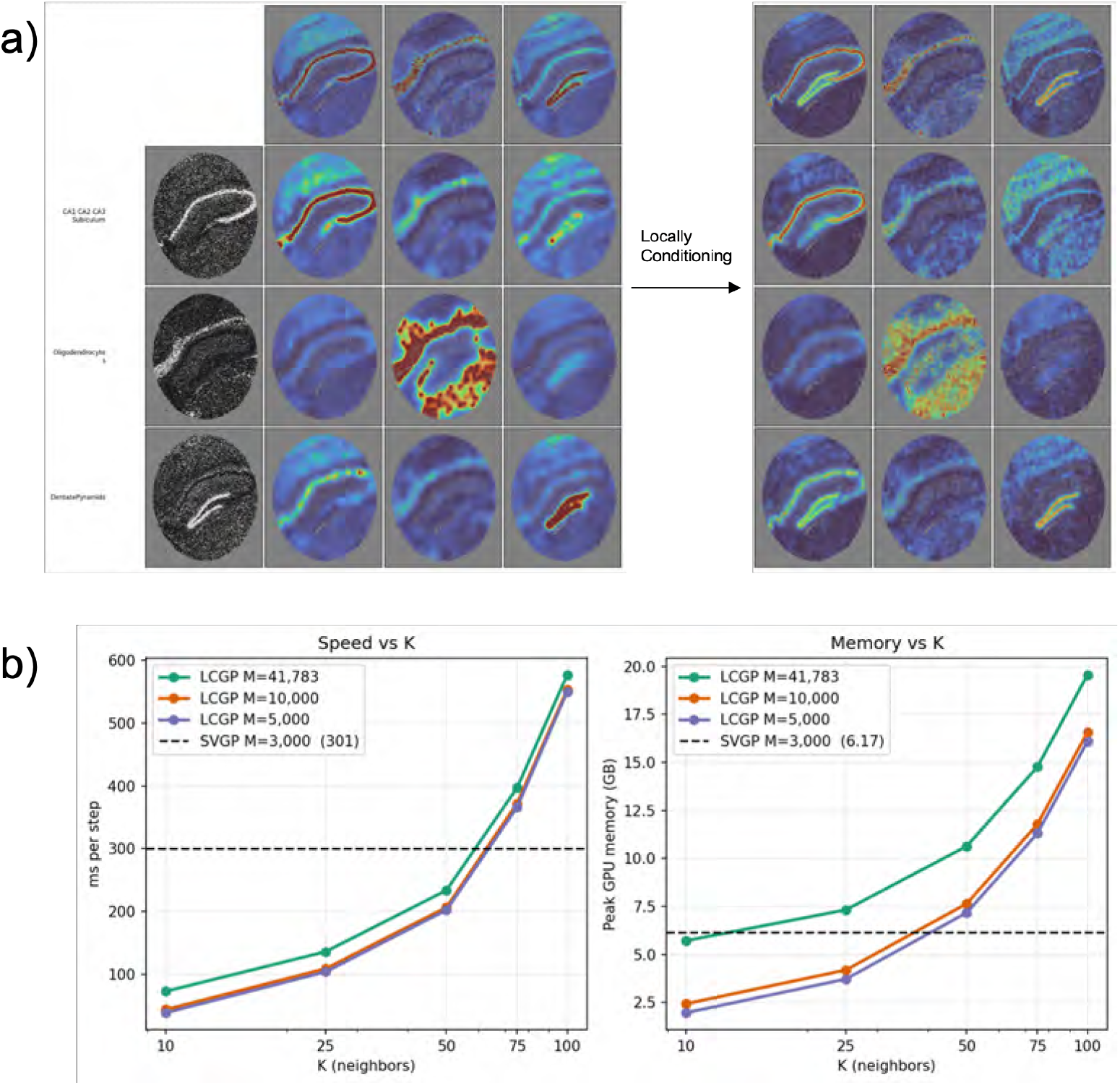
Comparison of MGGP-SVGP and MGGP-LCGP: cell-type conditional posteriors in mouse hippocampus and scalability with dataset size. **a)** Left: spatial factors learned by smNSF using an MGGP-SVGP prior (*M* = 3,000 global inducing points). Although conditioning on cell types is possible via the multi-group Gaussian process, the 3,000 inducing points must represent the full tissue across all annotated groups (here, 14 groups). The fraction of inducing points available to represent any single cell type is therefore small, yielding a weak and spatially underresolved conditional signal. Right: corresponding cell-type conditional posteriors from MGGP-LCGP, which uses all 41,783 cells as inducing points. Each cell’s posterior factor value is computed by conditioning on *K*=50 local neighbors (see Supplement), providing a dense approximation to the otherwise computationally intractable true posterior. Comparing the two models, LCGP recovers a qualitatively similar but substantially richer conditional signal, closer to the true posterior. Conditioning reveals distinct modes of cell-type specific spatial organization. Some cell types deplete the learned factor signal, others isolate it into a sharper spatial pattern, and others uncover spatial structure that is only apparent upon conditioning—spatial signals that are diluted or obscured when all cell types are pooled together, but emerge as distinct patterns within a single cell-type specific conditional posterior. **b)** Scalability comparison of LCGP versus SVGP as dataset size *N* increases. SVGP inference is restricted to a fixed number of inducing points (*M* = 3,000), limiting expressiveness regardless of data size. LCGP sets *M* = *N* in this benchmark, using all 41,783 Slide-seqV2 spots as inducing points, and replaces the cubic inducing-point bottleneck with local *K*-neighbor computations.

### 2.5 Cell-type conditional posteriors from smNSF

Now that we have the variational construction, we can compute a posterior GP that depends only on the corresponding inputs (*X*_*i*_, *C*_*i*_) [34].

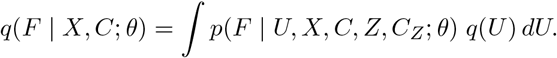

Critically, unlike a standard SVGP, the MGGP form permits predictions not only at new spatial locations but also under *new or altered group labels*. Enabling us to ask questions such as “what if the entire tissue were cell type A?”, turning the model into a practical, cell-type specific framework rather than stopping at a smooth factorization of heterogeneous cells.

To quantify the dependence of spatial factors on cell-type labels, we perform *in silico* perturbation experiments. Given a trained model, we hold spatial coordinates *X* fixed, and we replace all labels *C*_*X*_ with a new configuration 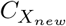, e.g., we set all labels to a single cell-type.

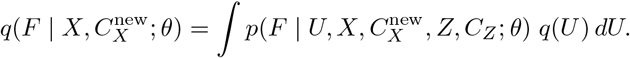

This allows us to generate posterior maps of spatial factors under controlled label perturbations, revealing three qualitatively distinct specificity classes for each factor with respect to group labels (Fig. 3):

**Fig 3.**
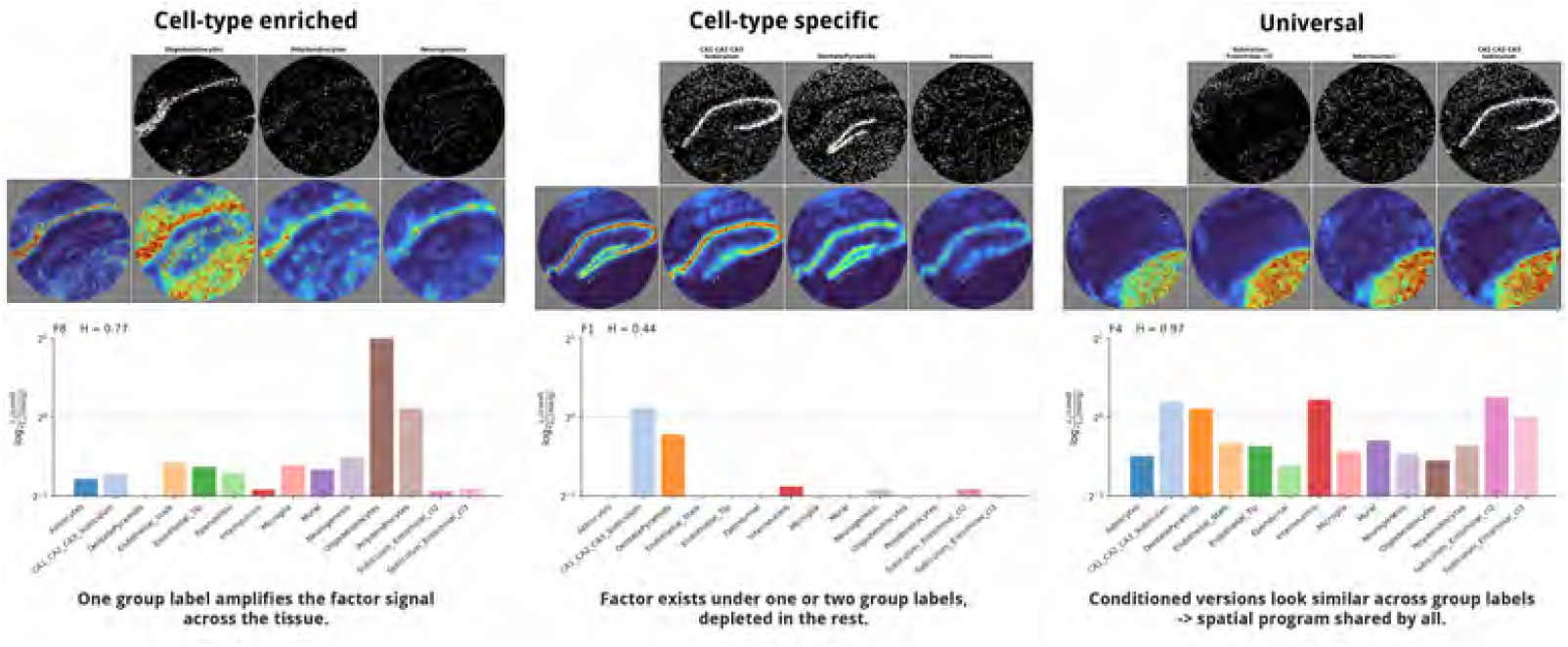
Three specificity classes of cell-type conditioning. Representative Slide-seqV2 hippocampus factors illustrating each class, ordered left to right as introduced in the text: *cell-type enriched* (Factor 8), *cell-type specific* (Factor 1), and *universal* (Factor 4). For each factor, the upper rows show the marginal factor alongside posteriors conditioned on individual cell-type labels; the lower bar plot shows the conditional-to-marginal *L*_1_-ratio across all groups. A cell-type enriched factor is amplified by a single group, a cell-type specific factor persists under one or two groups while depleting under the rest, and a universal factor is nearly invariant to conditioning.

- *cell-type enriched* — a spatial pattern that emerges from conditioning a factor on a given cell type but is absent or attenuated in the marginal factor considering all cell types. These patterns are only visible using the MGGP prior, and they reflect the patterns considering a hypothetical abundance of the conditioned cell type that are otherwise masked by heterogeneous and sparse cell-type representation.
- *cell-type specific* — a spatial pattern that is largely unchanged for a given cell type but is depleted (approaches zero) when conditioning on other groups, isolating the pattern to a single cell type.
- *universal* — a spatial pattern that is nearly invariant to group conditioning, representing spatial structure shared broadly across cell types.

We quantify these classes from each factor’s conditional profile across groups, combining the *L*_1_*-ratio* of conditional to marginal posterior with cross-label specificity. The *L*_1_-ratio is ∥*F*_*l*_ |*c*∥ _1_*/* ∥*F*_*l*_∥ _1_, where *F*_*l*_ is the marginal posterior of factor *l* and *F*_*l*_ *c* is its posterior conditioned on group label *c*; values *>* 1 indicate enrichment under *c*, values *<* 1 indicate depletion, and values ≈ 1 indicate invariance. The three classes use this metric differently: *cell-type enriched* flags a (factor, group) pair when its *L*_1_-ratio exceeds 1.5× (the threshold used throughout Results); *cell-type specific* requires the dominant group’s *L*_1_-ratio to stay near 1 while other groups deplete to near zero (i.e., cross-label depletion, not the single-group ratio alone); *universal* requires all conditional *L*_1_-ratios close to 1, equivalently near-maximal Shannon entropy across the group axis.

### 2.6 Implementation details

Unlike the original NSF, which requires an NMF warm-start to initialize its factors [18], smNSF initializes both the nonnegative loading matrix *W* and the GP variational parameters (*m, L*) randomly. We find that training reliably converges from random initialization under MGGP-SVGP and MGGP-LCGP priors (Fig. 2). Recovered factors are broadly consistent across random seeds; as expected given the convergence to a local minimum inherent to NMF, factor identity is not strictly seed-invariant, although the sparsity of the parts-based representation inherent in NMF encourages identifiability up to label switching. We apply stochastic variational inference with mini-batches of 6,000 cells. For LCGP, we set the neighborhood size *K* = 50 across all experiments. Spatial coordinates are standardized to unit scale per axis. All experiments used PyTorch on an NVIDIA A30 GPU (24 GB). Code is available at https://github.com/luisdiaz1997/GPzoo (MGGP and LCGP inference), https://github.com/luisdiaz1997/Probabilistic-NMF (nonnegative and spatial nonnegative factorizations), and https://github.com/luisdiaz1997/Spatial-Factorization (full smNSF pipeline).

### 2.7 Datasets and preprocessing

We applied smNSF to seven spatial transcriptomics datasets spanning different technologies, scales, and biological contexts.

#### Slide-seqV2 mouse hippocampus

Mouse hippocampus tissue with 41,783 spots that are composed of multiple cells, 17,702 genes, and 14 annotated groups (10 cell types and 4 hippocampal subfield regions) [35]; the dataset used in the original NSF publication [18]. Used for computational benchmarking (LCGP vs. standard SVGP, enabled by the moderate dataset size) and for group-conditional analysis of hippocampal cell types and subfield regions.

#### HuColonCa-FFPE MERFISH human colorectal cancer

The HuColonCa-FFPE dataset (Vizgen) [22, 36] spans approximately 1.1 million cells across multiple patients; we analyze Patient 1, which contains 844,468 cells before subsampling and 137,693 cells after 6 × subsampling, with 211 genes and 13 cell-type groups collapsed from the fine-grained annotation.

#### osmFISH mouse somatosensory cortex

osmFISH data from mouse primary somatosensory cortex with 4,727 cells, 33 genes, and 28 distinct cell-type labels [37]. Data accessed via Squidpy tutorials [38].

#### Human liver (healthy vs. fibrotic) MERFISH

FFPE MERFISH liver samples, 317 genes, with three healthy donors (AM042, AM048, AM061; 27,722–33,878 cells each) and three fibrotic donors (AM031, AM062, AM072; 65,668–82,178 cells each); three healthy donors are additionally re-fit under the joint healthy–fibrotic label schema for cross-condition comparison, yielding nine per-sample fits in total. Cell-type annotations cover 15 distinct types across the cohort (5–8 per sample, varying with tissue state). Used to demonstrate disease-specific spatial remodeling of factors and cell-type conditional posterior interpretations.

#### MERFISH mouse hypothalamus (3D)

Three-dimensional MERFISH data from mouse hypothalamic preoptic region with 71,939 cells across multiple z-slices and 161 genes [39]. The original dataset contains 16 annotated cell classes; after filtering rare cell types below 1% frequency, 11 cell-type groups were used for training. Demonstrates smNSF extension to volumetric spatial data. Data accessed via Squidpy tutorials [38].

#### 10x Visium adult mouse brain

Coronal Section 1 from the 10x Genomics Adult Mouse Brain dataset (2,688 spots, 16,944 genes, 15 brain region groups) accessed via Squidpy [38]. Sequencing-based spatial platform; each 55 μm spot mixes multiple cell types.

#### DLPFC (SDMBench, 10x Visium)

Human dorsolateral prefrontal cortex across 12 tissue sections [40], 3,611–4,381 spots per section, 33,538 genes, with 7 annotated cortical layers per section. As this dataset contains layer annotations rather than cell-type labels, we applied *K*-means clustering to smNSF factors and compared against published benchmarks from SDMBench [41].

Across all datasets we applied a common base pipeline: cells or spots with missing spatial coordinates, expression values, or group labels were removed, and any annotation group comprising fewer than 1% of the cells in a sample was dropped, with the surviving group labels re-encoded contiguously. For the two datasets ingested from raw transcriptome-wide counts—the Slide-seqV2 hippocampus and the 10x Visium mouse brain—we additionally applied standard single-cell quality control: cells or spots with fewer than 100 total counts were removed, genes detected in fewer than 10 cells were removed, and mitochondrial genes were excluded (the Slide-seqV2 puck additionally excluded beads with more than 20% mitochondrial counts). The targeted imaging panels (MERFISH colorectal cancer, osmFISH cortex, MERFISH hypothalamus, and MERFISH liver) carry small curated gene sets and were not subjected to count-based cell or gene filtering; the colorectal cancer sample was further subsampled to every sixth cell (844,468 → 137,693 cells) to reduce computational cost. The DLPFC sections were used as preprocessed by SDMBench [41], with the same group-fraction filter applied. All cell, spot, and gene counts reported above are post-quality-control. Spatial coordinates were standardized to unit scale per axis, and for LCGP inference we used mini-batches of 6,000 cells and *K* = 50 in all datasets.

### 2.8 Computational benchmark methodology for smNSF

To evaluate the computational efficiency of the LCGP approximation, we compared against standard SVGP on the Slide-seqV2 hippocampus dataset (41,783 barcodes). For LCGP, we set inducing points *M* = *N* (all data points serve as inducing points) and varied the neighborhood size *K* ϵ {10, 25, 50, 75}. For SVGP, we used *M* = 3,000 inducing points, representing the practical upper limit under equivalent GPU memory constraints (NVIDIA A30, 24 GB).

We measured: (i) wall-clock time per optimization step, (ii) peak GPU memory usage, and (iii) convergence behavior over 10,000 training steps. All experiments used identical hyperparameters (learning rate, batch size of 6,000 cells, 10 latent factors) and were repeated three times to account for variability.

### 2.9 Spatial domain benchmark methodology

To compare smNSF against spatial domain identification methods, we evaluated on the DLPFC dataset [40] using published benchmarks from SDMBench [41]. smNSF takes the Maynard et al. cortical layer annotations as its MGGP group labels at training time, and is a factorization rather than a clustering method. We nonetheless include this benchmark because SDMBench is the standard comparison axis for spatial transcriptomics methods on DLPFC. Concretely, we apply *K*-means (*K* = 7, matching the seven annotated layers) to the learned spatial factors and score the resulting cluster assignments against the same Maynard et al. layer annotations using the SDMBench metrics below. “Clustering accuracy” throughout this section refers to that cluster-vs-layer agreement.

We report the following metrics following the SDMBench protocol:

- **Adjusted rand index (ARI)**: Clustering accuracy adjusted for chance.
- **Normalized mutual information (NMI)**: Mutual information between predicted and true labels, normalized to [0, 1].
- **Homogeneity (HOM)**: Whether clusters contain only members of a single class.
- **Completeness (COM)**: Whether all members of a class are assigned to the same cluster.
- **CHAOS**: Spatial chaos score measuring cluster fragmentation.
- **Percentage of abnormal spots (PAS)**: Lower percentages indicate better spatial continuity.
- **Average silhouette width (ASW)**: Average silhouette width of the factor embedding.
- **Moran’s I**: Spatial autocorrelation of factors.

We compared results from smNSF against 14 methods benchmarked in SDMBench, including SpaGCN, BayesSpace [42], STAGATE [43], SpaceFlow, BASS, and others, using their published results on identical DLPFC sections.

### 2.10 Baseline methods

We benchmarked smNSF against a range of existing methods for spatial decomposition, domain discovery, and spatial deconvolution:

- **PCA / FA**: non-spatial linear factor models [11],
- **NMF / PNMF**: nonnegative factorization without spatial priors [12],
- **NSF**: nonnegative spatial factorization using exponentiated GP priors [18].

This broad landscape of spatial transcriptomics methods vary in fundamental ways; direct quantitative comparisons were performed against the three methods above (Table 1).

**Table 1.**
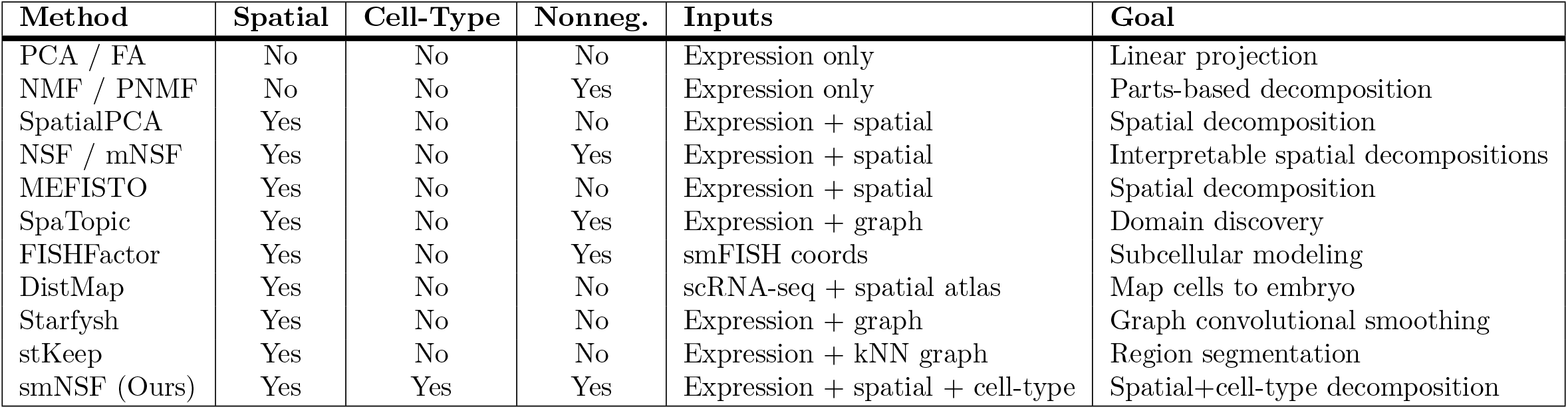
Comparison of spatial transcriptomics methods. Methods differ in whether they incorporate spatial information, account for cell types, use nonnegativity constraints, and in their modeling goals.

Three spatially aware methods that we discuss but are excluded from the head-to-head benchmark warrant brief justification. **MEFISTO** [16] uses a GP-over-spatial-coordinates prior over real-valued factors. NSF dominates MEFISTO on spatial-transcriptomics benchmarks [18], so the smNSF vs. NSF comparison below is the binding bar for spatially aware nonnegative factorization in this class.

**SpatialPCA** [17] shares the spatial-smoothing goal but lacks nonnegativity and cell-type conditioning. We therefore use NSF as the more relevant nonnegative spatial-factorization baseline.

**NSFH** [18] is the hybrid spatial/nonspatial variant of NSF. We neither incorporate this split into smNSF nor benchmark against NSFH, because the spatial/nonspatial decomposition addresses a different problem from the cell-type conditioned spatial factorization studied here.

## 3 Results

### Overview

We evaluate smNSF on seven spatial transcriptomics datasets spanning MERFISH, Slide-seq, osmFISH, and 10x Visium technologies: mouse hippocampus (Slide-seqV2, 41,783 spots, 14 groups), human colorectal cancer (MERFISH, 137,693 cells, 13 cell-type groups), mouse somatosensory cortex (osmFISH, 4,727 cells, 28 cell-type groups), human liver (MERFISH, 9 samples, 27k–82k cells each, 15 cell-type groups across the cohort), mouse hypothalamic preoptic region (MERFISH, 71,939 cells, 11 cell-type groups), mouse brain (10x Visium, 2,688 spots, 15 brain region groups), and human DLPFC (12 Visium sections, ~ 4,200 spots each, 7 cortical layer groups). We assess (i) scalability and computational efficiency of LCGP versus standard SVGP, (ii) factor interpretability through sparsity and spatial coherence, and (iii) cell-type specificity through conditional posterior analysis. We benchmark clustering accuracy on DLPFC against NSF, PCA, NMF, and 14 methods from SDMBench [41].

### LCGP enables scalable inference with accurate variance estimation in smNSF

A key limitation of NSF is the (*OM* ^3^) cost of standard SVGP inference, which restricts inducing points to *M* ≈ 3,000 under typical GPU memory constraints. For large ST datasets with *>*100,000 cells, this severely limits model expressiveness.

We compared LCGP against SVGP on the Slide-seqV2 hippocampus dataset [35](~41,783 spots, 14 groups) to evaluate computational trade-offs (Fig. 2b). Per training step with *L* factors and batch size *B*, SVGP scales as *O* (*LBM* ^2^) while LCGP scales as *O* (*LBK*^3^), reflecting the cost of inverting *K*×*K* matrices in the local approximation (full derivation in Supplement §5). Critically, LCGP’s cost depends on *K*, not *M*, allowing *M* =*N* without the cubic growth in *M* that limits standard SVGPs.

For LCGP, we set *M* = *N* (inducing points equal to data points) and varied the neighborhood size *K* ϵ{10, 25, 50, 75}. For SVGP, we used *M* = 3,000 inducing points—the practical limit under equivalent memory constraints (NVIDIA A30, 24 GB). At *K* = 10, LCGP was 4× faster than SVGP (73 vs. 301 ms/step) with lower peak memory (5.72 vs. 6.17 GB). At *K* = 50, per-step time remained faster than SVGP (234 vs. 301 ms) while memory grew to 10.64 GB. At *K* = 75, both time (397 ms) and memory (14.77 GB) exceeded SVGP as the full *M* =*N* inducing set saturates GPU capacity. We select *K* = 50 across all experiments to balance speed and expressiveness.

Beyond computational efficiency, LCGP also produces substantially richer cell-type conditional posteriors than SVGP (Fig. 2a). Conditioning on individual cell types reveals distinct modes of spatial organization: some cell types deplete the learned factor signal (e.g., oligodendrocytes in factors 2 and 4), others isolate it into a sharper spatial pattern (e.g., dentate pyramidal neurons in factor 4), and others uncover spatial structure that is only apparent upon conditioning—spatial signals that are diluted or obscured when all cell types are pooled together, but emerge as distinct patterns within a single cell-type conditional posterior (e.g., CA1/CA2/CA3 subiculum neurons and oligodendrocytes in Fig. 2a, where each cell type’s conditional posterior reveals a sharp tissue program in a factor whose marginal map is near-uniform). Although ground-truth conditional posteriors are unavailable for these data, the agreement between SVGP and LCGP—which use fundamentally different variational approximations—and across random initializations suggests these patterns reflect genuine biological structure rather than inference artifacts.

### Gain from finite group-distance coupling

The previous subsection establishes that MGGP-LCGP produces interpretable cell-type conditional posteriors. A natural follow-up question is whether the multi-group component of the MGGP framework is contributing anything beyond what an ensemble of independent per-cell-type Gaussian processes would deliver. The non-separable Matérn 3/2 MGGP kernel of §2.2 has a single group-distance parameter *a* governing cross-group correlation: at *a* = 0 the kernel reduces to a standard single-group Matérn (all cells share one GP regardless of cell type), and at *a*→ ∞ the cross-group covariance collapses to zero and each cell type is fit by a separate GP, with no shared signal across groups. The regime 0 *< a <* ∞ is where the multi-group machinery does useful work—each cell type’s conditional posterior borrows strength from cells in correlated cell types without being forced to share a single global mean.

To isolate the contribution of this finite-*a* coupling we fit two MGGP-LCGP variants that differ only in the value of *a*. Everything else—kernel form, *K* = 50, *M* = *N*, probabilistic neighbor selection, loadings parameterization, training schedule, and the fixed cell-type distance matrix—is held fixed. The first variant pins *a* = 1, a moderate cross-group coupling intermediate between *a*→ 0 (which collapses the prior toward a single shared GP across all cell types) and *a*→ ∞ (which recovers independent per-cell-type GPs); the second variant pins *a* = 10^6^, large enough to drive the off-diagonal MGGP covariance below numerical precision and recover the separate-GPs limit in practice (Figure 4).

**Fig 4.**
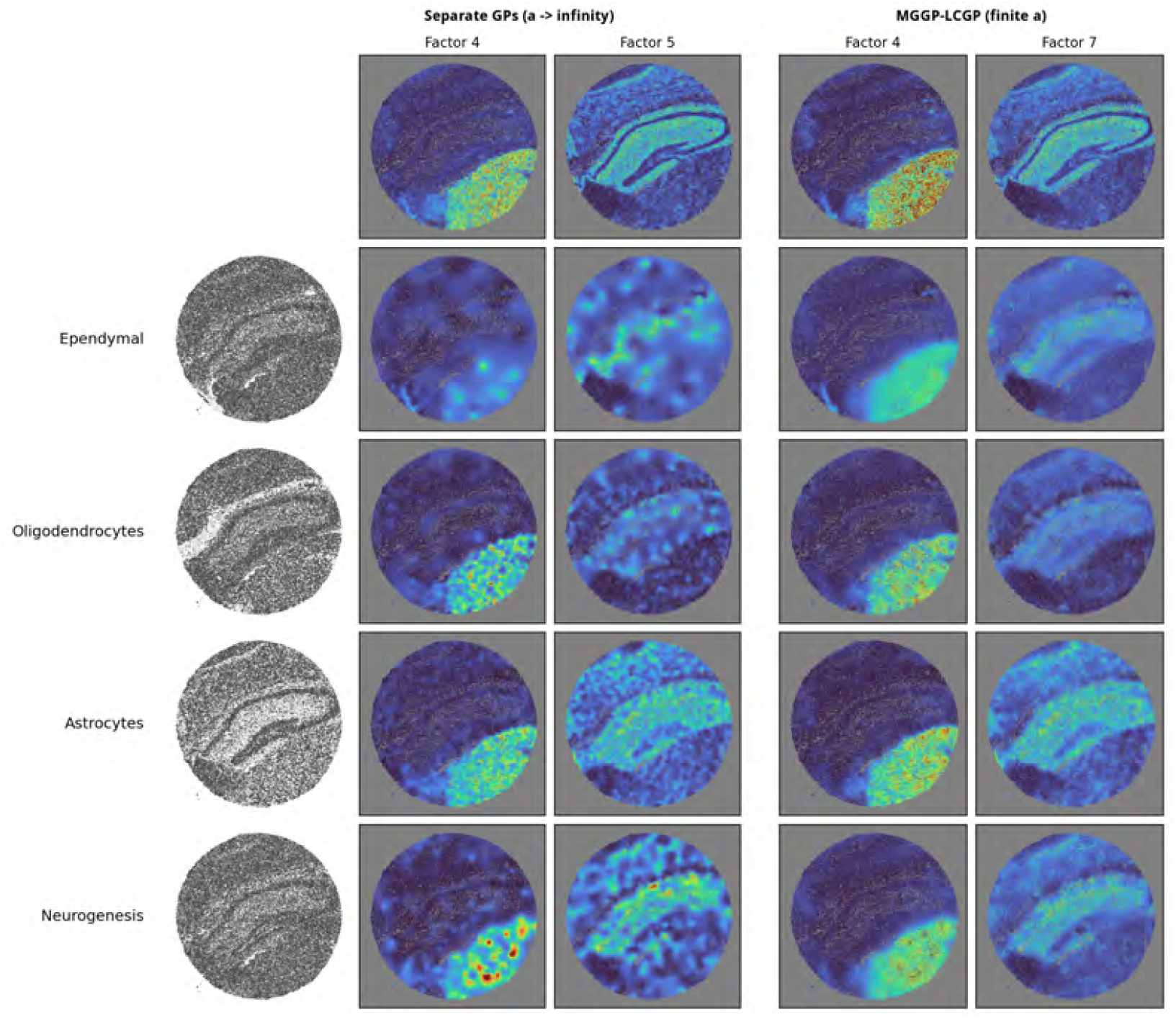
Gain from finite group-distance coupling on Slide-seqV2 mouse hippocampus. Both blocks are MGGP-LCGP fits with identical kernel form, *K* = 50, *M* = *N*, probabilistic neighbor selection, and the same fixed cell-type distance matrix; only the MGGP group-distance parameter *a* differs. Left: *a* → ∞ (specifically *a* = 10^6^), the separate-GPs limit. Right: *a* = 1, a moderate cross-group coupling where the multi-group kernel borrows strength across cell types. Each block shows the unconditional spatial factor map (top row) and four cell-type conditional posteriors below; the leftmost column of the left block shows which cells belong to each group. See main text for factor and cell-type selection rationale.

We selected the two universal factors on the finite-*a* side (factors whose Shannon entropy of normalized *L*_1_ specificity ratios is near uniform across cell types) and matched them to the *a*→∞ side by best Pearson correlation on the per-factor gene-loading vectors, so that the same biological gene program is being compared across blocks; exact factor indices are not preserved across fits because nonnegative matrix factorization is rotation- and permutation-invariant in the nonnegative cone. Cell-type rows are the four groups (ependymal, oligodendrocytes, astrocytes, neurogenesis) where the matched *a*→ ∞ factors are most depleted by the *L*_1_ specificity ratio.

The marginal posteriors in the top row are broadly similar across the two fits, as expected for factors with diffuse cell-type usage. The per-cell-type conditionals illustrate the role of cross-group coupling. Under large *a* the model is still LCGP—the probabilistic neighbor-selection scheme continues to weight candidates by kernel similarity—but the group-distance term drives those weights to favor same-group neighbors, so each cell type’s conditional posterior is in practice built with cells from its cell type alone. Cross-group covariance collapses to zero, and the model reduces to a collection of decoupled per-group GPs that share inducing locations but exchange no information at inference time.

This decoupling diminishes the conditional posterior of universal factors specifically.

A universal factor carries shared signal across many cell types; without a non-zero between-group covariance the model has no mechanism for one group’s data to inform another group’s conditional posterior on that factor. Each per-group GP must then recover the shared structure from its own observations alone, and where the per-group sample is small or noisy the conditional reconstruction degrades (Fig. 4). Under finite *a*, the same factor is anchored to a shared latent variable through non-zero off-diagonal group-kernel entries, and the conditional posteriors recover a coherent spatial pattern across the same groups.

### Comparison to baseline and published methods on DLPFC

To benchmark smNSF against established dimension reduction methods, we evaluated these methods on the DLPFC dataset [40], which contains twelve tissue sections with seven annotated cortical layers. We compared six model variants: PCA, PNMF (NMF without a spatial prior), SVGP (standard variational GP, equivalent to NSF), LCGP NSF (our scalable GP version of NSF), MGGP-SVGP, and MGGP-LCGP (smNSF). For all methods, we applied *k*-means clustering (*K* = 7) to the learned factors and evaluated against ground-truth layer annotations using metrics from SDMBench [41].

smNSF (MGGP-LCGP) achieved the highest clustering accuracy among our methods (ARI: 0.37, NMI: 0.51), substantially exceeding the standard spatial GP baselines (LCGP ARI: 0.22, NMI: 0.32; SVGP ARI: 0.33, NMI: 0.44) and non-spatial methods (PCA ARI: 0.15, NMI: 0.23; PNMF ARI: 0.30, NMI: 0.41; Fig. 5). The MGGP prior provided a 59% improvement in NMI over LCGP alone on these data, demonstrating that even region-level group structure aids factor recovery.

**Fig 5.**
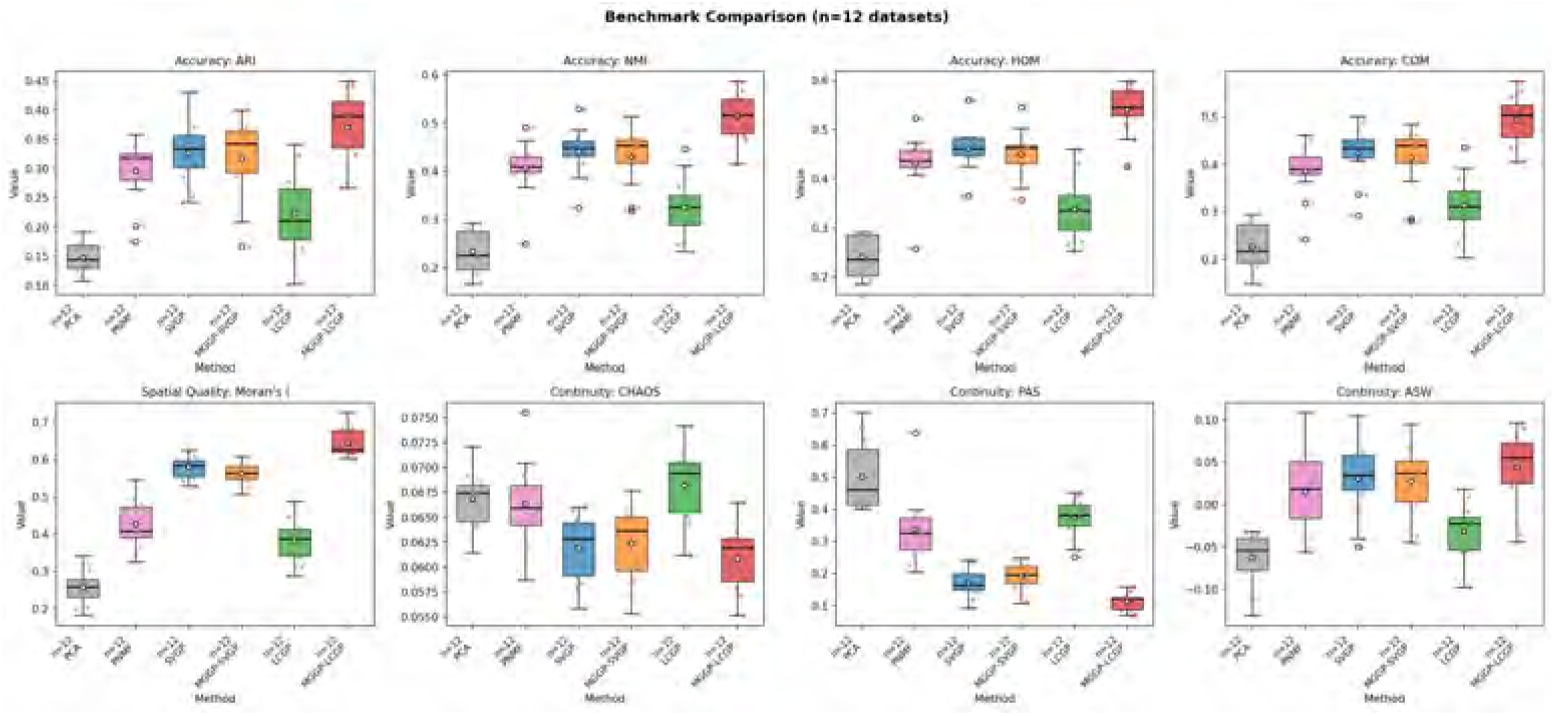
Benchmark comparison on DLPFC dataset. Box plots show performance across 12 tissue sections for PCA, NMF, LCGP (spatial GP without cell-type information), and smNSF (MGGP with group labels). **Top row**: Clustering accuracy metrics—adjusted Rand index (ARI), normalized mutual information (NMI), homogeneity (HOM), and completeness (COM). Higher values indicate better agreement with ground-truth layer annotations. **Middle row**: Spatial continuity metrics—CHAOS score, percentage of abnormal spots (PAS; lower is better), and average silhouette width (ASW). **Bottom row**: Factor spatial autocorrelation—Moran’s I. smNSF achieves the highest clustering accuracy, while both GP-based methods (LCGP, smNSF) show dramatically improved spatial continuity (low PAS) compared to non-spatial methods.

We further compared smNSF against 14 published spatial domain identification methods from SDMBench (Fig. 6). smNSF achieved the highest factor spatial autocorrelation of all 20 methods (Moran’s I = 0.645 vs. 0.385 for the best published method, SEDR), reflecting the MGGP prior’s strong spatial regularization (Fig. 7; Figs. S2–S13). On clustering accuracy, smNSF ranked 8th of 20 (NMI = 0.514), which is notable given that smNSF is designed to leverage cell-type labels as input rather than to predict spatial domains, and competitive with dedicated domain identification methods that leverage spatial neighborhood graphs (GraphST: 0.613, BASS: 0.609, BayesSpace: 0.601).

**Fig 6.**
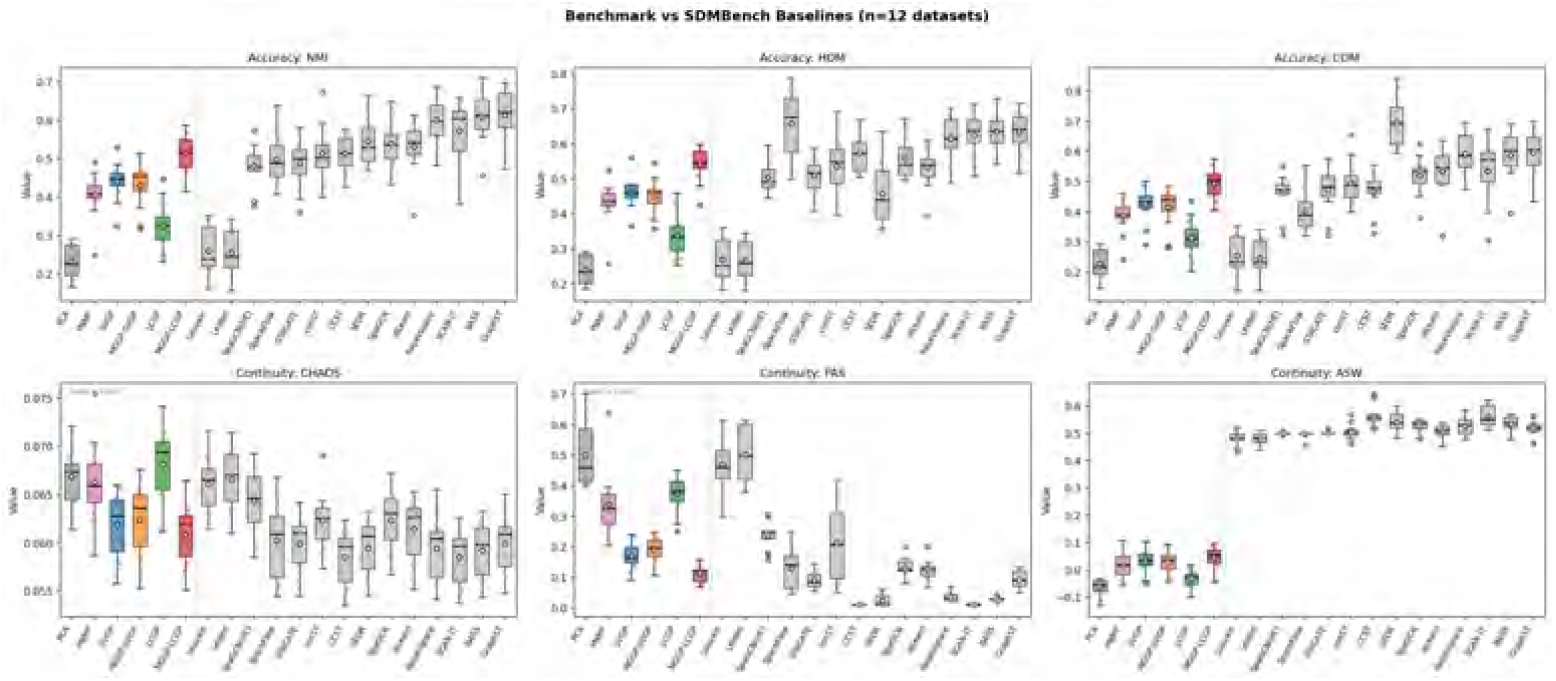
Comparison with SDMBench baselines on DLPFC dataset. Box plots show performance across 12 tissue sections for our six model variants (PCA, NMF, SVGP, LCGP, MGGP-SVGP, smNSF) alongside 14 published spatial domain identification methods from SDMBench [41]: BayesSpace, BASS, CCST, conST, GraphST, Leiden, Louvain, SCAN-IT, SEDR, SpaGCN, SpaGCN(HE), STAGATE, stLearn, SpaceFlow. SDMBench methods are sorted left-to-right by median NMI. Metrics shared with SDMBench: NMI (normalized mutual information), HOM (homogeneity), COM (completeness), CHAOS, PAS (percentage of abnormal spots; lower is better), and ASW (average silhouette width). smNSF achieves competitive clustering accuracy (NMI, HOM, COM) relative to top SDMBench methods, while both GP-based methods (LCGP, smNSF) show markedly better spatial continuity (low PAS, low CHAOS) than all 14 baselines.

**Fig 7.**
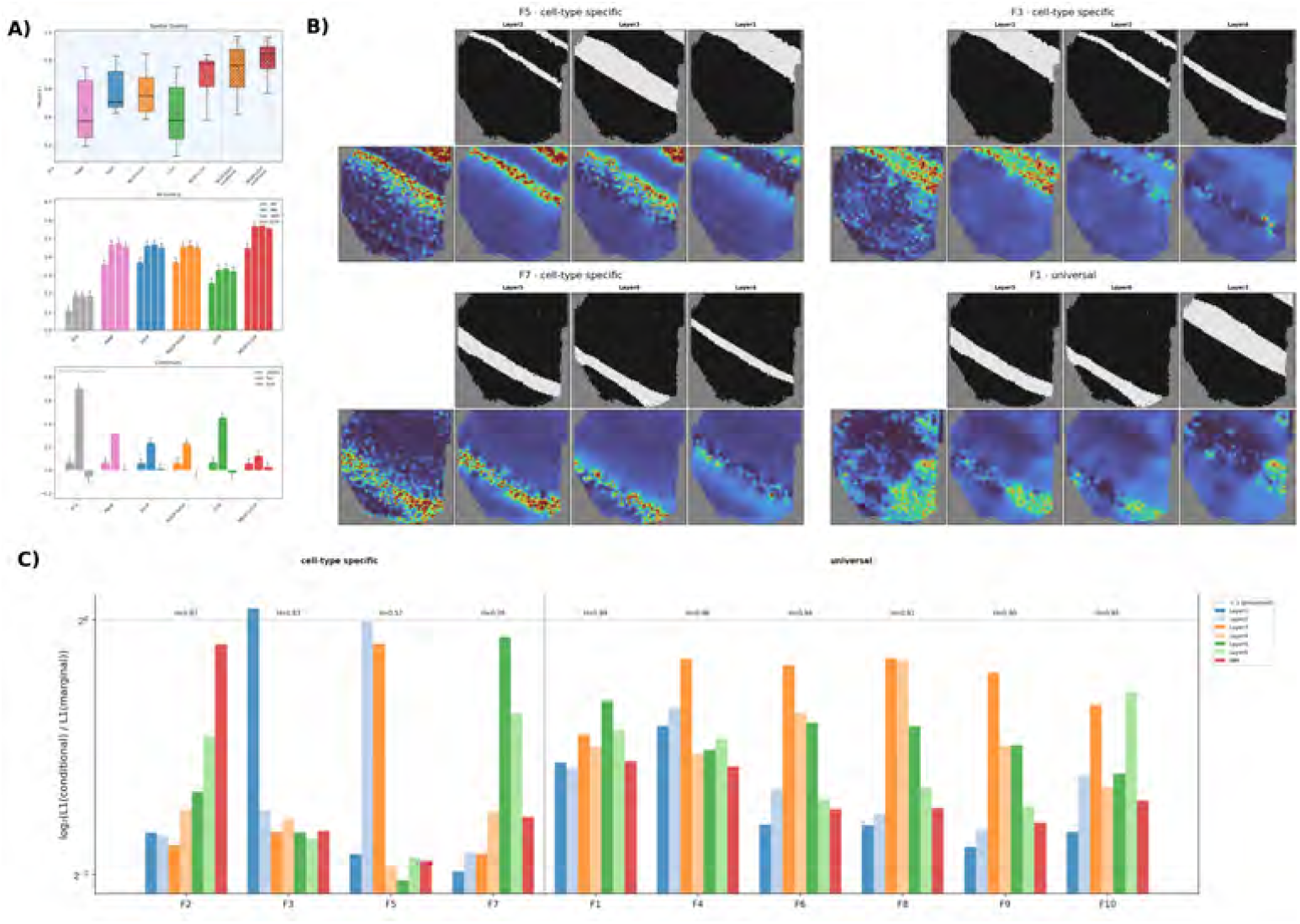
smNSF analysis of DLPFC slice 151507 (representative). **a)** Cross-method benchmark comparison on a single DLPFC slice. **b)** Curated layer-conditional posterior factors across specificity classes. Within each factor block: top row shows spatial domains of the selected layers (binary masks); bottom row shows the marginal factor (leftmost) followed by posteriors conditioned on the layer above each column. **c)** Factor specificity bar chart showing per-layer enrichment/depletion patterns. No cell-type enrichments exceed 1.5× across factors, consistent with cortical layers representing spatially segregated regions rather than intermingled cell types.

The DLPFC annotations represent cortical *layers* (spatial regions) rather than *cell types*. On these region labels, smNSF achieves competitive clustering accuracy and the highest spatial coherence among all benchmarked methods, but no layer exceeds the 1.5 × *L*_1_-ratio threshold defined in §2.5 (Fig. 7c). The next section summarizes the cell-type-vs-region pattern this exemplifies across the seven datasets in our analysis.

### Cell-type labels are enriched across regions; region groups are localized

The smNSF result reflects a structural property of the MGGP prior that generalizes across datasets. When smNSF conditions on *cell types*, which are scattered throughout the tissue, signal propagates across the full spatial domain, producing conditional posteriors that show enrichment of single cell types in specific factors (*L*_1_-ratio *>* 1.5×). When it conditions on *spatial regions*, which are locally confined, the prior collapses signal to that single location, suppressing enrichment everywhere else. We quantified this pattern by computing, for every group across all seven datasets, whether at least one factor reached an *L*_1_-ratio above 1.5× (Table 2).

**Table 2.** Cell-type groups enrich; region groups do not. Percentage of groups with at least one factor exceeding 1.5 × *L*_1_-ratio enrichment, stratified by group label type (cell type vs. region).

| Dataset | Cell types | % $>1.5\times$ | Regions | % $>1.5\times$ |
| --- | --- | --- | --- | --- |
| Slide-seq hippocampus | 10 | 50.0% | 4 | 0.0% |
| MERFISH colon | 13 | 69.2% | 0 | — |
| osmFISH cortex | 26 | 38.5% | 2 | 0.0% |
| MERFISH hypothalamus | 11 | 81.8% | 0 | — |
| 10x Visium brain | 0 | — | 15 | 0.0% |
| DLPFC (12 Visium slices) | 0 | — | 7 | 0.0% |
| MERFISH liver (9 samples) | 15 | 86.7% | 0 | — |
| **Total** | **75** | **61.3%** | **28** | **0.0%** |

Across 75 cell-type groups, 61% (46/75) showed enrichment above 1.5×. Across 28 region groups, 0 out of 28 reached 1.5 × —the strongest region enrichment across any dataset was 1.41× (CA1 subfield, Slide-seqV2 mouse hippocampus). This distinction holds within individual datasets that contain both group types: in the Slide-seq hippocampus, 5/10 cell types were enriched above 1.5×(microglia: 5.4×, oligodendrocytes: 3.2×, astrocytes: 2.6 ×) while 0/4 hippocampal subfields were enriched; in osmFISH, 10/26 cell types were enriched while 0/2 region groups were enriched. Datasets annotated exclusively with cell types showed consistently high enrichment rates (colon 69%, MERFISH hypothalamus 82%, liver 87%); datasets annotated exclusively with regions (10x Visium, DLPFC) showed no enrichment. This structural distinction—scattered cell types propagate signal, localized region labels collapse it—explains why smNSF’s cell-type conditional signals are strongest in tissues with intermingled, heterogeneous cell types and absent in region-annotated data, and provides a diagnostic for interpreting enrichment patterns in new datasets.

### Three specificity classes organize cell-type conditional posteriors

The cell-type-vs-region distinction above establishes that smNSF’s conditional posteriors carry interpretable cell-type information; the next question is how those conditional responses are distributed across factors. We organize the per-dataset results that follow around the three specificity classes introduced in §2.5—*cell-type enriched, cell-type specific*, and *universal* —computed from each factor’s conditional profile across groups, with cell-type groups specialized to the cell-type conditional case and region or layer groups specialized to the region-or layer-conditional case.

- **Cell-type enriched** factors supply most of the biological findings: spatial programs that are masked by composition effects in the marginal and become visible only under conditioning on one cell type. We flag a factor as enriched for a group when its *L*_1_-ratio exceeds 1.5 ×, the threshold used in the aggregate count above.
- **Cell-type specific** factors are present in the marginal and stay present under their dominant cell type, but deplete toward zero under the others; their identification therefore depends on cross-label depletion rather than the single-group *L*_1_-ratio alone.
- **Universal** factors are nearly invariant to label conditioning—all conditional *L*_1_-ratios close to 1, with near-maximal Shannon entropy across the cell-type axis. These represent tissue-level architecture rather than cell-type specific gene programs.

The three classes are descriptive end-points of a continuous specificity spectrum, parameterized by the conditional-to-marginal *L*_1_ ratios together with the cross-label specificity (depletion or entropy) that distinguishes single-label dominance from broad invariance. The per-dataset walkthroughs that follow name the dominant class for each factor they discuss, and the curated factor panels in each dataset figure show concrete examples of all three classes.

### smNSF resolves cell-type spatial programs in Slide-seqV2 hippocampus

The Slide-seqV2 hippocampus dataset [35] contains 41,783 barcoded beads. For the smNSF analysis, each bead is assigned one group label from a 14-label annotation set. The 14 labels comprise two biologically different label types: ten identify cell types, which are spatially scattered across the tissue, and four identify hippocampal subfield regions, which are spatially localized anatomical domains. This mixed label set provides a direct within-dataset test of whether smNSF behaves differently when the group variable encodes dispersed cell identity versus confined anatomical region. We quantify the distinction using the maximum conditional-to-marginal *L*_1_-ratio for each group across all factors: a group is counted as enriched if at least one factor exceeds the 1.5× threshold. smNSF fit *L* = 10 spatial factors under the MGGP prior (Fig. 8; full conditional posteriors in S14 Fig).

**Fig 8.**
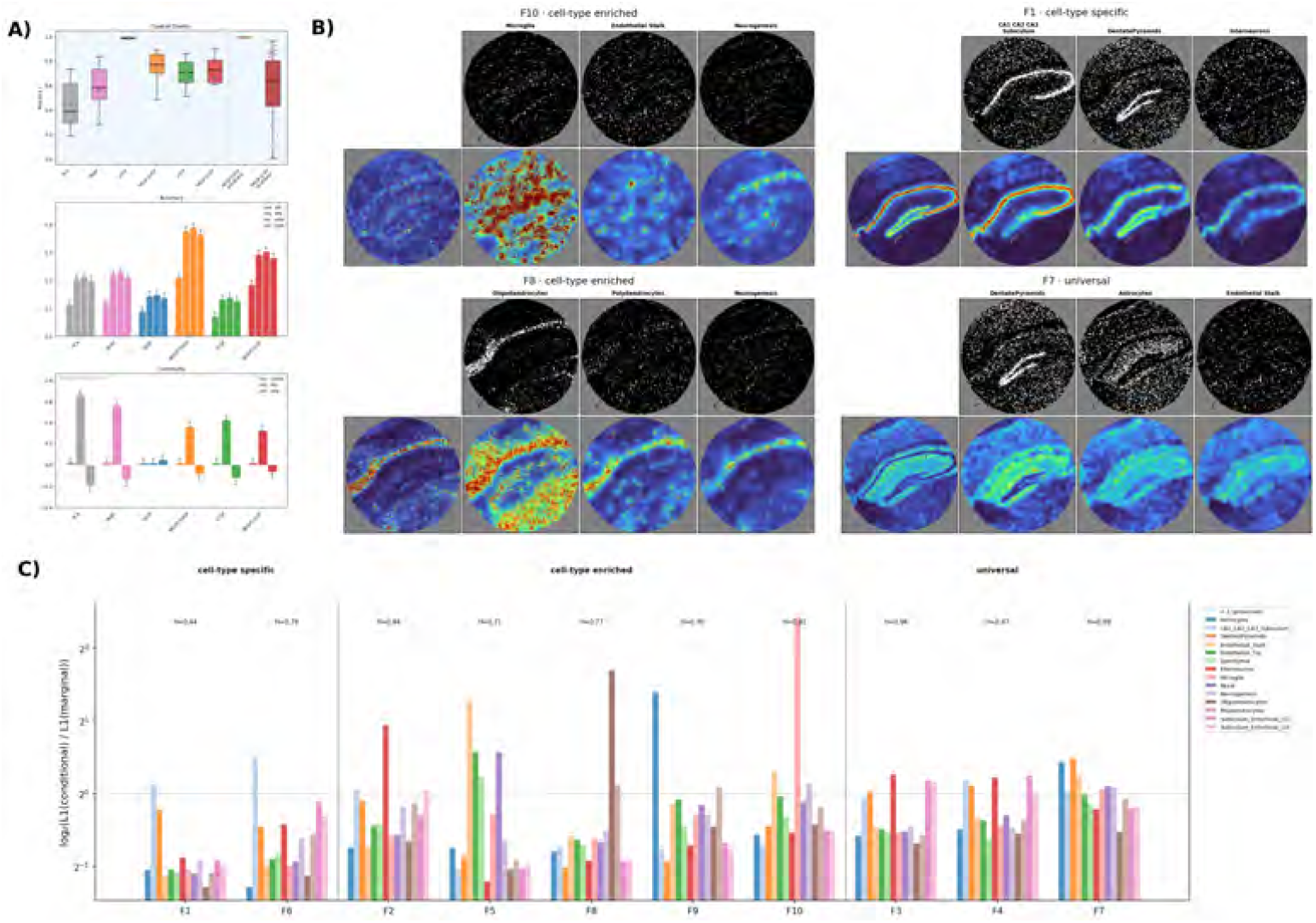
smNSF analysis of Slide-seqV2 mouse hippocampus. **a)** Cross-method benchmark comparison showing spatial quality (Moran’s I), clustering accuracy (ARI, NMI, HOM, COM), and spatial continuity (CHAOS, PAS, ASW) across PCA, PNMF, SVGP, MGGP-SVGP, LCGP, MGGP-LCGP, and a conditional baseline. **b)** Curated cell-type conditional posterior factors (4 representative factors spanning cell-type enriched, cell-type specific, and universal classes). Within each factor block: top row shows the spatial domains of three selected cell types (binary masks of cell-type membership); bottom row shows the marginal factor (leftmost) followed by posteriors conditioned on the cell type shown directly above each column. **c)** Factor specificity quantified as log_2_(∥conditional ∥_1_*/* ∥marginal ∥_1_) per factor per cell type; values above 1 indicate enrichment, below 1 indicate depletion.

The cell-type conditional analysis revealed five cell types with enrichment above 1.5×(Table 3). Conditioning on each cell type revealed a distinct spatial program:

**Table 3.** Factor specificity in Slide-seqV2 hippocampus. Maximum *L*_1_-ratio per group across ten factors. Groups classified as cell type or region.

| Group | Type | Max $L_1$-ratio (Factor) |
| --- | --- | --- |
| Microglia | cell type | $5.36\times$ (F10) |
| Oligodendrocytes | cell type | $3.24\times$ (F8) |
| Astrocytes | cell type | $2.63\times$ (F9) |
| Endothelial Stalk | cell type | $2.43\times$ (F5) |
| Interneurons | cell type | $1.92\times$ (F2) |
| Endothelial Tip | cell type | $1.49\times$ (F5) |
| Mural | cell type | $1.48\times$ (F5) |
| CA1/CA2/CA3 Subiculum | region | $1.41\times$ (F6) |
| Dentate Pyramids | region | $1.40\times$ (F7) |
| Subiculum Entorhinal cl2 | region | $1.19\times$ (F4) |
| Ependymal | cell type | $1.17\times$ (F5) |
| Subiculum Entorhinal cl3 | region | $1.11\times$ (F3) |
| Neurogenesis | cell type | $1.10\times$ (F10) |
| Polydendrocytes | cell type | $1.09\times$ (F8) |

**Microglia** (5.4 ×, Factor 10) showed the strongest enrichment. In the marginal factor, their signal is diluted by the dominant neuronal and glial populations; conditioning on microglia reveals a spatial program that spans the full hippocampal formation, consistent with the broad parenchymal distribution of microglia in the annotated dataset [35].

**Oligodendrocytes** (3.2×, Factor 8) were enriched in a factor that anatomically maps to the white matter tracts of the fimbria and alveus. Conditioning isolates this white-matter compartment: the conditional posterior is strong along the fimbria and weak elsewhere, matching the spatial distribution of oligodendrocyte-annotated bins in the source dataset [35].

**Astrocytes** (2.6 ×, Factor 9) mark a glial niche distinct from the oligodendrocyte program. Although astrocytes are broadly distributed across the hippocampal parenchyma in the annotated dataset [35], smNSF finds that they carry a coherent spatial signal that does not coincide with the dominant neuronal factors.

**Vascular endothelium** (2.4×, Factor 5) was enriched jointly with endothelial tip (1.5 ×) and mural cells (1.5×in the same factor). smNSF discovers this vascular unit without being given any vascular grouping: the three vasculature-associated cell types annotated in the dataset [35] converge on Factor 5 because they share a common spatial program.

**Interneurons** (1.9×, Factor 2) were enriched jointly with CA1/CA2/CA3 subfield neurons. The conditional posterior is strongest in the CA1 subfield, matching the spatial distribution of interneuron-annotated bins in the source dataset [35].

Three additional factors (F3, F4, and F7) showed near-maximal Shannon entropy (0.96–0.99) and were classified as universal—their spatial patterns were nearly unperturbed by cell-type conditioning, representing spatial structure shared broadly across cell types.

In contrast, none of the four hippocampal subfield regions (CA1/CA2/CA3, dentate pyramids, subiculum/entorhinal clades) included groups that exceeded 1.5× enrichment. The strongest region signal was CA1/CA2/CA3 at 1.41× in Factor 6, compared to 5.36×for microglia. This within-dataset contrast—scattered cell types enriched up to 5.4×while localized subfields stall below 1.5 × —is the within-dataset signature of the same MGGP behavior: scattered cell-type labels let conditional signal propagate across the tissue and produce strong factor enrichments, while spatially confined region labels restrict signal to their own subfield and suppress it.

Across methods, smNSF achieved moderate spatial autocorrelation (Moran’s I = 0.66 vs. SVGP: 0.99; Fig. 8a), consistent with its cell-type specific factorization producing spatially localized factors rather than the globally smooth patterns of the non-grouped GP.

### smNSF separates macrophage, stromal, and T-cell programs in MERFISH colorectal cancer

We applied smNSF to a 6 × subsample of Patient 1 from the HuColonCa-FFPE MERFISH dataset [22, 36] (137,693 cells, 211 genes, 13 cell-type groups). smNSF fit *L* = 10 spatial factors that exhibited distinct anatomical localizations across the tissue (Fig. 9; full grid in S15 Fig).

**Fig 9.**
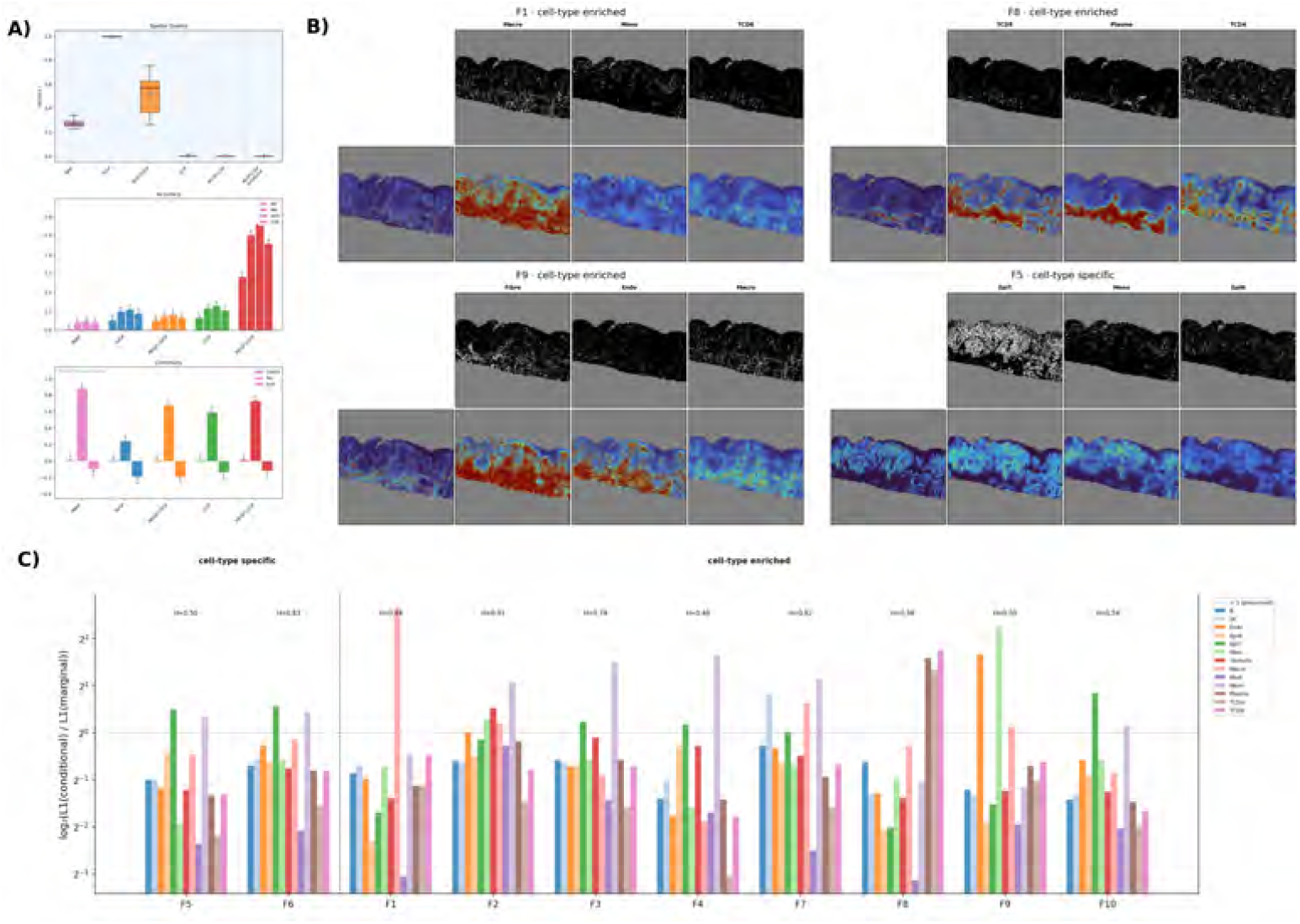
smNSF analysis of MERFISH human colorectal cancer (HuColonCa-FFPE). **a)** Cross-method benchmark comparison. **b)** Curated cell-type conditional posterior factors across specificity classes (cell-type enriched, cell-type specific, universal). Within each factor block: top row shows spatial domains of the selected cell types (binary masks); bottom row shows the marginal factor (leftmost) followed by posteriors conditioned on the cell type above each column. **c)** Factor specificity bar chart showing per cell-type enrichment or depletion patterns.

The cell-type conditional analysis revealed three dominant spatial programs, each defined by a distinct set of co-enriched cell types (Fig. 9b, Table 4):

**Table 4.**
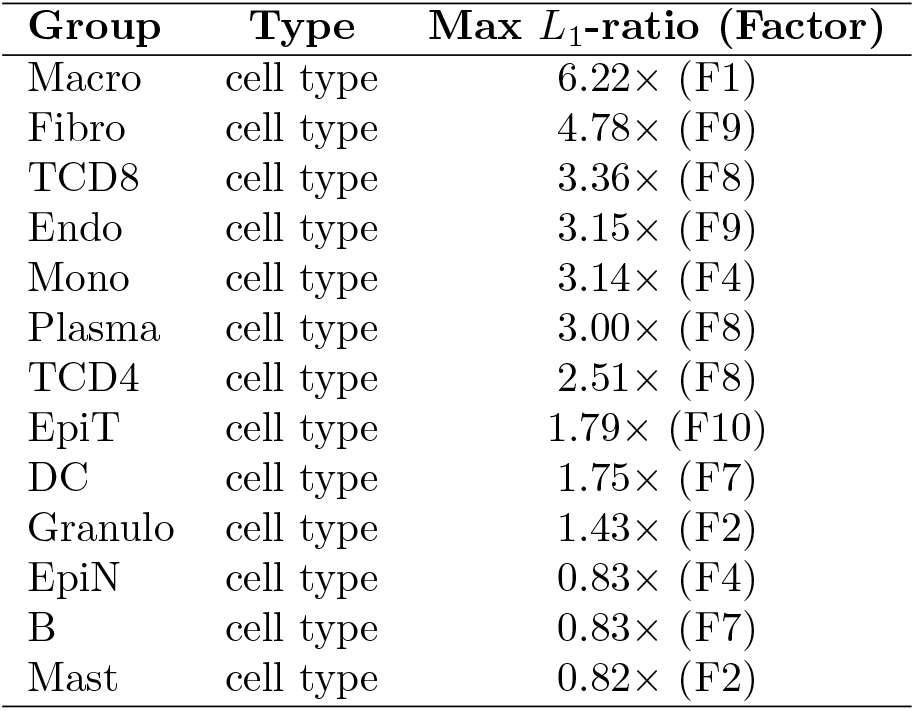
Factor specificity in MERFISH colorectal cancer. Maximum *L*_1_-ratio per group across ten factors. All 13 groups are cell types.

#### Macrophage program (Factor 1, 6.2×)

Macrophages were the most strongly enriched cell type. The conditional posterior reveals a spatial pattern concentrated at the tumor periphery, consistent with the distribution of macrophages along the invasive front in the source dataset [36]. Without conditioning, this signal is masked by the dominant epithelial and stromal populations that fill the tumor bulk.

#### Stromal–vascular niche (Factor 9, fibroblasts: 4.8×, endothelial cells: 3.2×)

Fibroblasts and endothelial cells co-enrich in a single factor. smNSF recovers this stromal–vascular unit without being given any spatial annotation of the tumor microenvironment. The conditional posterior maps to the stromal bands that separate tumor epithelial nests [36].

#### T-cell infiltrate program (Factor 8, CD8^+^ T cells: 3.4×, CD4^+^ T cells: 2.5×, plasma cells: 3.0 ×)

Three adaptive immune cell types co-enrich in the same factor, identifying a coordinated immune infiltrate program. Mast cells, in contrast, are strongly depleted in this factor (0.1×), indicating that their spatial distribution is anti-correlated with the T-cell/plasma cell compartment. This separation—adaptive immune cells on one side, mast cells on the other—mirrors the functional organization of the tumor immune microenvironment, where lymphoid and myeloid compartments occupy distinct spatial niches [36].

smNSF’s per cell-type factors showed near-zero spatial autocorrelation (Moran’s I *<* 0.001; Fig. 9a), reflecting the disentanglement of cell-type specific domains—each factor is spatially active only where its associated cell types reside, in contrast to SVGP’s uniform spatial autocorrelation (0.999) from over-smoothing across cell-type boundaries.

### smNSF recovers the oligodendrocyte lineage and interneuron subtypes in osmFISH

The osmFISH mouse cortex dataset [37] (4,727 cells, 33 genes, 28 groups: 26 cell types and 2 region annotations) provides a dense cell-type taxonomy for testing smNSF’s ability to resolve fine-grained subtypes. smNSF fit *L* = 10 spatial factors (Fig. 10; full grid in S16 Fig).

**Fig 10.**
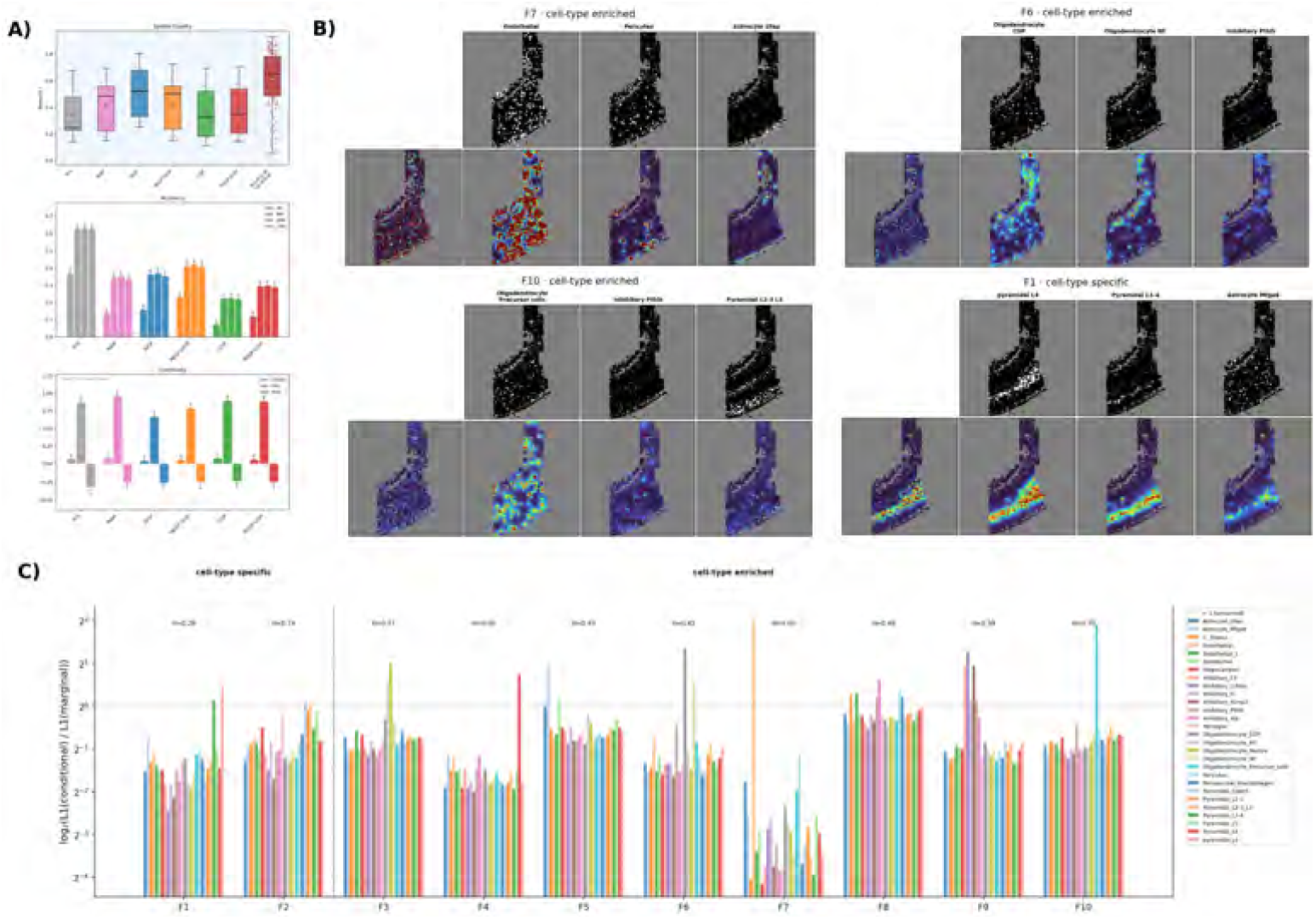
smNSF analysis of osmFISH mouse cortex. **a)** Cross-method benchmark comparison. **b)** Curated cell-type conditional posterior factors across specificity classes (cell-type enriched, cell-type specific, universal). Within each factor block: top row shows spatial domains of the selected cell types (binary masks); bottom row shows the marginal factor (leftmost) followed by posteriors conditioned on the cell type above each column. **c)** Factor specificity bar chart showing per cell-type enrichment or depletion patterns.

Ten cell types showed enrichment above 1.5×(Table 5), and the enriched factors partitioned into biologically interpretable programs:

**Table 5.** Factor specificity in osmFISH mouse cortex. Maximum *L*_1_-ratio per group across ten factors. Full 28-group table in Supplement.

| Group | Type | Max $L_1$-ratio (Factor) |
| --- | --- | --- |
| Endothelial | cell type | $4.10 \times$ (F7) |
| Oligodendrocyte precursor | cell type | $3.73 \times$ (F10) |
| Oligodendrocyte COP | cell type | $2.55 \times$ (F6) |
| Inhibitory Crhbp | cell type | $2.42 \times$ (F9) |
| Oligodendrocyte Mature | cell type | $2.02 \times$ (F3) |
| Astrocyte Mfge8 | cell type | $1.98 \times$ (F5) |
| Inhibitory Kcnip2 | cell type | $1.93 \times$ (F9) |
| Inhibitory CP | cell type | $1.92 \times$ (F9) |
| Pyramidal L6 | cell type | $1.66 \times$ (F4) |
| Inhibitory Vip | cell type | $1.54 \times$ (F8) |
| *(16 additional cell types in Supplement)* |  |  |
| C. plexus | region | $1.21 \times$ (F8) |
| Hippocampus | region | $0.85 \times$ (F8) |

#### Vascular program (Factor 7, endothelial cells: 4.1×)

Endothelial cells were strongly enriched and nearly absent from all neuronal populations. This is the same vascular signal seen across datasets—endothelial cells form blood vessels, a spatially coherent structure that is independent of neuronal or glial organization. The conditional posterior maps to the tissue’s microvasculature.

#### Oligodendrocyte lineage (Factors 10, 6, and 3)

The annotated oligodendrocyte lineage stages [37]—oligodendrocyte precursor cells (OPCs), committed oligodendrocyte precursors (COP), and mature oligodendrocytes—were resolved across three separate factors: OPCs enriched Factor 10 (3.7×), COPs enriched Factor 6 (2.6×), and mature oligodendrocytes enriched Factor 3 (2.0×). smNSF separates immature and mature oligodendrocyte stages into distinct spatial factors without being given any developmental ordering.

#### Inhibitory interneuron program (Factor 9)

Three inhibitory interneuron subtypes—Crhbp (2.4×), Kcnip2 (1.9 ×), and CP (1.9 ×)—converge on Factor 9. These subtypes are annotated as distinct cell types in the source dataset [37], yet smNSF assigns them to a shared spatial factor, indicating that their spatial distributions in this cortical section overlap.

#### Astrocyte and pyramidal neuron factors (Factors 5 and 4)

Astrocyte Mfge8 selectively enriched Factor 5 (2.0 ×), and deep-layer pyramidal L6 neurons enriched Factor 4 (1.7 ×). These enrichments are more moderate, consistent with the broader spatial distributions of astrocyte-annotated and L6-annotated cells in this section [37]; their conditional posteriors are less sharply localized than those of the vascular or oligodendrocyte programs.

The two region groups—C. plexus and hippocampus—showed no enrichment above 1.5× (max 1.21 ×), consistent with the cell-type versus region distinction observed across all datasets.

smNSF achieved moderate spatial autocorrelation (Moran’s I = 0.38 vs. SVGP: 0.51; Fig. 10a), balancing spatial continuity with the cell-type specificity needed to resolve 28 groups including 26 fine-grained subtypes.

### Human liver: macrophage–stellate sinusoidal program across conditions

We analyzed nine MERFISH human liver samples (three healthy, six fibrotic) [44] with approximately 100,000 cells each. Here we focus on a matched healthy–diseased pair from donor AM048 (Fig. 11–12); per-sample panels for the other seven samples are in S17–S23 Figs; conditional posteriors for all nine samples in S24–S32 Figs.

**Fig 11.**
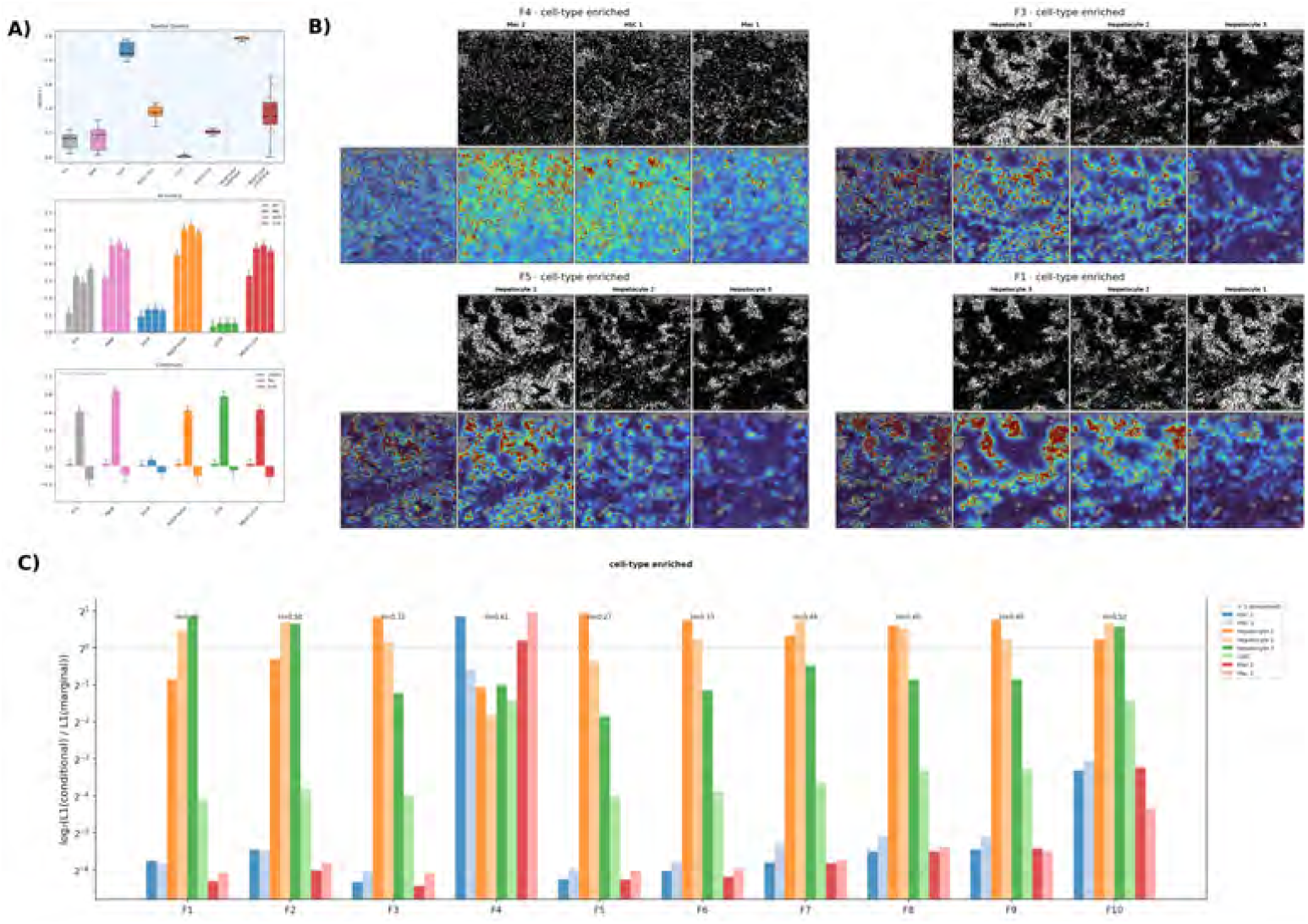
smNSF analysis of MERFISH human liver — healthy sample AM048. **a)** Cross-method benchmark comparison. **b)** Curated cell-type conditional posterior factors across specificity classes (cell-type enriched, cell-type specific, universal). Within each factor block: top row shows spatial domains of the selected cell types (binary masks); bottom row shows the marginal factor (leftmost) followed by posteriors conditioned on the cell type above each column. **c)** Factor specificity bar chart showing per cell-type enrichment or depletion patterns.

**Fig 12.**
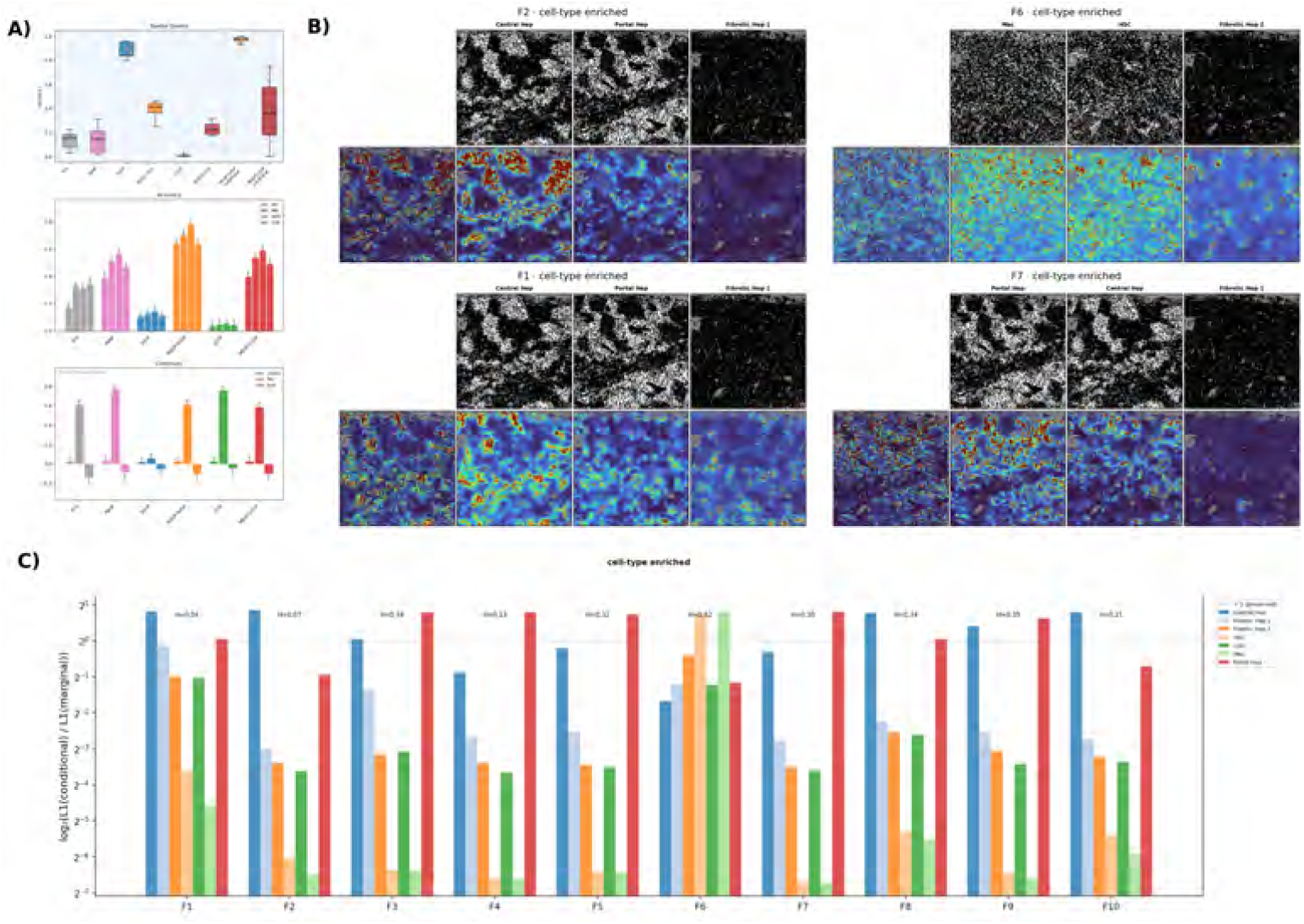
smNSF analysis of MERFISH human liver — diseased sample AM048. **a)** Cross-method benchmark comparison. **b)** Curated cell-type conditional posterior factors across specificity classes (cell-type enriched, cell-type specific, universal). Within each factor block: top row shows spatial domains of the selected cell types (binary masks); bottom row shows the marginal factor (leftmost) followed by posteriors conditioned on the cell type above each column. **c)** Factor specificity bar chart showing per cell-type enrichment or depletion patterns.

Across all nine samples, hepatic stellate cells (HSC) and macrophages (mac) were the most consistently enriched cell types in *L*_1_-ratio, reaching 1.7–1.9× in AM048 and up to 5.3× in other samples. In AM048 healthy liver, mac 2 and HSC 1 co-enriched in

Factor 4 (1.9 ×and 1.8 ×, respectively; Table 6), while in AM048 diseased liver, mac and HSC co-enriched in Factor 6 (1.8× and 1.7×). smNSF assigns these two annotated cell types to a shared spatial factor; in the source dataset, HSCs are annotated as perisinusoidal and macrophages as sinusoidal Kupffer cells [44]. In both conditions, mac and HSC are depleted from each other’s secondary factors, indicating that smNSF captures their coupled spatial co-occurrence without being given their anatomical proximity.

**Table 6.**
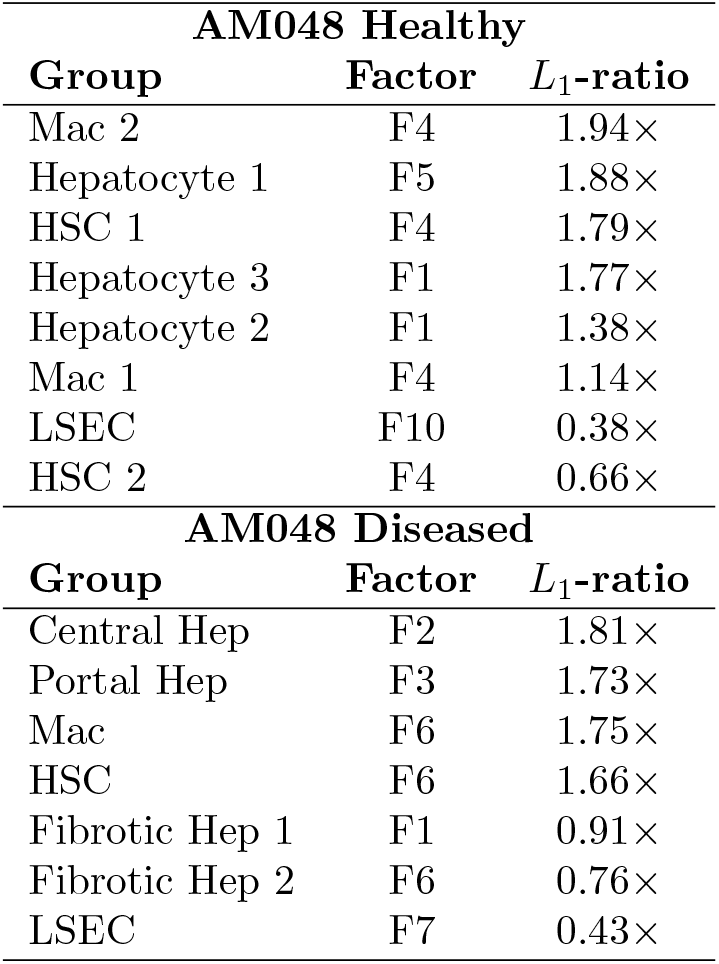
Factor specificity in human liver sample AM048. Top *L*_1_-ratio per group. Healthy (eight groups) and diseased (seven groups) from matched donors.

| AM048 Healthy |  |  |
| --- | --- | --- |
| Group | Factor | $L_1$-ratio |
| Mac 2 | F4 | 1.94× |
| Hepatocyte 1 | F5 | 1.88× |
| HSC 1 | F4 | 1.79× |
| Hepatocyte 3 | F1 | 1.77× |
| Hepatocyte 2 | F1 | 1.38× |
| Mac 1 | F4 | 1.14× |
| LSEC | F10 | 0.38× |
| HSC 2 | F4 | 0.66× |
| AM048 Diseased |  |  |
| Group | Factor | $L_1$-ratio |
| Central Hep | F2 | 1.81× |
| Portal Hep | F3 | 1.73× |
| Mac | F6 | 1.75× |
| HSC | F6 | 1.66× |
| Fibrotic Hep 1 | F1 | 0.91× |
| Fibrotic Hep 2 | F6 | 0.76× |
| LSEC | F7 | 0.43× |

Hepatocyte subtypes (central hep, portal hep, hepatocyte 1/2/3) showed moderate enrichment (1.8–1.9 ×in AM048 healthy; 1.8× for central hep and portal hep in AM048 diseased), consistent with metabolic zonation being a continuous pericentral-to-periportal gradient rather than a discrete group signal. The MGGP’s group-discrete kernel is inherently less sensitive to continuous spatial gradients—this is a feature, not a bug, as it prevents zonation from dominating the factorization.

In fibrotic samples, disease-emergent compartments appeared: Fibrotic hepatocyte populations were depleted below 1.0× in AM048 diseased (fibrotic hep 1: 0.91 ×, fibrotic hep 2: 0.76×), indicating that the fibrotic niche loses spatial coherence relative to the dominant sinusoidal program. In other diseased samples (S19–S23 Figs), annotated fibrotic hepatocyte subtypes [44] formed their own spatially coherent compartments at up to 3.3× enrichment, suggesting smNSF can detect disease-specific spatial restructuring when the fibrotic signal is sufficiently strong.

### smNSF on volumetric MERFISH hypothalamus

We applied smNSF to MERFISH data from the mouse hypothalamic preoptic region (71,939 cells across multiple z-slices, 161 genes, 11 cell-type groups) [39]. This is the only three-dimensional dataset in our study, and the only one where every one of the ten factors was classified as cell-type enriched (Fig. 13; full conditional posteriors in S33 Fig). The hypothalamus dataset contains many spatially structured, molecularly distinct annotated cell classes [39], which likely drives the uniformly high enrichment rate: each factor captures a different compartment of the annotated tissue.

**Fig 13.**
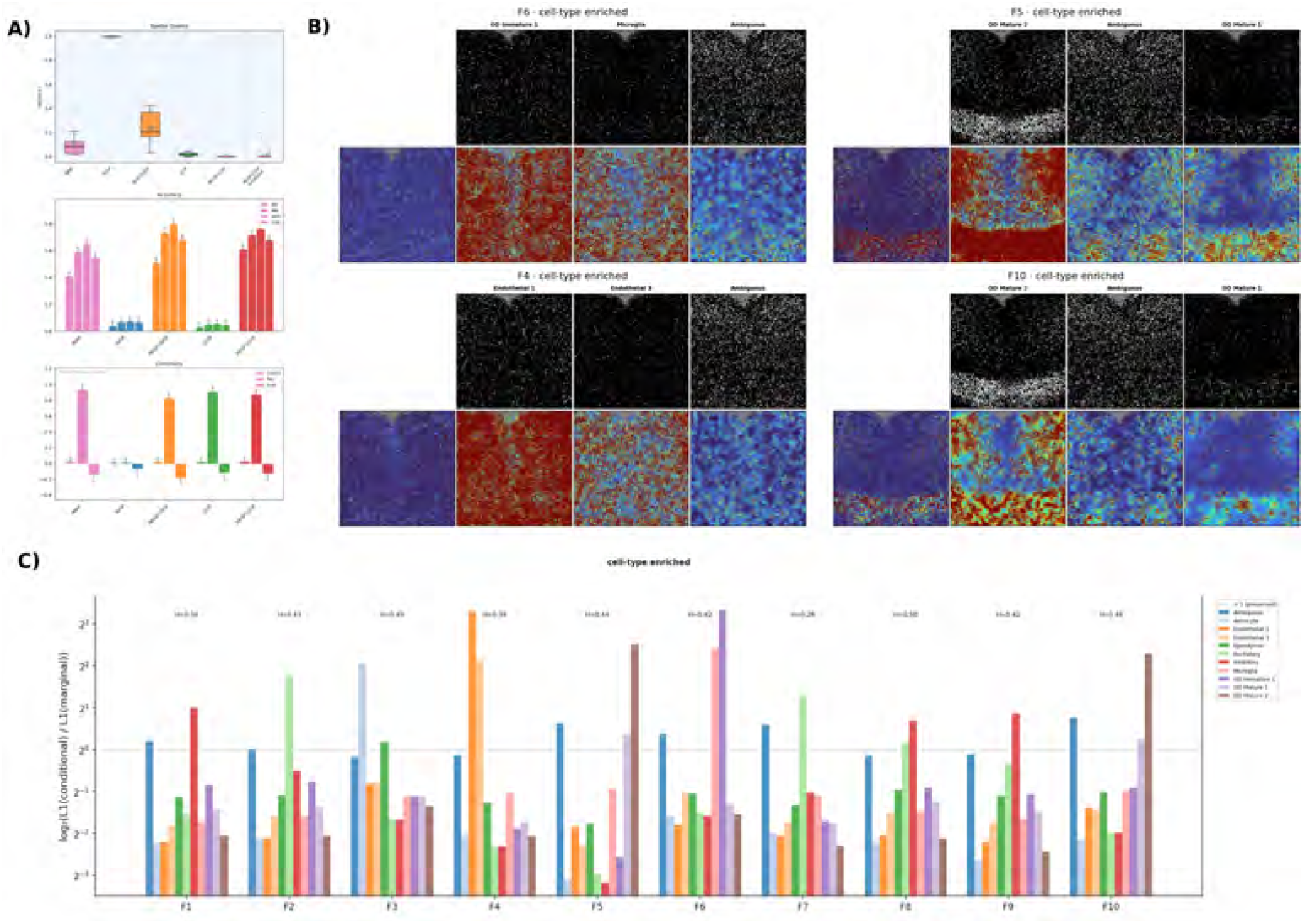
smNSF analysis of MERFISH mouse hypothalamus. **a)** Cross-method benchmark comparison. **b)** Curated cell-type conditional posterior factors across specificity classes (cell-type enriched, cell-type specific, universal). Within each factor block: top row shows spatial domains of the selected cell types (binary masks); bottom row shows the marginal factor (leftmost) followed by posteriors conditioned on the cell type above each column. **c)** Factor specificity bar chart showing per cell-type enrichment or depletion patterns.

The strongest *L*_1_-ratio enrichments reached 10.1× (OD immature 1 in Factor 6) and 9.9 × (endothelial 1 in Factor 4), the highest *L*_1_-ratios observed across any dataset (Fig. 13b, Table 7). Four biological programs emerged:

**Table 7.**
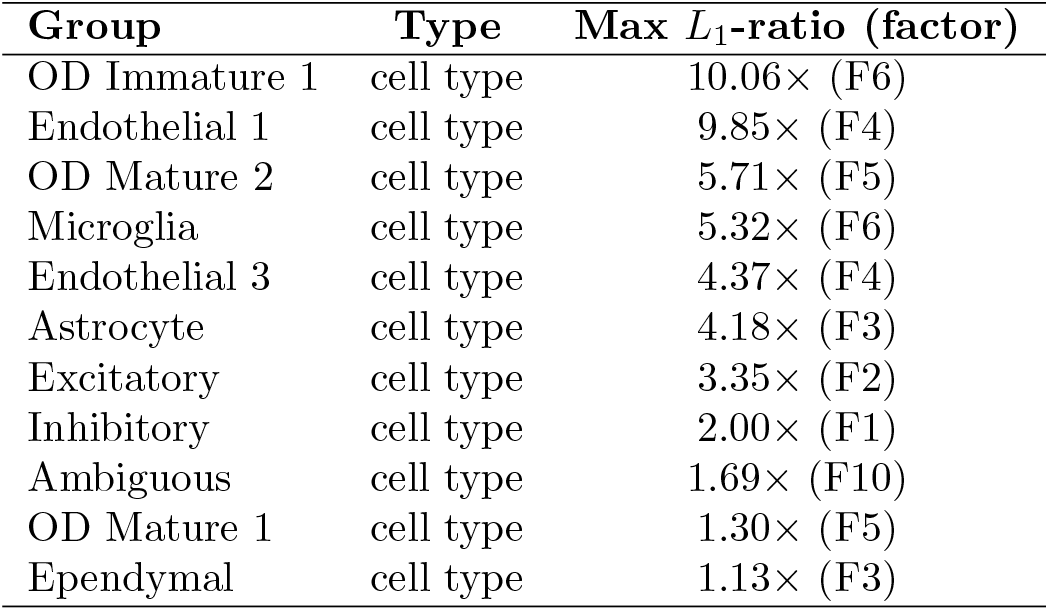
Factor specificity in MERFISH mouse hypothalamus. Maximum *L*_1_-ratio per group across ten factors. All eleven group labels are cell types; 9/11 are enriched above 1.5× in specific factors (the highest rate across datasets).

#### Oligodendrocyte lineage (Factors 6, 5, and 10)

As in osmFISH, immature and mature oligodendrocyte populations [39] separated into distinct factors: OD immature 1 dominated Factor 6 (10.1 ×) while OD mature 2 enriched Factors 5 (5.7×) and 10 (4.9 ×). The separation of the two annotated stages into different spatial axes is recovered by smNSF without supervision.

#### Vascular program (Factor 4)

Endothelial 1 (9.9×) and endothelial 3 (4.4 ×) co-enriched in Factor 4. This is the same vascular signal observed in Slide-seqV2, colon, and osmFISH—endothelial cells consistently co-enrich because they share the spatial architecture of the tissue’s blood vessel network, regardless of species or brain region.

#### Neuronal excitation–inhibition split (Factors 2, 7, 1, and 9)

Excitatory and inhibitory neurons [39] were cleanly assigned to separate factors: excitatory neurons were enriched in Factors 2 (3.4×) and 7 (2.5×), while inhibitory neurons were enriched in Factors 1 (2.0×) and 9 (1.8×). This separation is not imposed by smNSF—the model receives only cell-type labels, not neurotransmitter identity—yet it recovers the excitatory/inhibitory division annotated in the source dataset into distinct spatial axes.

#### Astrocyte program (Factor 3)

Astrocytes were selectively enriched in Factor 3 (4.2 ×) with depletion in most other groups. Their isolation into a single factor indicates a coherent spatial organization that is independent of the neuronal and vascular programs in this section [39].

As in colon, smNSF’s factors showed near-zero spatial autocorrelation (Moran’s I *<* 0.001; Fig. 13a), consistent with the dataset’s cell-type annotations and the MGGP’s disentanglement of cell-type specific spatial domains.

### smNSF generalizes to 10x Visium with region-only annotations

We applied smNSF to a 10x Visium adult mouse brain coronal section [38] (2,688 spots, 16,944 genes, 15 groups). All 15 groups are anatomically defined brain regions (cortex subregions, hippocampus, thalamus, hypothalamus, striatum, fiber tracts). Consistent with the cell-type versus region label distinction, no group exceeded 1.5× *L*_1_-ratio enrichment (Table 8). The greatest enrichment was fiber tract at 1.38 ×in Factor 8.

**Table 8.** Factor specificity in 10x Visium mouse brain. All fifteen groups are brain regions; none exceed 1.5× *L*_1_-ratio enrichment.

| Group | Type | Max $L_1$-ratio (Factor) |
| --- | --- | --- |
| Fiber_tract | region | $1.38 \times$ (F8) |
| Lateral_ventricle | region | $1.23 \times$ (F10) |
| Thalamus_1 | region | $1.11 \times$ (F3) |
| Hippocampus | region | $0.96 \times$ (F9) |
| Cortex_1 | region | $0.95 \times$ (F2) |
| Hypothalamus_1 | region | $0.92 \times$ (F1) |
| Striatum | region | $0.91 \times$ (F4) |
| Thalamus_2 | region | $0.90 \times$ (F6) |
| Cortex_5 | region | $0.90 \times$ (F9) |
| Cortex_3 | region | $0.90 \times$ (F2) |
| Cortex_2 | region | $0.85 \times$ (F5) |
| Hypothalamus_2 | region | $0.84 \times$ (F6) |
| Cortex_4 | region | $0.83 \times$ (F7) |
| Pyramidal_layer_dentate_gyrus | region | $0.75 \times$ (F9) |
| Pyramidal_layer | region | $0.70 \times$ (F7) |

Despite the absence of group-level enrichment, smNSF recovered anatomically structured factors (Fig. 14; full grid in S34 Fig), with hippocampus, cortex, and white matter regions occupying distinct factor axes. The weaker specificity relative to MERFISH datasets reflects both the region-only annotation and the inherent per-spot cell-type mixing of Visium’s 55 µm spots. This dataset serves as a negative control confirming that region annotations, regardless of platform, do not produce enriched conditional posteriors under the MGGP prior.

**Fig 14.**
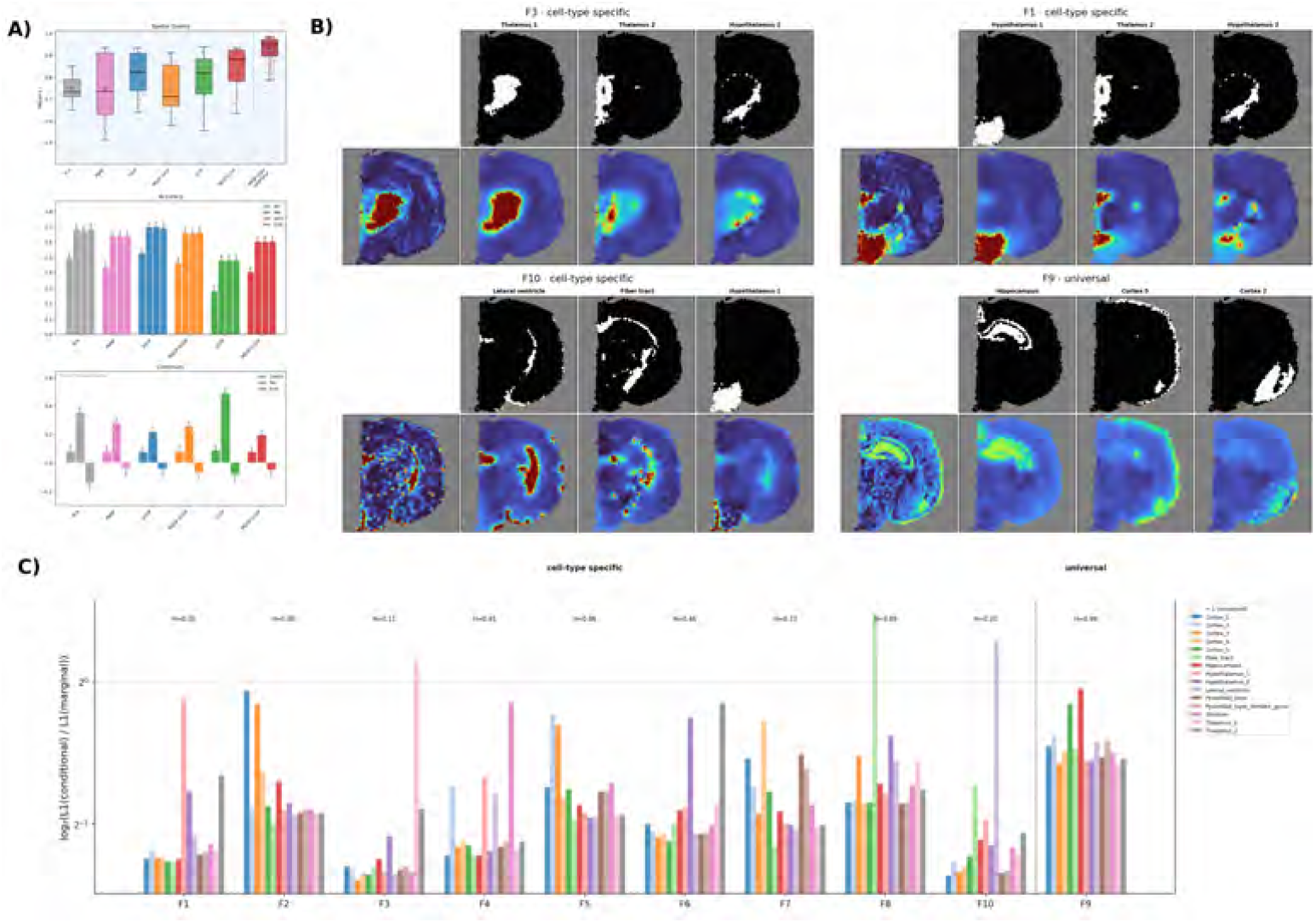
smNSF analysis of 10x Visium adult mouse brain coronal section. **a)** Cross-method benchmark comparison. **b)** Curated region-conditional posterior factors across specificity classes. Within each factor block: top row shows spatial domains of the selected brain regions (binary masks); bottom row shows the marginal factor (leftmost) followed by posteriors conditioned on the region above each column. **c)** Factor specificity bar chart showing per-region enrichment or depletion patterns.

smNSF achieved the highest spatial autocorrelation among methods (Moran’s I = 0.84 vs. SVGP: 0.82, LCGP: 0.79; Fig. 14a), consistent with the pattern on DLPFC. For region-only annotations, the MGGP prior provides stronger spatial regularization than standard GPs: the group-discrete kernel reduces correlation across groups, so signal propagates less between samples from different groups. This effect is amplified for region labels because each region occupies a confined spatial domain—unlike scattered cell types, region-specific signal cannot spread beyond its anatomical boundary.

## Discussion and Conclusion

smNSF addresses a fundamental challenge in spatial transcriptomics: distinguishing spatial patterns driven by gene regulation within a cell type from those driven by shifts in cell-type composition across space. By conditioning the GP prior on cell-type labels through multi-group kernels, smNSF places each spatial factor on a specificity spectrum—from universal factors shared across all cell types, to cell-type enriched factors, to factors dominated by a single cell type—and supports counterfactual queries of the form “what would this factor look like if the tissue were composed entirely of cell type *X*?”

A consistent structural property of the MGGP prior emerged from the cross-dataset analysis: enrichment depends on whether group labels are spatially scattered or confined in a local neighborhood. Cell-type labels, typically intermingled across the tissue, allow cell-type specific signal to be shared globally and produce cell-type enriched conditional posteriors in 61% of groups (46/75). On the other hand, region labels, locally confined by definition, never exceeded the 1.5× enrichment threshold in conditional posteriors (0/28). Within datasets containing both annotation types —Slide-seqV2 hippocampus and osmFISH cortex—this distinction held. This behavior is not a limitation but a diagnostic: it indicates when smNSF will yield interpretable cell-type specific factors and when the given annotation structure precludes it.

Recovered factors in suitable datasets with purely cell-type labels captured dataset-consistent spatial organization. For example, smNSF finds that hepatic stellate cells and macrophages are co-enriched in a shared sinusoidal program across healthy and fibrotic liver, the oligodendrocyte lineage resolves across three factors in osmFISH cortex, and macrophage, stromal, vascular, and T-cell compartments separate cleanly in specific colorectal tumor samples.

The LCGP inference framework is independently useful. By approximating the variational posterior locally rather than globally, LCGP allows *M* = *N* on datasets with tens of thousands of cells (demonstrated on the Slide-seqV2 hippocampus) and lifts the *O*(*M* ^3^) ceiling that has effectively capped GP-based methods at a few thousand inducing points; the same inference machinery scales to larger datasets. The approach is not specific to smNSF and applies to any latent-variable GP model. The computational trade-offs favor LCGP when the number of cells exceeds ~ 40,000 or when capturing local spatial heterogeneity is important; for smaller datasets with smooth spatial patterns, standard SVGP remains efficient.

Several considerations apply to use of our method.

### Group labels and label quality

smNSF requires discrete group labels for conditional analysis: labels capturing cell types, anatomical regions, or cortical layers were used across our datasets. When those labels are inferred cell types rather than ground-truth annotations—most ST datasets lack the latter, so labels are typically obtained from unsupervised clustering or label transfer from reference scRNA-seq data [10, 13]—label quality propagates into the conditional posteriors: the MGGP prior treats the supplied labels as known, so noisy or misassigned labels may smooth or distort the per-group factor maps.

### Equidistant group structure

The choice of group distance 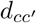 in the MGGP kernel currently treats all distinct group pairs as equidistant. Incorporating cell-type hierarchies or continuous embeddings could improve modeling of related cell states.

### Volumetric data

While we demonstrate extensions to 3D data, systematic evaluation of smNSF on volumetric tissues remains future work.

### Factor capacity

Our analyses used *L* = 10 factors across all datasets. With richer factorizations (*L* = 50–100), smNSF could in principle reconstruct the full dataset and support gene-expression imputation under specific cell-type compositions—asking, for example, how the gene expression of the tissue would appear if only macrophages were present. More work is needed to scale the MGGP to higher latent dimensions—either by scaling multi-group LCGP or using a different approximation for the multi-group GP—while maintaining numerical stability.

### In silico, not causal

The cell-type conditional posterior is an *in silico* query under the fitted MGGP prior rather than a prediction of an experimental perturbation: it answers how the model expects spatial factors to change for a hypothetical cell-type composition, given the data, but does not establish a causal effect of cell-type identity on gene expression. Causal interpretation would require an experimental intervention—e.g., targeted ablation of a cell population or directed differentiation—and the model’s predictions for such interventions would need separate validation.

smNSF’s conditional posteriors describe how spatial factors shift when conditioning on a cell type, not how individual genes vary across cell types. This makes factor-level validation more difficult: the method cannot directly answer questions such as “is gene X upregulated in macrophages specifically?” The gene programs underlying each factor are encoded in the loadings *W*, but linking those programs to a specific cell type and not to individual factors that cannot be conditioned on cell type would require a different model.

Our method provides a foundation for several extensions. The MGGP framework can incorporate multiple grouping variables (e.g., cell type and disease state) through product kernels, and could be combined with multi-sample extensions of NSF such as mNSF [20] to enable joint cell-type aware factorization across samples. Integration with differential expression analysis could identify genes whose spatial patterns differ across conditions while controlling for cell-type composition. Also, combining smNSF with velocity or trajectory inference could reveal how spatial gene programs evolve during development or disease progression.

Taken together, smNSF with LCGP addresses two bottlenecks that have limited the adoption of GP-based spatial factorization methods: the inability to disentangle group-driven from gene-expression-driven spatial patterns, and the computational ceiling that restricted spatial GP priors to a few thousand observations. The consistent enrichment pattern across 75 cell-type and 28 region groups (103 groups total; Table 2) provides a practical diagnostic: if group labels are scattered, the MGGP prior enriches conditionally; if they are spatially confined, it does not—giving us a clear criterion for when conditional analysis is likely to be informative. More broadly, the group structure in the MGGP prior is not restricted to cell types: groups can encode treatment conditions, time points, disease states, donors, or other experimental covariates, providing a generative framework to ask biological questions directly within a dimension reduction framework.

## Supporting information

Supplemental Figures and Tables for NNSF

## Data Availability

All datasets analyzed in this study are publicly available. Slide-seqV2 mouse hippocampus data are from Stickels et al. (2021) [35], accessed via the Squidpy tutorial data repository [38]. osmFISH mouse somatosensory cortex data are from Codeluppi et al. (2018) [37], accessed via the Squidpy tutorial data repository [38]. MERFISH mouse hypothalamus data are from Moffitt et al. (2018) [39], accessed via the Squidpy tutorial data repository [38]. 10x Visium adult mouse brain data (Coronal Section 1) are from 10x Genomics, accessed via the Squidpy tutorial data repository [38]. Human colorectal cancer MERFISH data (HuColonCa-FFPE) are from the Vizgen MERFISH data release, accessed via Laursen et al. (2026) [22]. Human DLPFC Visium data are from Maynard et al. (2021) [40] via the spatialLIBD resource (http://spatial.libd.org/), accessed through the SDMBench benchmark suite [41]. Human liver MERFISH data are from Watson et al. (2025) [44].

All code is available at https://github.com/luisdiaz1997/GPzoo (MGGP and LCGP inference), https://github.com/luisdiaz1997/Probabilistic-NMF (nonnegative and spatial nonnegative factorizations), and https://github.com/luisdiaz1997/Spatial-Factorization (full smNSF pipeline).

## AI-assisted writing disclosure

The authors used large language model tools, including ChatGPT and Claude, to assist with manuscript editing, organization, and wording during drafting and revision. The authors reviewed and edited all AI-assisted text, verified factual claims and citations, and take full responsibility for the final content. No AI tools were used to generate primary data or to perform the reported statistical analyses.

## Acknowledgments

LCD, PS, and BEE were funded in part by grants from the Parker Institute for Cancer Immunology (PICI), the Chan-Zuckerberg Institute (CZI), the Biswas Family Foundation, NIH NHGRI R01 HG012967, and NIH NHGRI R01 HG013736. BEE is a CIFAR Fellow in the Multiscale Human Program.

BEE is on the Scientific Advisory Board for ArrePath Inc, GSK AI for Cancer, and Freenome; she consults for Neumora.

## Supporting Information

**S1 Fig. MGGP prior illustration on a 2D toy dataset**. Samples drawn from the Matérn-3/2 MGGP prior across four group-kernel lengthscales (*a* ∈ {0.01, 0.1, 1.0, 10.0}) over four stripe-pattern ground-truth group configurations, visualized as 3D surfaces in both a smooth regime and a noisy regime. Illustrates how the group-difference parameter *a* controls sharing between groups (cell types): small *a* yields near-identical across-group factors; large *a* yields group-specific factors.

**S2 Fig. MGGP-LCGP group-conditional spatial factors for DLPFC slice 151507**. Full grid of group-conditional spatial factors for Maynard et al. DLPFC slice 151507 (Visium; 7 annotated cortical layers).

**S3 Fig. MGGP-LCGP group-conditional spatial factors for DLPFC slice 151508**. Full grid of group-conditional spatial factors for Maynard et al. DLPFC slice 151508.

**S4 Fig. MGGP-LCGP group-conditional spatial factors for DLPFC slice 151509**. Full grid of group-conditional spatial factors for Maynard et al. DLPFC slice 151509.

**S5 Fig. MGGP-LCGP group-conditional spatial factors for DLPFC slice 151510**. Full grid of group-conditional spatial factors for Maynard et al. DLPFC slice 151510.

**S6 Fig. MGGP-LCGP group-conditional spatial factors for DLPFC slice 151669**. Full grid of group-conditional spatial factors for Maynard et al. DLPFC slice 151669.

**S7 Fig. MGGP-LCGP group-conditional spatial factors for DLPFC slice 151670**. Full grid of group-conditional spatial factors for Maynard et al. DLPFC slice 151670.

**S8 Fig. MGGP-LCGP group-conditional spatial factors for DLPFC slice 151671**. Full grid of group-conditional spatial factors for Maynard et al. DLPFC slice 151671.

**S9 Fig. MGGP-LCGP group-conditional spatial factors for DLPFC slice 151672**. Full grid of group-conditional spatial factors for Maynard et al. DLPFC slice 151672.

**S10 Fig. MGGP-LCGP group-conditional spatial factors for DLPFC slice 151673**. Full grid of group-conditional spatial factors for Maynard et al. DLPFC slice 151673.

**S11 Fig. MGGP-LCGP group-conditional spatial factors for DLPFC slice 151674**. Full grid of group-conditional spatial factors for Maynard et al. DLPFC slice 151674.

**S12 Fig. MGGP-LCGP group-conditional spatial factors for DLPFC slice 151675**. Full grid of group-conditional spatial factors for Maynard et al. DLPFC slice 151675.

**S13 Fig. MGGP-LCGP group-conditional spatial factors for DLPFC slice 151676**. Full grid of group-conditional spatial factors for Maynard et al. DLPFC slice 151676.

**S14 Fig. MGGP-LCGP group-conditional spatial factors for Slide-seqV2 mouse hippocampus**. Full grid of all *L* = 10 smNSF spatial factors conditioned on each of the 14 annotated groups (10 cell types and 4 hippocampal subfield regions) in

the Slide-seqV2 hippocampus dataset (41,783 spatial barcodes). Complements the curated subset shown in Fig 2 and the consolidated Slide-seq main-text panel.

**S15 Fig. MGGP-LCGP cell-type conditional spatial factors for HuColonCa-FFPE MERFISH human colorectal cancer**. Full grid of cell-type conditional spatial factors for the HuColonCa-FFPE MERFISH human colorectal cancer dataset (137,693 cells after 6× subsample, 13 cell-type groups collapsed from ~ 50 fine annotations). Complements the curated subset shown in the colon main-text panel.

**S16 Fig. MGGP-LCGP cell-type conditional spatial factors for osmFISH mouse somatosensory cortex**. Full grid of cell-type conditional spatial factors for the osmFISH mouse cortex dataset (4,727 cells, 33 genes, 28 cell types). Complements the curated subset shown in the osmFISH main-text panel.

**S17 Fig. smNSF analysis of MERFISH human liver — healthy sample AM042**. Per-sample panel for the healthy AM042 liver sample.

**S18 Fig. smNSF analysis of MERFISH human liver — healthy sample AM061**. Per-sample panel for the healthy AM061 liver sample.

**S19 Fig. smNSF analysis of MERFISH human liver — diseased sample AM031**. Per-sample panel for the diseased AM031 liver sample.

**S20 Fig. smNSF analysis of MERFISH human liver — diseased sample AM042**. Per-sample panel for the diseased AM042 liver sample.

**S21 Fig. smNSF analysis of MERFISH human liver — diseased sample AM061**. Per-sample panel for the diseased AM061 liver sample.

**S22 Fig. smNSF analysis of MERFISH human liver — diseased sample AM062**. Per-sample panel for the diseased AM062 liver sample.

**S23 Fig. smNSF analysis of MERFISH human liver — diseased sample AM072**. Per-sample panel for the diseased AM072 liver sample.

**S24 Fig. MGGP-LCGP cell-type conditional spatial factors for MERFISH human liver — healthy sample AM048**. Full grid of cell-type conditional spatial factors for the healthy AM048 liver sample.

**S25 Fig. MGGP-LCGP cell-type conditional spatial factors for MERFISH human liver — healthy sample AM042**. Full grid of cell-type conditional spatial factors for the healthy AM042 liver sample.

**S26 Fig. MGGP-LCGP cell-type conditional spatial factors for MERFISH human liver — healthy sample AM061**. Full grid of cell-type conditional spatial factors for the healthy AM061 liver sample.

**S27 Fig. MGGP-LCGP cell-type conditional spatial factors for MERFISH human liver — diseased sample AM031**. Full grid of cell-type conditional spatial factors for the diseased AM031 liver sample.

**S28 Fig. MGGP-LCGP cell-type conditional spatial factors for MERFISH human liver — diseased sample AM042**. Full grid of cell-type conditional spatial factors for the diseased AM042 liver sample.

**S29 Fig. MGGP-LCGP cell-type conditional spatial factors for MERFISH human liver — diseased sample AM048**. Full grid of cell-type conditional spatial factors for the diseased AM048 liver sample.

**S30 Fig. MGGP-LCGP cell-type conditional spatial factors for MERFISH human liver — diseased sample AM061**. Full grid of cell-type conditional spatial factors for the diseased AM061 liver sample.

**S31 Fig. MGGP-LCGP cell-type conditional spatial factors for MERFISH human liver — diseased sample AM062**. Full grid of cell-type conditional spatial factors for the diseased AM062 liver sample.

**S32 Fig. MGGP-LCGP cell-type conditional spatial factors for MERFISH human liver — diseased sample AM072**. Full grid of cell-type conditional spatial factors for the diseased AM072 liver sample.

**S33 Fig. MGGP-LCGP cell-type conditional spatial factors for MERFISH mouse hypothalamus**. Full grid of cell-type conditional spatial factors for the MERFISH hypothalamus dataset (Moffitt2018; 71,939 cells, 161 genes, 11 groups).

**S34 Fig. MGGP-LCGP group-conditional spatial factors for 10x Visium Adult Mouse Brain (Coronal Section 1)**. Full grid of group-conditional spatial factors for the 10x Visium mouse brain slice (2,688 spots, 16,944 genes, 15 brain region groups). Complements the curated subset shown in the Visium main-text panel.

**S35 Fig. Convergence of the three multiplicative-update methods on a synthetic nonnegative-matrix benchmark**. Side-by-side comparison of the Jensen lower bound (Method 1), the expanded/hybrid estimator (Method 2), and full Monte Carlo (Method 3) when factorizing a random nonnegative matrix into *L* = 10 components. The expanded estimator is substantially more stable than full Monte Carlo while attaining a lower final loss than the analytic lower bound, motivating its use as the default in all spatial experiments.

**S36 Fig. Kernel-conditioned neighbor selection remains stable as data density grows**. Each panel shows the *K*=50 neighbors (orange) of the same query point in subsamples of the Slide-seqV2 hippocampus dataset at *N* ∈ {5,000, 10,000, 40,000} cells. **Top row:** baseline Euclidean KNN—the neighborhood radius collapses with increasing *N*. **Bottom row:** LCGP with kernel-conditioned neighbor selection (RBF kernel, lengthscale *l*=8.0)—the neighborhood extent stays well-behaved across all densities.

**S37 Fig. LCGP replaces the global KL with** *M* **local KLs**. *Left:* the exact MGGP-SVGP KL connects every inducing point to every other and costs *O* (*M* ^3^). *Middle:* LCGP samples a neighbor set *n*(*j*) of size *K* for each inducing point *j* with probability proportional to the MGGP kernel *k* ((*z*_*j*_, *c*_*j*_)), (*z*_*i*_, *c*_*i*_); same-group neighbors (blue circles) are preferred but cross-group neighbors (red triangles) enter the set whenever the kernel ranks them highly. *Right:* repeating this sampling at every inducing point yields the LCGP connectivity graph—a sparse graph whose local KL costs O(*MK*^3^), making *M* = *N* inducing points tractable for the KL term.

**S38 Fig. Probabilistic vs**. *K***-nearest-neighbor selection in LCGP on Slide-seqV2 hippocampus**. Both blocks are MGGP-LCGP fits at *a* = 10^6^ (so the multi-group coupling is effectively removed and the only remaining difference is the neighbor-selection rule). Each block shows the unconditional spatial factor map (top row) and two cell-type conditional posteriors below; the leftmost column of each row marks which cells belong to that group. Reference factors and cell-type rows are picked from the VNNGP/KNN side (left block): the top two factors by max-over-groups *L*_1_ specificity ratio, with each factor’s argmax-enriched cell type as its row (Oligodendrocytes, Astrocytes). The corresponding columns of the LCGP/probabilistic block are not the same numerical factor indices but are the best Pearson matches on the per-factor gene-loading vectors, so that the same biological gene program is being compared across methods. Driving selection from the VNNGP side is the conservative framing: the baseline picks the factors and cell-type rows where it shows the strongest signal, so the LCGP block’s differences (more anatomically structured conditional maps on the same rows) cannot be attributed to LCGP-favorable factor selection.

