## Supplemental Figures and Tables for NNSF for "Scalable multi-group nonnegative spatial factorization for spatial genomics data with cell-type heterogeneity"

### Supporting information

#### S1 Variational inference for MGGPs

We develop a complete variational inference framework for multi-group Gaussian processes (MGGPs) [1], instantiated for the smNSF observation model that extends nonnegative spatial factorization (NSF) [2] with cell-type-conditional latent factors. Let $X$ denote spatial inputs with labels $C_X$, inducing locations $Z$ with labels $C_Z$, and latent values $F$ and $U$ at those locations.

##### S1.1 Prior specification

The MGGP prior over the latent function values is:

$$p(F \mid X, C_X; \theta) = \mathcal{MGGP}(F \mid 0, K((X, C_X), (X, C_X))).$$

We can express this prior as a marginalization over inducing variables:

$$p(F|X, C_X; \theta) = \int p(F|U, X, C_X, Z, C_Z; \theta)\, p(U|Z, C_Z; \theta)\, dU$$

where the inducing distribution is also an MGGP:

$$p(U|Z, C_Z; \theta) = \mathcal{MGGP}(U \mid 0, K((Z, C_Z), (Z, C_Z))).$$

For our implementation, inducing points $Z$ are chosen from the distribution of inputs within each group, ensuring adequate coverage across all cell types.

##### S1.2 ELBO derivation

The log marginal likelihood is:

$$\log p(Y|X, C_X; \theta) = \log \int p(Y|F)\, p(F|U, X, C_X, Z, C_Z; \theta)\, p(U|Z, C_Z; \theta)\, dU\, dF$$

Introducing variational distributions $q(F, U)$ and applying Jensen's inequality:

$$\log p(Y|X, C_X; \theta) \geq \int \log \frac{p(Y|F)\, p(F|U, X, C_X, Z, C_Z; \theta)\, p(U|Z, C_Z; \theta)}{q(F, U)}\, q(F, U)\, dU\, dF$$

Following [3], we set $q(F|U) = p(F|U)$, giving:

$$q(F, U) = p(F|U, X, C_X, Z, C_Z; \theta)\, q(U)$$

This simplifies the ELBO to:

$$\log p(Y|X, C_X; \theta) \geq \mathbb{E}_{q(F)}[\log p(Y|F)] - \mathrm{KL}\big[q(U) \,\|\, p(U|Z, C_Z; \theta)\big] \tag{1}$$

The expectation term is approximated via Monte Carlo sampling for non-Gaussian likelihoods (the full expansion for the smNSF Poisson likelihood is given in §Inference and Training in smNSF); the closed form of the KL divergence is derived in §KL divergence computation.

##### S1.3 Variational distribution specification

We use a mean-field factorization across latent factors:

$$q(U) = \prod_{l=1}^{L} q(u_\ell), \qquad q(u_\ell) = \mathcal{N}(u_\ell \mid m_\ell, L_\ell L_\ell^T)$$

where $L_\ell$ is a lower-triangular Cholesky factor ensuring positive definiteness.

##### S1.4 Posterior marginalization

The variational posterior over latent function values at the data locations is obtained by marginalizing the inducing variables:

$$q(f_\ell \mid X, C_X; \theta) = \int p(f_\ell \mid u_\ell, X, C_X, Z, C_Z; \theta)\, q(u_\ell)\, du_\ell.$$

This is the standard SVGP predictive form [4], obtained via Gaussian marginalization (§Gaussian Marginalization):

$$q(f_\ell) = \mathcal{N}\Big(m_\ell^f,\ S_\ell^f\Big),$$

with moments

$$\begin{aligned} m_\ell^f &= K_{xz}(K_{zz})^{-1} m_\ell, \\ S_\ell^f &= K_{xx} - K_{xz}(K_{zz})^{-1}K_{zx} + K_{xz}(K_{zz})^{-1} L_\ell L_\ell^\top (K_{zz})^{-1} K_{zx}. \end{aligned}$$

The per-data-point marginal at spatial location $x_i$ with group label $c_i$ and factor $\ell$ is therefore

$$q(f_{i\ell}) = \mathcal{N}\Big(m_\ell^f(x_i, c_i),\ S_\ell^f((x_i, c_i), (x_i, c_i))\Big),$$

where $m_\ell^f(x_i, c_i)$ is the entry of $m_\ell^f$ corresponding to observation $i$ with spatial coordinate $x_i$ and group $c_i$, and $S_\ell^f((x_i, c_i), (x_i, c_i))$ is the corresponding diagonal element of $S_\ell^f$. The full latent representation at observation $i$ factorizes as

$$q(F_{i\cdot}) = \prod_{\ell=1}^{L} q(f_{i\ell}),$$

a consequence of the mean-field structure over factors in (1). The Poisson expected log-likelihood expansion in §Inference and Training in smNSF requires only these marginal means and variances, making the derivations independent of the specific variational parameterization chosen for $q(U)$.

#### S2 KL divergence computation

The KL term in the SVGP ELBO (1) between the variational posterior $q(U)$ and the inducing-prior $p(U \mid Z, C_Z; \theta)$ admits a closed form [3, 5]. Under the mean-field factorization across factors, it decomposes as:

$$\mathrm{KL}[q(U) \| p(U)] = \sum_{l=1}^{L} \mathrm{KL}[q(u_\ell) \| p(u_\ell)]$$

We work in the whitened parameterization $u_\ell = L g_\ell$ where $LL^\top = K_{zz}$ and $p(g_\ell) = \mathcal{N}(0, I)$; the KL divergence is invariant under this change of variables [5, 6]. With $q(g_\ell) = \mathcal{N}(m_\ell^g, L_\ell^g L_\ell^{g\top})$, the closed form follows from the Gaussian KL identity (§KL divergence):

$$\mathrm{KL}[q(g_\ell) \| p(g_\ell)] = \frac{1}{2}\left[-2\sum_{m=1}^{M} \log(L_\ell^g)_{mm} - M + \|L_\ell^g\|_F^2 + \|m_\ell^g\|_2^2\right]$$

where $(L_\ell^g)_{mm}$ denotes the $m$-th diagonal element and $\|\cdot\|_F$ is the Frobenius norm.

#### S2.1 Predictive posteriors

A key feature of our variational framework is that the posterior depends only on the corresponding inputs $(X, C_X)$ [4]. This enables prediction at new spatial locations or under altered cell-type configurations:

$$q(F \mid X_{\text{new}}, C_{X_{\text{new}}}; \theta) = \int p(F \mid U, X_{\text{new}}, C_{X_{\text{new}}}, Z, C_Z; \theta)\, q(U)\, dU$$

This allows *in silico* perturbation experiments where we fix spatial coordinates $X$ and vary the cell-type labels $C_X$ to assess cell-type specificity of each spatial factor.

### S3 Non-separable Matérn MGGP kernel

We extend the multi-group Gaussian process construction of [1], which provides the framework for valid covariances on $\mathbb{R}^p \times \mathcal{C}$ (where $\mathcal{C}$ is a finite set of group labels), to an inseparable Matérn form. Separable space–group kernels can produce undesirable smoothness artifacts at group boundaries (the "ridges" phenomenon discussed in the spatio-temporal context by [7]); we therefore embed group relationships directly into the spatial covariance with a Matérn form that depends jointly on spatial and inter-group distances. Let $x, x' \in \mathbb{R}^p$ denote spatial coordinates, $c_i, c_j$ group labels, $d_{ij}$ a graph-based group distance, and $a \geq 0$ a scale controlling how strongly group separation modulates spatial smoothness. The multi-group Matérn kernel is

$$K((x, c_i), (x', c_j)) = \frac{\sigma^2}{\left(a^2 d_{ij}^2 + 1\right)^{p/2}} \frac{2^{1-\nu}}{\Gamma(\nu)} \left( \frac{\sqrt{2\nu}\, \|x - x'\|}{\ell \sqrt{a^2 d_{ij}^2 + 1}} \right)^{\nu} K_\nu \left( \frac{\sqrt{2\nu}\, \|x - x'\|}{\ell \sqrt{a^2 d_{ij}^2 + 1}} \right), \tag{2}$$

where $\ell$ and $\sigma^2$ are the base spatial length scale and marginal variance, $\nu$ controls smoothness, and $K_\nu(\cdot)$ is a modified Bessel function of the second kind. The denominator penalizes cross-group pairs in proportion to $d_{ij}$, shrinking both the signal variance and the effective length scale when groups differ. Setting $a = 0$ (all groups identical) recovers the usual Matérn kernel, whereas increasing $a$ enlarges cross-group distances and attenuates correlation between dissimilar groups.

##### Reduction to the standard Matérn

When $a = 0$ the $(1 + a^2 d_{ij}^2)$ factors equal 1 for all $i, j$, yielding

$$K((x, c_i), (x', c_i)) = \sigma^2 \frac{2^{1-\nu}}{\Gamma(\nu)} \left( \frac{\sqrt{2\nu}\, \|x - x'\|}{\ell} \right)^{\nu} K_\nu \left( \frac{\sqrt{2\nu}\, \|x - x'\|}{\ell} \right),$$

the canonical Matérn kernel with parameters $(\sigma^2, \ell, \nu)$.

##### Closed-form special cases

Let $r = \|x - x'\|$. For half-integer $\nu$ the Bessel term simplifies to polynomials in $r$ times exponentials:

$\nu = \frac{1}{2}$

$$K((x, c_i), (x', c_j)) = \sigma^2 \frac{1}{\left(1 + a^2 d_{ij}^2\right)^{p/2}} \exp\left( - \frac{r}{\ell \sqrt{1 + a^2 d_{ij}^2}} \right).$$

$\nu = \frac{3}{2}$

$$K((x, c_i), (x', c_j)) = \sigma^2 \, \frac{1}{\big(1 + a^2 d_{ij}^2\big)^{p/2}} \left(1 + \frac{\sqrt{3}\, r}{\ell \sqrt{1 + a^2 d_{ij}^2}}\right) \exp\left(- \frac{\sqrt{3}\, r}{\ell \sqrt{1 + a^2 d_{ij}^2}}\right).$$

$\nu = \frac{5}{2}$

$$K((x, c_i), (x', c_j)) = \sigma^2 \, \frac{1}{\big(1 + a^2 d_{ij}^2\big)^{p/2}} \left(1 + \frac{\sqrt{5}\, r}{\ell \sqrt{1 + a^2 d_{ij}^2}} + \frac{5\, r^2}{3\, \ell^2 \left(1 + a^2 d_{ij}^2\right)}\right) \exp\left(- \frac{\sqrt{5}\, r}{\ell \sqrt{1 + }}\right.$$ [illegible]

These closed forms make it straightforward to evaluate gradients of $\ell$, $\sigma$, $a$, or $\nu$ without special-function libraries.

#### Positive definiteness

Write $\alpha = \nu + \frac{p}{2}$ and $\lambda = \sqrt{2\nu}/\ell$. By Bochner's theorem (§Bochner's theorem and complete monotonicity) $K((x, c_i), (x', c_j)) = \int e^{i\omega^\top (x - x')} \, S_{ij}(\omega) \, d\omega$ with spectra

$$S_{ij}(\omega) = \sigma^2 C_{\nu,p} \, (1 + a^2 d_{ij}^2)^{-\alpha} \Big( \|\omega\|^2 + \frac{\lambda^2}{1 + a^2 d_{ij}^2} \Big)^{-\alpha} =: \Phi_\omega(a^2 d_{ij}^2),$$

where $C_{\nu,p}$ collects constants independent of $i, j$. For fixed $\omega$ we need $[\Phi_\omega(a^2 d_{ij}^2)]_{ij} \succeq 0$. Using

$$(1 + t)^{-\alpha} \Big( \|\omega\|^2 + \frac{\lambda^2}{1 + t} \Big)^{-\alpha} = \big( \|\omega\|^2 (1 + t) + \lambda^2 \big)^{-\alpha},$$

and that $x \mapsto x^{-\alpha}$ is completely monotone for $\alpha > 0$, $\Phi_\omega(t)$ is completely monotone on $[0, \infty)$. Schoenberg's theorem then implies $[\Phi_\omega(a^2 d_{ij}^2)]_{ij} \succeq 0$ for every $\omega$ whenever $[d_{ij}^2]$ is a squared-distance matrix of negative type, so the kernel in Eq. 2 is positive definite for any such group distance matrix and any $a \geq 0$. Taking $\nu \to \infty$ in the spectral density (using $\Gamma(\nu + p/2)/\Gamma(\nu) \sim \nu^{p/2}$ and the standard limit $(1 + x/\nu)^{-\nu} \to e^{-x}$) yields $S_{ij}(\omega) \propto \exp\big( - \frac{\ell^2}{2} \|\omega\|^2 (1 + a^2 d_{ij}^2) \big)$, recovering the non-separable RBF MGGP for any $a \geq 0$, so the MGGP Matérn strictly generalizes that family while retaining probabilistic validity.

### S4 Bochner's theorem and complete monotonicity

We collect the classical positive-definiteness results invoked in §Non-separable Matérn MGGP kernel; for textbook treatments see [8]. Bochner's theorem states a continuous stationary kernel $K(x - x')$ on $\mathbb{R}^p$ is positive-definite iff it is the Fourier transform of a finite non-negative measure $\mu(\omega)$:

$$K(r) = \int_{\mathbb{R}^p} e^{i\omega^\top r} \, d\mu(\omega) = \int_{\mathbb{R}^p} \cos(\omega^\top r) \, d\mu(\omega), \quad r = x - x'.$$

If $\mu$ has density $S(\omega) \geq 0$, then $K(r) = \int_{\mathbb{R}^p} e^{i\omega^\top r} \, S(\omega) \, d\omega$. For multi-output kernels the spectral density becomes PSD: $K_{ij}(r) = \int e^{i\omega^\top r} \, S_{ij}(\omega) \, d\omega$.

A function $\phi : [0, \infty) \to \mathbb{R}$ is completely monotone (CM) if $(-1)^k \phi^{(k)}(t) \geq 0$ for all $k \geq 0$. Bernstein's theorem implies CM functions are Laplace transforms of finite non-negative measures. Schoenberg's theorem links CM functions to radial PD kernels: if $\phi$ is CM, then $K(x, x') = \phi(\|x - x'\|^2)$ is PD for all ambient dimensions, and conversely global positive definiteness of such radial kernels implies $\phi$ is CM. Mixtures of exponentials (e.g., rational-quadratic, Matérn) therefore remain PD across dimensions, justifying the distance-dependent factors of the MGGP Matérn kernel.

### S5 Locally Conditioned KL Approximation for Multivariate Gaussian Variational Inference

We approximate the KL divergence between the variational distribution $q(U)$ and the prior $p(U)$ using a locally conditional chain rule factorization. Let $U = [U_1, \ldots, U_M]$ be the collection of inducing variables. The geometric picture (S37 Fig) is that the dense all-pairs connectivity of the global KL (cost $\mathcal{O}(M^3)$) is replaced by a sparse graph whose local KL terms cost $\mathcal{O}(MK^3)$ to evaluate, with each inducing point connected to a kernel-weighted neighborhood that prefers same-group neighbors but admits cross-group ones when the kernel ranks them highly.

We begin by expanding the $p(U)$ prior using the chain rule:

$$\begin{aligned} p(U) &= p(U_M|U_{1,\ldots,M-1})p(U_{1,\ldots,M-1}) \\ &= p(U_M|U_{1,\ldots,M-1})p(U_{M-1}|U_{1,\ldots,M-2})p(U_{1,\ldots,M-2}) \\ &= p(U_1)\prod_{j=2}^{M} p(U_j|U_{1,\ldots,j-1}). \end{aligned}$$

Under the local conditional approximation, this becomes:

$$p(U) \approx \prod_{j=1}^{M} p(U_j|U_{n(j)}).$$

Then, the variational distribution $q(U)$ can be expanded similarly:

$$\begin{aligned} q(U) &= q(U_1)\prod_{j=2}^{M} q(U_j|U_{1,\ldots,j-1}) \\ &\approx \prod_{j=1}^{M} q(U_j|U_{n(j)}) \end{aligned}$$

The KL divergence between $q(U)$ and $p(U)$ is given by:

$$\mathrm{KL}(q(U)\|p(U)) = \mathbb{E}_{q(U)}\left[\log\frac{q(U)}{p(U)}\right]$$

Next, the log ratio can be expanded and approximated as the product of $M$ terms:

$$\mathbb{E}_{q(U)}\left[\log\frac{q(U)}{p(U)}\right] \approx \mathbb{E}_{q(U)}\left[\log\prod_{j=1}^{M}\frac{q(U_j|U_{n(j)})}{p(U_j|U_{n(j)})}\right]$$

Becoming a linear sum of expectations:

$$= \mathbb{E}_{q(U)}\left[\sum_{j=1}^{M}\log\frac{q(U_j|U_{n(j)})}{p(U_j|U_{n(j)})}\right]$$

$$= \sum_{j=1}^{M} \mathbb{E}_{q(U)} \left[ \log \frac{q(U_j | U_{n(j)})}{p(U_j | U_{n(j)})} \right]$$

We then apply **Law of total expectation** which gives:

$$= \sum_{j=1}^{M} \mathbb{E}_{q(U_{n(j)})} \left[ \mathbb{E}_{q(U_j | U_{n(j)})} \left[ \log \frac{q(U_j | U_{n(j)})}{p(U_j | U_{n(j)})} \right] \right]$$

We assume that both the prior and variational posterior distributions over $U = [U_1, \ldots, U_M]$ are multivariate Gaussian. In particular, we write

$$p(U) = \mathcal{N}(U \mid 0, K), \qquad q(U) = \mathcal{N}(U \mid m, S),$$

where $K$ and $S$ denote the prior and variational covariance matrices, and $m = [m_1, \ldots, m_M]$ is the variational mean. Here we assume a zero-mean prior.
The conditional prior is given by:

$$p(U_j | U_{n(j)}) = \mathcal{N} \left( K_{jn(j)} K_{n(j)n(j)}^{-1} U_{n(j)},\ k_{jj} - K_{jn(j)} K_{n(j)n(j)}^{-1} K_{n(j)j} \right)$$

The conditional variational distribution is given by:

$$q(U_j | U_{n(j)}) = \mathcal{N} \left( m_j + S_{jn(j)} S_{n(j)n(j)}^{-1} (U_{n(j)} - m_{n(j)}),\ s_{jj} - S_{jn(j)} S_{n(j)n(j)}^{-1} S_{n(j)j} \right)$$

This reduces to $q(U_j) = \mathcal{N}(m_j, s_{jj})$ if $s_{jj}$ is diagonal.
For notational compactness in subsequent expressions, we define these conditionals:

$$p(U_j | U_{n(j)}) = \mathcal{N} \left( \beta_j^\top U_{n(j)},\ \tau_j^2 \right)$$
$$q(U_j | U_{n(j)}) = \mathcal{N} \left( m_j + \alpha_j^\top (U_{n(j)} - m_{n(j)}),\ \tilde{\tau}_j^2 \right)$$

Where:

$$\beta_j^\top = K_{jn(j)} K_{n(j)n(j)}^{-1}, \quad \tau_j^2 = k_{jj} - K_{jn(j)} K_{n(j)n(j)}^{-1} K_{n(j)j}$$
$$\alpha_j^\top = S_{jn(j)} S_{n(j)n(j)}^{-1}, \quad \tilde{\tau}_j^2 = s_{jj} - S_{jn(j)} S_{n(j)n(j)}^{-1} S_{n(j)j}$$

The KL divergence between two univariate Gaussians is:

$$\mathrm{KL}(q \,\|\, p) = \frac{1}{2} \left[ \log \frac{s_p^2}{s_q^2} + \frac{s_q^2}{s_p^2} + \frac{(\mu_q - \mu_p)^2}{s_p^2} - 1 \right].$$

The KL between $q(U_j | U_{n(j)})$ and $p(U_j | U_{n(j)})$ is given by:

$$\mathrm{KL}\big(q(U_j \mid U_{n(j)}) \,\|\, p(U_j \mid U_{n(j)})\big) =$$
$$\frac{1}{2} \left[ \log \frac{\tau_j^2}{\tilde{\tau}_j^2} + \frac{\tilde{\tau}_j^2}{\tau_j^2} + \frac{\big(m_j + \alpha_j^\top (U_{n(j)} - m_{n(j)}) - \beta_j^\top U_{n(j)}\big)^2}{\tau_j^2} - 1 \right].$$

Then we calculate the expectation of this KL under $q(U_{n(j)})$:

$$\Rightarrow \mathbb{E}_{q(U_{n(j)})} \left[ \mathrm{KL} \left( q(U_j | U_{n(j)}) \| p(U_j | U_{n(j)}) \right) \right]$$

$$= \frac{1}{2}\left[\log\frac{\tau_j^2}{\tilde{\tau}_j^2} + \frac{\tilde{\tau}_j^2}{\tau_j^2} + \frac{\mathbb{E}_{q(U_{n(j)})}\left[\left(m_j + \alpha_j^\top(U_{n(j)} - m_{n(j)}) - \beta_j^\top U_{n(j)}\right)^2\right]}{\tau_j^2} - 1\right].$$

We treat the mean term:

$$\left(m_j + \alpha_j^\top(U_{n(j)} - m_{n(j)}) - \beta_j^\top U_{n(j)}\right)^2$$

This is rewritten as:

$$= \left((\alpha_j - \beta_j)^\top U_{n(j)} + (m_j - \alpha_j^\top m_{n(j)})\right)^2$$

Using the following identity:

$$\mathbb{E}_{x\sim\mathcal{N}(\mu,\Sigma)}\left[(a^\top x + b)^2\right] = a^\top \Sigma a + (a^\top \mu + b)^2$$

Applying this gives:

$$\mathbb{E}_{q(U_{n(j)})}\left[\left(m_j + \alpha_j^\top(U_{n(j)} - m_{n(j)}) - \beta_j^\top U_{n(j)}\right)^2\right] = (\alpha_j - \beta_j)^\top S_{n(j)n(j)}(\alpha_j - \beta_j) + \left[\,(\alpha_j - \beta_j)^\top m_{n(j)} + \left(m_j - \alpha_j^\top m_{n(j)}\right)\right]^2.$$

This expectation can be expanded further and simplified as:

$$= \alpha_j^\top S_{n(j)n(j)}\alpha_j + \beta_j^\top S_{n(j)n(j)}\beta_j - 2\alpha_j^\top S_{n(j)n(j)}\beta_j + (\beta_j^\top m_{n(j)} - m_j)^2$$

Substituting this into the KL formula yields the final approximation:

$$\mathrm{KL}(q(U)\|p(U)) \approx \sum_{j=1}^{M} \mathbb{E}_{q(U_{n(j)})}\left[\mathbb{E}_{q(U_j|U_{n(j)})}\left[\log\frac{q(U_j|U_{n(j)})}{p(U_j|U_{n(j)})}\right]\right]$$

$$= \sum_{j=1}^{M}\frac{1}{2}\left[\log\frac{\tau_j^2}{\tilde{\tau}_j^2} + \frac{\tilde{\tau}_j^2}{\tau_j^2} + \frac{\left(m_j - \beta_j^\top m_{n(j)}\right)^2}{\tau_j^2} + \frac{\beta_j^\top S_{n(j)n(j)}\beta_j}{\tau_j^2} + \frac{\alpha_j^\top S_{n(j)n(j)}\alpha_j}{\tau_j^2} - \frac{2\alpha_j^\top S_{n(j)n(j)}\beta_j}{\tau_j^2} - 1\right].$$

This decomposition allows for scalable computation of the KL divergence in sparse MGGP models using only local neighbor subsets $n(j)$. It is especially useful for variational nearest-neighbor Gaussian processes (VNNGPs) [9], and can be efficiently computed in parallel across all nodes.

### S6 Parameterizing the Variational Covariance

We consider parameterizations for the variational covariance $S$ that admit efficient computation of the local quantities $\alpha_j$ and $\tilde{\tau}_j^2$. We avoid a pure diagonal $S$: as shown in §Diagonal Variational Covariance Underestimates Posterior Variance, diagonal variational covariances elementwise underestimate posterior marginal variances and, under standard kernel jittering, can collapse to zero irrespective of the data. Instead, we develop a low-rank parameterization that retains the dominant between-inducing-point correlations while remaining tractable.

#### S6.1 Low-Rank Factor

Let:

$$S = L_u L_u^\top \tag{3}$$

where $L_u \in \mathbb{R}^{M\times R}$ with $R \le M$, and let $L_{n(j)} \in \mathbb{R}^{|n(j)|\times R}$ denote the rows corresponding to the neighbor set.

The neighborhood covariance and cross-covariance are:

$$S_{n(j)n(j)} = L_{n(j)} L_{n(j)}^\top, \quad S_{jn(j)} = L_j L_{n(j)}^\top \tag{4}$$

If $R < |n(j)|$, this is rank-deficient. Assuming $R \ge |n(j)|$ or adding regularization, the conditional variance is:

$$\tilde{\tau}_j^2 = L_j L_j^\top - L_j L_{n(j)}^\top (L_{n(j)} L_{n(j)}^\top)^{-1} L_{n(j)} L_j^\top \tag{5}$$

**Limitation**: This parameterization forces $\text{rank}(S) \le R$. For full-rank $S$, we need $R = M$, losing the computational benefit.

#### S6.2 Choosing the Rank $R$

The rank $R$ of the low-rank factor $L_u \in \mathbb{R}^{M\times R}$ controls how well $S = L_u L_u^\top$ can approximate the true posterior covariance. To choose $R$ appropriately, we appeal to the spectral structure of the prior kernel $k$.

**Spectral interpretation.** By Mercer's theorem, any continuous, symmetric, positive semi-definite kernel $k : \mathcal{X} \times \mathcal{X} \to \mathbb{R}$ on a compact set $\mathcal{X} \subset \mathbb{R}^d$ admits the expansion:

$$k(\boldsymbol{x}, \boldsymbol{x}') = \lim_{r\to\infty} \sum_{i=1}^{r} \lambda_i \psi_i(\boldsymbol{x}) \psi_i(\boldsymbol{x}'), \tag{6}$$

where $\lambda_1 \ge \lambda_2 \ge \cdots \ge \lambda_r \ge 0$ are eigenvalues and $\psi_i : \mathcal{X} \to \mathbb{R}$ are orthonormal eigenfunctions, with the series converging absolutely and uniformly on $\mathcal{X} \times \mathcal{X}$. The optimal rank-$R$ approximation to the kernel matrix $K$ retains the leading $R$ eigenpairs. The variational covariance $S = L_u L_u^\top$ is at best rank $R$, so it can only represent the posterior covariance faithfully up to this truncation error. Concretely, $R$ must satisfy

$$\frac{\sum_{j=1}^{R} \lambda_j}{\sum_{j=1}^{\infty} \lambda_j} \ge p, \tag{7}$$

for a desired variance-explained threshold $p$ (e.g. $p = 0.99$).

**Eigenvalue approximation via the power spectral density.** For a stationary kernel on a domain of width $L$, the eigenvalues of the integral operator $T$ can be approximated directly from the kernel's power spectral density (PSD) $S(\omega)$. Specifically, on the domain $[-L/2, L/2]$ the natural basis functions are the Fourier modes at frequencies $\omega_j = \pi j/L$, $j = 0, 1, 2, \ldots$, and the corresponding eigenvalues satisfy

$$\lambda_j \;\approx\; S(\omega_j) \;=\; S\left(\frac{\pi j}{L}\right). \tag{8}$$

This approximation becomes exact as $L \to \infty$ (Bochner's theorem) and is accurate whenever $L \gg l$. It converts the abstract eigenvalue problem into a concrete evaluation of the kernel's spectrum, enabling analytic reasoning about rank without forming $K$ explicitly.

For the two kernels most commonly used in practice:

$$\text{RBF:} \quad S(\omega) \propto \exp\left(-\tfrac{1}{2}\omega^2 l^2\right) \quad \text{(Gaussian decay)},$$
$$\text{Matérn-}\tfrac{3}{2}\text{:} \quad S(\omega) \propto \left(\tfrac{3}{l^2} + \omega^2\right)^{-2} \quad \text{(polynomial decay)}.$$

Substituting into (7), the cumulative variance explained by the first $R$ modes is

$$\frac{\sum_{j=0}^{R} S(\pi j/L)}{\sum_{j=0}^{\infty} S(\pi j/L)} \geq p,$$

which can be evaluated in closed form or by a fast one-dimensional sum, with no kernel matrix construction required.

**Closed-form scaling law.** We estimate $R$ directly from the Matérn-$\frac{3}{2}$ PSD evaluated on a Fourier grid, without forming $K$ explicitly. Extending the PSD to $d$ dimensions, the eigenvalues of the kernel integral operator are approximated as

$$\lambda_r \propto \left(\frac{3}{l^2} + \sum_{i=1}^{d} \omega_i^2\right)^{-(\nu+d/2)}, \qquad \omega_i = \frac{\pi j_i}{L}, \quad \nu = \tfrac{3}{2},$$

where $r$ indexes the eigenvalues sorted in descending order, and the exponent evaluates to 2 for $d = 1$ and $\frac{5}{2}$ for $d = 2$. The effective rank is then

$$R = \min\left\{ r \;\middle|\; \frac{\sum_{i=1}^{r} \lambda_i}{\sum_i \lambda_i} \geq p \right\}.$$

We recommend $p = 0.90$ as the default. When $L/l$ is small, $R \ll M$ and the low-rank parameterization offers a genuine computational saving. When $L/l$ is large, $R$ approaches $M$ and one should instead add a diagonal nugget $S = L_u L_u^\top + \delta I$ rather than increasing $R$ further.

### S7 Neighbor Selection

For each inducing point $j$, the neighbor set $n(j)$ is constructed by sampling $K$ points without replacement from the remaining $M - 1$ inducing locations, with inclusion probabilities proportional to the kernel evaluated between the query point and each candidate:

$$\Pr(i \in n(j)) \;\propto\; k(\mathbf{z}_j, \mathbf{z}_i), \qquad i \neq j,$$

where $k$ is the same kernel used in the GP prior (e.g. RBF or Matérn-$\frac{3}{2}$) with lengthscale $\ell$. Points within a few lengthscales of the query receive most of the probability mass, so the effective neighborhood radius is governed by $\ell$ rather than by data density. This contrasts with deterministic $K$-nearest-neighbor selection, where the spatial extent of $n(j)$ shrinks in dense regions and expands in sparse ones.

Concretely, the normalized sampling weights are

$$w_i^{(j)} = \frac{k(\mathbf{z}_j, \mathbf{z}_i)}{\sum_{i' \neq j} k(\mathbf{z}_j, \mathbf{z}_{i'})},$$

and $n(j)$ is drawn as a size-$K$ subset from $\{1, \ldots, M\} \setminus \{j\}$ with probabilities $\{w_i^{(j)}\}$. Because the kernel already encodes the correlation structure of the prior, this procedure preferentially includes the inducing points that most inform the conditional $p(U_j \mid U_{n(j)})$, improving the fidelity of the local approximation at fixed $K$.

### S8 LCGP Robustness

A key design choice in LCGP is how neighbors are selected for each inducing point. Standard approaches such as VNNGP [9] use deterministic Euclidean $K$-nearest neighbors, but LCGP instead conditions neighbor selection on the kernel: neighbors are sampled with probabilities proportional to the kernel similarity. Because the kernel already encodes the correlation structure of the prior, this procedure preferentially selects inducing points whose spatial and group relationships meaningfully inform the conditional $p(U_j \mid U_{n(j)})$, preserving cross-cell and cross-group correlations within each neighborhood. As demonstrated in S36 Fig, this kernel-conditioned selection also yields neighborhoods whose spatial extent remains stable as data density grows, whereas the Euclidean KNN radius collapses in dense regions—a critical advantage for ST data where cell density varies dramatically across spatial domains.

### S9 Empirical comparison: kernel-conditioned vs. Euclidean-KNN neighbor selection

The robustness argument in §S8 predicts that kernel-conditioned neighbor sampling and Euclidean $K$-nearest-neighbor (KNN) selection should produce different downstream conditional posteriors when the kernel encodes group structure that Euclidean distance does not respect. Here we show this empirically on Slide-seqV2 mouse hippocampus by holding everything else fixed.

We fit two MGGP-LCGP variants on the same 41,783-cell tissue with the same kernel, the same $K = 50$, the same $M = N$, the same loadings parameterization, and an identical training schedule. The only difference is the neighbor-selection rule: one variant draws the $K$ neighbors from a Euclidean KNN graph (the VNNGP [9] construction), the other draws them probabilistically from the MGGP kernel weights (§S7). To isolate the neighbor-selection contribution from the group-coupling contribution, we set the MGGP group-distance parameter $a = 10^6$ in both runs, which drives the cross-group covariance below numerical precision and turns the MGGP into a set of effectively independent per-group GPs. The remaining difference between the two variants is then purely the neighbor-selection rule.

S38 Fig shows where the difference between the two methods actually appears, and where it does not. The unconditional factor maps (top row of each block) are broadly similar across methods: the kernel and the data are the same, so the marginal posterior recovered by an LCGP-style sparse approximation does not depend much on whether neighbors are sampled by Euclidean KNN or by kernel weight. "Similar" rather than identical because nonnegative matrix factorization is itself non-identifiable—factor identity can permute across runs and rotations within the nonnegative cone can produce different but equivalent factorizations of the same data; once factors are matched across the two runs by gene-loading Pearson correlation, the matched factor maps trace the same anatomical structure. The methods diverge in the *conditional* groupwise posteriors below.

Under KNN neighbor selection (left block, rows below the unconditional row), the cell-type-conditional posterior is built from a fixed Euclidean-KNN neighborhood around each query cell. Many of those neighbors are not the queried cell type, so the conditional signal collapses onto small, locally connected patches of the queried cells—the conditional map looks like a scattering of small clusters wherever a tight pocket of the queried cell type happens to sit. The conditional posterior fails to extend to other regions of the tissue that contain the queried cell type but are not in the query's Euclidean neighborhood, and the resulting map has no contiguous spatial structure beyond those small clusters.

Under probabilistic (LCGP) selection (right block), neighbors are drawn from the MGGP kernel, which weights candidate cells by group similarity in addition to spatial proximity. The conditional posterior can therefore borrow strength from same-cell-type cells that remain highly weighted under the MGGP kernel and spatial lengthscale, including cells outside the immediate Euclidean KNN neighborhood; the resulting maps are more spatially coherent and trace recognizable anatomical structures (oligodendrocyte white-matter tracts, astrocyte distribution). Quantitatively, with $K = 50$ and the same training budget, probabilistic selection lifts mean Moran's $I$ across the ten factors from 0.705 (KNN) to 0.717 (probabilistic) and changes the reconstruction error by less than 0.001—modest gaps because both methods recover essentially the same marginal posterior. The qualitative gain is concentrated in the conditional posteriors, which is the inference type that LCGP explicitly motivates and that the smNSF analyses in the main paper rely on. This mirrors the pattern Wu et al. [9] reported for VNNGP on standard regression tasks: KNN is competitive when the kernel is single-group and Euclidean is the right notion of proximity, but degrades whenever the kernel encodes group structure that Euclidean distance does not respect.

### S10 Diagonal Variational Covariance Underestimates Posterior Variance

We examine the effect of constraining the variational covariance to be diagonal. For clarity we work in the unwhitened parameterization here; the conclusions carry over to the whitened parameterization used in other sections. To isolate the diagonal constraint, we analyze the simpler case $X = Z$—the LCGP regime where inducing points coincide with data locations—which provides a worst-case probe of the underestimation caused by disallowing off-diagonal structure.

Under $X = Z$, the SVGP-implied marginal at the data locations satisfies $q(f) = \mathcal{N}(m_u, S_u)$. For a Gaussian likelihood $p(y \mid f) = \mathcal{N}(f, \sigma^2 I)$, the $S_u$-dependent part of the ELBO is

$$\mathcal{L}(S_u) = \tfrac{1}{2}\log|S_u| - \tfrac{1}{2}\operatorname{tr}(K_{uu}^{-1}S_u) - \tfrac{1}{2\sigma^2}\operatorname{tr}(S_u).$$

The unconstrained optimum, derived in [3], is $S_u^\star = (K_{uu}^{-1} + \sigma^{-2}I)^{-1}$.

**Diagonal-constrained optimum.** Constraining $S_u = \operatorname{diag}(s_1, \ldots, s_M)$ makes the objective separable:

$$\mathcal{L}(S_u) = \sum_{i=1}^{M}\left[\tfrac{1}{2}\log s_i - \tfrac{1}{2}(K_{uu}^{-1})_{ii}s_i - \tfrac{1}{2\sigma^2}s_i\right] + \text{const}.$$

Setting $\partial\mathcal{L}/\partial s_i = 0$ yields the coordinatewise solution

$$s_i^\star = \frac{1}{(K_{uu}^{-1})_{ii} + \sigma^{-2}}, \qquad i = 1, \ldots, M.$$

**Diagonal systematically underestimates marginal variances.** Define $A := K_{uu}^{-1} + \sigma^{-2}I \succ 0$. The full optimum gives $(S_u^\star)_{ii} = (A^{-1})_{ii}$, while the diagonal optimum gives $s_i^\star = 1/A_{ii}$. For any symmetric positive-definite $A$,

$$(A^{-1})_{ii} \geq \frac{1}{A_{ii}},$$

with strict inequality whenever $A$ has nonzero off-diagonal entries. This follows from the Schur complement: writing $A$ in block form with the $i$-th coordinate partitioned out,

$$(A^{-1})_{ii} = \frac{1}{A_{ii} - A_{i,-i}A_{-i,-i}^{-1}A_{-i,i}} \geq \frac{1}{A_{ii}},$$

since $A_{i,-i}A_{-i,-i}^{-1}A_{-i,i} \geq 0$ for $A \succ 0$. Thus the diagonal variational solution elementwise underestimates every marginal posterior variance whenever the prior couples different inducing points.

**Jitter-induced variance collapse.** In practice a small jitter $\varepsilon > 0$ is added to the kernel: $K_{uu} \leftarrow K_{uu} + \varepsilon I$. Let $K_{uu} = \sum_{j=1}^{M}(\lambda_j + \varepsilon)u_j u_j^\top$ be the eigendecomposition, so that $(K_{uu}^{-1})_{ii} = \sum_{j=1}^{M} u_{ij}^2/(\lambda_j + \varepsilon)$ with $\sum_j u_{ij}^2 = 1$. If $K_{uu}$ has small eigenvalues (as occurs when the kernel lengthscale is long relative to the domain width—a common regime in ST data where coordinates are normalized to unit scale), then for any coordinate $i$ with nonzero weight on near-null directions,

$$(K_{uu}^{-1})_{ii} \approx \frac{c_i}{\varepsilon} \quad \Longrightarrow \quad s_i^\star \approx c_i^{-1}\varepsilon \downarrow 0 \quad \text{as } \varepsilon \to 0.$$

The mechanism is: jitter prevents $K_{uu}^{-1}$ from blowing up but inflates its diagonal entries whenever $K_{uu}$ has near-zero eigenvalues; the diagonal-only solution takes the reciprocal of these inflated values and collapses to $O(\varepsilon)$. The full-covariance optimum remains at $O(1)$ regardless of jitter.

Taken together, these results show that diagonal variational covariances systematically underestimate posterior marginal variances and, under standard jittering of $K_{uu}$, can drive them to zero irrespective of the data. This motivates using a low-rank plus diagonal parameterization for $S_u$, as described in §Parameterizing the Variational Covariance: the low-rank factor retains the dominant between-inducing-point correlations that diagonal approaches forfeit, while the diagonal component guards against the collapse documented above. For empirical evidence in spatial GP models, see [9].

### S11 Useful Identities

In this section we display Gaussian marginalization and the computation of the KL divergence of two Gaussian distributions, both derived in Damianou et al. 2015 [10] and Townes et al. 2023 [2] respectively.

#### S11.1 Gaussian Marginalization

We start by defining the following distributions:

$$p(f|u) = \mathcal{N}(f|Wu, \Sigma_f)$$

$$p(u) = \mathcal{N}(u|m_u, \Sigma_u)$$

The marginal distribution becomes the following:

$$p(f) = \mathcal{N}(f|Wm_u, \Sigma_f + W\Sigma_u W^T)$$

##### S11.2 KL divergence

We define the following distributions, where $u \in \mathbb{R}^N$:

$$p(u) = \mathcal{N}(u|\mu, \Sigma)$$
$$q(u) = \mathcal{N}(u|m, S)$$

The closed form of the KL divergence over $q(u)$, becomes:

$$\mathrm{KL}\left(q(\mathbf{u}) \parallel p(\mathbf{u})\right) = \frac{1}{2}\left[\log\frac{|\Sigma|}{|S|} - N + \mathrm{tr}\left\{\Sigma^{-1}S\right\} + (m-\mu)'\Sigma^{-1}(m-\mu)\right]$$

### S12 Inference and Training in smNSF

We describe the inference and training procedures that make smNSF applicable to large-scale ST datasets. The derivations below use only the per-data-point marginal moments $q(f_{i\ell}) = \mathcal{N}(\mu_{i\ell}, \sigma^2_{i\ell})$ obtained from the SVGP predictive form in §Variational inference for MGGPs; because these marginal moments are available for any multivariate normal variational distribution, the algebra below applies regardless of the specific parameterization chosen for $q(U)$.

**Expansion of the Poisson expected log-likelihood.** We start from the per-observation expected log-likelihood term that appears in the ELBO,

$$\mathcal{L}_1 = \sum_{j=1}^{D}\sum_{i=1}^{N} \mathbb{E}_{q(F_{i\cdot})}[\log p(Y_{ij} \mid F_{i\cdot})]\,,$$

and derive its explicit form for the smNSF observation model. For a Poisson likelihood with rate $\lambda_{ij} > 0$,

$$p(Y_{ij} \mid \lambda_{ij}) = \frac{\lambda_{ij}^{Y_{ij}}\, e^{-\lambda_{ij}}}{Y_{ij}!},$$

the log-likelihood is

$$\log p(Y_{ij} \mid \lambda_{ij}) = Y_{ij}\,\log\lambda_{ij} \;-\; \lambda_{ij} \;-\; \log(Y_{ij}!).$$

In the NSF parameterization the rate is a nonnegative-loadings linear combination of exponentiated factors,

$$\lambda_{ij} \;=\; \sum_{\ell=1}^{L} W_{j\ell}\, e^{F_{i\ell}}, \qquad W_{j\ell} \geq 0,$$

so the log-likelihood becomes

$$\log p(Y_{ij} \mid F_{i\cdot}) = Y_{ij}\,\log\left(\sum_{\ell} W_{j\ell}\, e^{F_{i\ell}}\right) \;-\; \sum_{\ell} W_{j\ell}\, e^{F_{i\ell}} \;-\; \log(Y_{ij}!).$$

Taking the expectation under $q(F_{i\cdot})$ and using linearity of expectation,

$$\mathbb{E}_{q(F_{i\cdot})}[\log p(Y_{ij} \mid F_{i\cdot})] = Y_{ij}\,\mathbb{E}_{q(F_{i\cdot})}\left[\log\sum_{\ell} W_{j\ell}\, e^{F_{i\ell}}\right] \;-\; \mathbb{E}_{q(F_{i\cdot})}\left[\sum_{\ell} W_{j\ell}\, e^{F_{i\ell}}\right] \;-\; \log(Y_{ij}!).$$

The mean-field factorization $q(F_{i\cdot}) = \prod_\ell q(F_{i\ell})$ established in §Posterior marginalization lets the second expectation pass through the sum,

$$\mathbb{E}_{q(F_{i\cdot})}\left[\sum_\ell W_{j\ell}\, e^{F_{i\ell}}\right] \;=\; \sum_\ell W_{j\ell}\, \mathbb{E}_{q(F_{i\ell})}\left[e^{F_{i\ell}}\right],$$

and since $F_{i\ell} \sim \mathcal{N}(\mu_{i\ell},\, \sigma_{i\ell}^2)$ under the variational posterior, the Gaussian moment-generating function gives the closed form

$$\mathbb{E}_{q(F_{i\ell})}\left[e^{F_{i\ell}}\right] \;=\; \exp\!\left(\mu_{i\ell} + \tfrac{1}{2}\sigma_{i\ell}^2\right).$$

Combining these pieces, the per-observation expected log-likelihood reduces to

$$\mathbb{E}_{q(F_{i\cdot})}[\log p(Y_{ij} \mid F_{i\cdot})] = Y_{ij}\, \mathbb{E}_{q(F_{i\cdot})}\left[\log \sum_\ell W_{j\ell}\, e^{F_{i\ell}}\right] - \sum_\ell W_{j\ell}\, \exp\!\left(\mu_{i\ell} + \tfrac{1}{2}\sigma_{i\ell}^2\right) - \log(Y_{ij}!).$$

The linear term and the constant $-\log(Y_{ij}!)$ are now in closed form; only the log-sum-exp expectation $\mathbb{E}_{q(F_{i\cdot})}[\log \sum_\ell W_{j\ell} e^{F_{i\ell}}]$ remains analytically intractable. We address it next.

**Jensen sandwich bounds for the log-sum-exp.** The log-sum-exp term $\mathbb{E}[\log \sum_\ell W_{j\ell} e^{F_{i\ell}}]$ has no closed form. Let $g(z) = \log \sum_m W_{jm} e^{z_m}$ (convex in $z$) and $S = \sum_m W_{jm} e^{F_{im}} > 0$. Applying Jensen's inequality to $g$ and to log gives a sandwich:

$$\boxed{\log \sum_m W_{jm}\, e^{\mathbb{E}[F_{im}]} \;\le\; \mathbb{E}\left[\log \sum_m W_{jm} e^{F_{im}}\right] \;\le\; \log \sum_m W_{jm}\, \mathbb{E}\left[e^{F_{im}}\right]}$$

(the left bound uses convexity of $g$; the right uses concavity of log: $\mathbb{E}[\log S] \le \log \mathbb{E}[S]$). For Gaussian marginals this becomes

$$\log \sum_m W_{jm}\, e^{\mu_{im}} \;\le\; \mathbb{E}\left[\log \sum_m W_{jm} e^{F_{im}}\right] \;\le\; \log \sum_m W_{jm}\, \exp\!\left(\mu_{im} + \tfrac{1}{2}\sigma_{im}^2\right).$$

**Multiplicative update rules for the loadings $W$.** The nonnegative loadings matrix $W$ is updated via multiplicative update rules (MUR) [11, 12]. The gradient of the ELBO with respect to $W_{j\ell}$ splits into a positive and a negative part, $\nabla_{W_{j\ell}}\mathcal{L} = [\nabla_{W_{j\ell}}\mathcal{L}]^+ - [\nabla_{W_{j\ell}}\mathcal{L}]^-$, and the MUR iteration is $W_{j\ell} \leftarrow W_{j\ell}\, [\nabla_{W_{j\ell}}\mathcal{L}]^+ \,/\, [\nabla_{W_{j\ell}}\mathcal{L}]^-$. Depending on how the log-sum-exp expectation is handled, three modes arise:

**Method 1: Jensen lower bound.** Using the left Jensen bound $\mathbb{E}[\log \sum_\ell W_{j\ell} e^{F_{i\ell}}] \ge \log \sum_\ell W_{j\ell} e^{\mu_{i\ell}}$ gives a fully analytic ELBO:

$$\mathcal{L}_{\mathrm{LB}} = \sum_{i,j} \left[Y_{ij} \log\left(\sum_\ell W_{j\ell} e^{\mu_{i\ell}}\right) - \sum_\ell W_{j\ell} e^{\mu_{i\ell} + \frac{1}{2}\sigma_{i\ell}^2}\right].$$

Let $\hat{R}_{ij}^{\mu} = \sum_k W_{jk} e^{\mu_{ik}}$. The MUR is

$$\boxed{W_{j\ell} \leftarrow W_{j\ell}\, \frac{\sum_i \frac{Y_{ij}}{\hat{R}_{ij}^{\mu}} e^{\mu_{i\ell}}}{\sum_i e^{\mu_{i\ell} + \frac{1}{2}\sigma_{i\ell}^2}}.}$$

The denominator includes the variance penalty, effectively regularizing factors with high uncertainty.

**Method 2: Expanded likelihood.** The linear term is analytic while the log-sum-exp is estimated via $S$ Monte Carlo samples $F_{i\ell}^{(s)} \sim q(F_{i\ell})$:

$$\mathcal{L}_{\text{Hyb}} = \sum_{i,j} \Big[ \tfrac{1}{S} \sum_{s=1}^{S} Y_{ij} \log \big( \sum_{\ell} W_{j\ell} e^{F_{i\ell}^{(s)}} \big) - \sum_{\ell} W_{j\ell} e^{\mu_{i\ell} + \frac{1}{2}\sigma_{i\ell}^2} \Big].$$

Let $R_{ij}^{(s)} = \sum_k W_{jk} e^{F_{ik}^{(s)}}$. The MUR is

$$\boxed{W_{j\ell} \leftarrow W_{j\ell} \, \frac{\frac{1}{S} \sum_{s=1}^{S} \sum_i \frac{Y_{ij}}{R_{ij}^{(s)}} e^{F_{i\ell}^{(s)}}}{\sum_i e^{\mu_{i\ell} + \frac{1}{2}\sigma_{i\ell}^2}}.}$$

This is the default ELBO mode used in all experiments in this work. The numerator averages gradients from $S$ noisy reconstructions while the denominator remains analytic, yielding lower Monte Carlo variance than Method 3 (see S35 Fig for an empirical comparison of all three methods).

**Method 3: Full Monte Carlo.** Both terms are estimated via samples:

$$\mathcal{L}_{\text{MC}} = \tfrac{1}{S} \sum_{s=1}^{S} \sum_{i,j} \Big[ Y_{ij} \log \big( \sum_{\ell} W_{j\ell} e^{F_{i\ell}^{(s)}} \big) - \sum_{\ell} W_{j\ell} e^{F_{i\ell}^{(s)}} \Big].$$

The MUR,

$$\boxed{W_{j\ell} \leftarrow W_{j\ell} \, \frac{\sum_{s=1}^{S} \sum_i \frac{Y_{ij}}{R_{ij}^{(s)}} e^{F_{i\ell}^{(s)}}}{\sum_{s=1}^{S} \sum_i e^{F_{i\ell}^{(s)}}},}$$

has both numerator and denominator subject to sampling noise and typically requires larger $S$ for stability. The expanded estimator (Method 2) offers the best trade-off between stability and final loss (S35 Fig).

**Mini-batch estimation.** For large ST datasets, evaluating the full ELBO over all $N \times D$ entries is impractical. Following the SVGP minibatching framework of [3, 5] and stochastic variational inference [13], we form an unbiased mini-batch estimator by subsampling rows and columns independently. Let $\mathcal{B}_N \subset \{1, \dots, N\}$ of size $N_b$ and $\mathcal{B}_D \subset \{1, \dots, D\}$ of size $D_b$ be drawn uniformly without replacement. The standard SVGP mini-batch estimator is

$$\hat{\mathcal{L}}_{\text{GP}} = \frac{ND}{N_b D_b} \sum_{i \in \mathcal{B}_N} \sum_{j \in \mathcal{B}_D} \mathbb{E}_{q(F_{i\ell})}[\log p(Y_{ij} \mid F_{i\cdot})] \; - \; \sum_{\ell=1}^{L} \text{KL}(q(U_\ell) \, \| \, p(U_\ell)) \, .$$

This estimator is unbiased: the $ND/(N_b D_b)$ rescaling on the likelihood term compensates for uniform subsampling of rows and columns, and the KL term does not depend on the mini-batch and is computed exactly.

**$N/M$ correction for the inducing-point KL.** The SVGP ELBO is itself a lower bound whose tightness depends on $M$. When $N \gg M$ (typical in ST applications where $N$ can reach $10^6$ cells with $M \approx 3{,}000$ inducing points), the KL penalty scales as $M \cdot L$ rather than $N \cdot L$, causing under-regularization of the latent factors. To see this, consider the limit $\ell \to 0$ where the kernel becomes diagonal: the GP prior degenerates to an i.i.d. prior $p(F_{i\ell}) = \mathcal{N}(0, \sigma^2)$, and the spatial ELBO should coincide with a non-spatial ELBO whose KL scales as $N \cdot L$. Yet the SVGP KL retains scaling $\propto M \cdot L$.

This is underweighted by a factor of $N/M$, and the likelihood term (scaling as $N \times D$) dominates the objective.

We therefore propose scaling the inducing-point KL by $N/M$:

$$\hat{\mathcal{L}}_{\mathrm{GP}}^{\mathrm{corr}} = \frac{ND}{N_b D_b} \sum_{i \in \mathcal{B}_N} \sum_{j \in \mathcal{B}_D} \mathbb{E}_{q(F_{i\ell})}[\log p(Y_{ij} \mid F_{i\cdot})] \; - \; \frac{N}{M} \sum_{\ell=1}^{L} \mathrm{KL}(q(U_\ell) \,\|\, p(U_\ell))\,.$$

In the LCGP regime where $M = N$, the correction factor $N/M = 1$ and the standard estimator $(ND/N_b D_b) \sum_{i \in \mathcal{B}_N} \sum_{j \in \mathcal{B}_D} \mathbb{E}[\log p(Y_{ij} \mid F_{i\cdot})] - \sum_\ell \mathrm{KL}(q(U_\ell) \| p(U_\ell))$ is already correctly scaled. For general $M < N$, each inducing point is responsible for approximately $N/M$ data locations; the correction restores the balance between likelihood and prior. Our experiments indicate that improper mini-batch scaling can slow convergence by 10–100× or yield uninformative, sparse factorizations, while the corrected estimator produces well-regularized factors across a wide range of spatial resolutions.

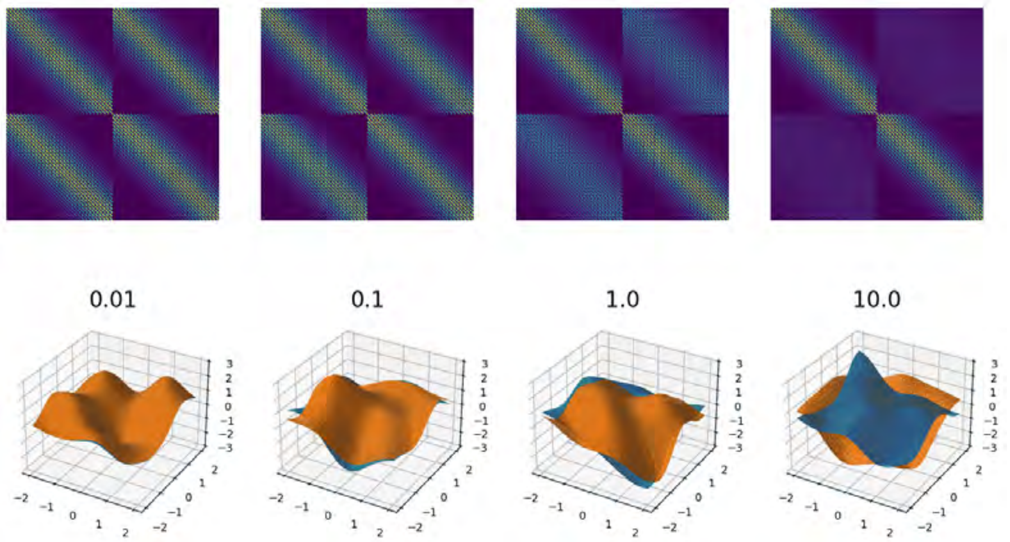


**Fig 1. MGGP prior illustration on a 2D toy dataset.** Samples drawn from the Matérn-3/2 MGGP prior across four group-kernel lengthscales ($a \in \{0.01, 0.1, 1.0, 10.0\}$) over four stripe-pattern ground-truth group configurations, visualized as 3D surfaces in both a smooth regime and a noisy regime. Illustrates how the group-difference parameter $a$ controls sharing between groups (cell types): small $a$ yields near-identical across-group factors; large $a$ yields group-specific factors.

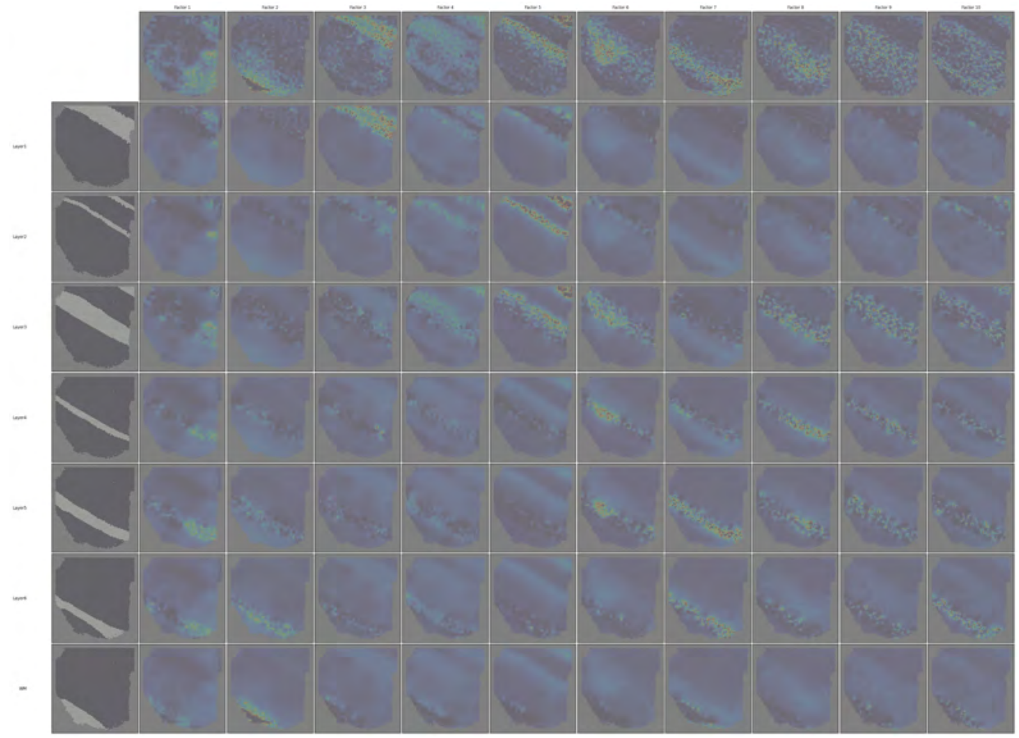


**Fig 2. MGGP-LCGP group-conditional spatial factors for DLPFC slice 151507.** Full grid of group-conditional spatial factors for Maynard et al. DLPFC slice 151507 (Visium; 7 annotated cortical layers).

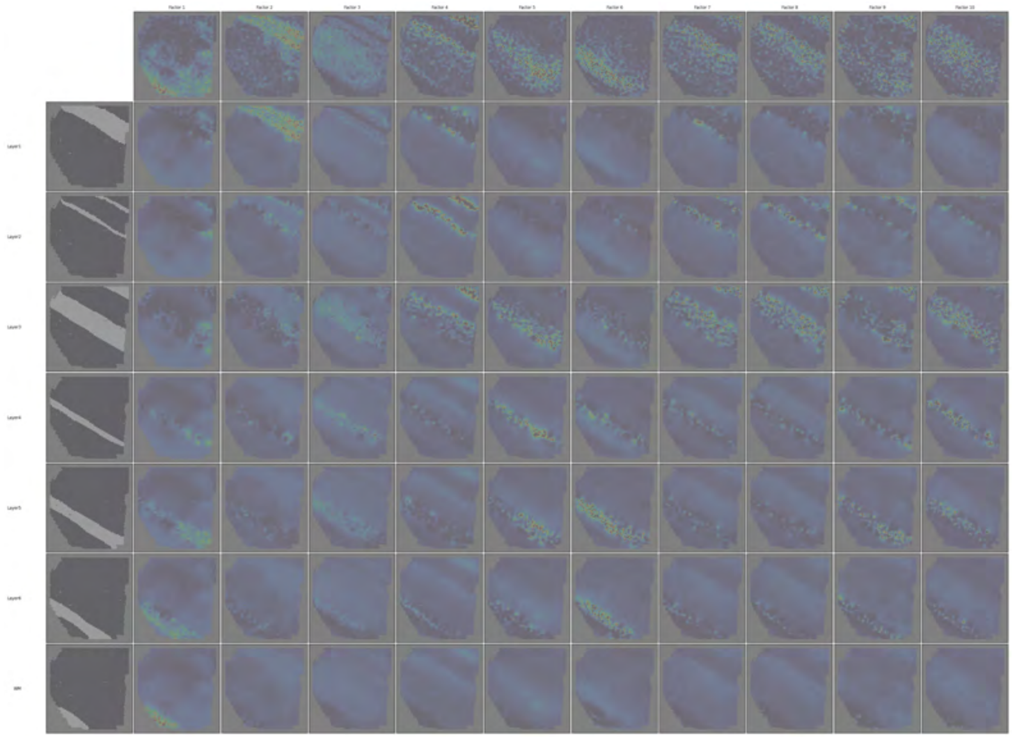


**Fig 3. MGGP-LCGP group-conditional spatial factors for DLPFC slice 151508.** Full grid of group-conditional spatial factors for Maynard et al. DLPFC slice 151508.

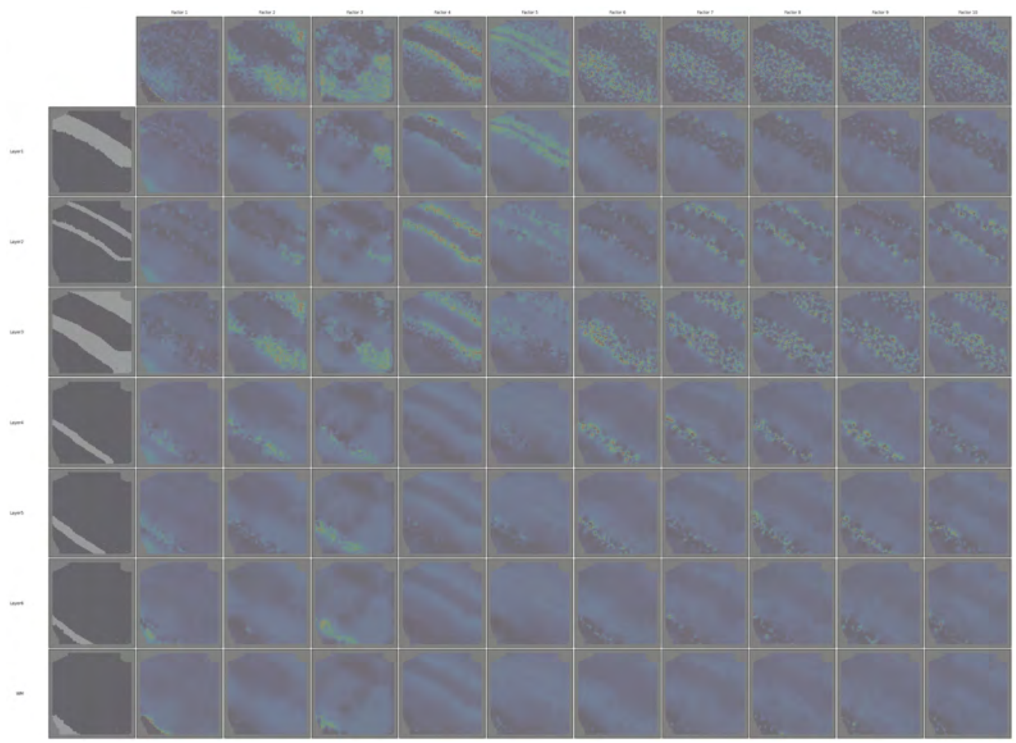

**Fig 4. MGGP-LCGP group-conditional spatial factors for DLPFC slice 151509.** Full grid of group-conditional spatial factors for Maynard et al. DLPFC slice 151509.

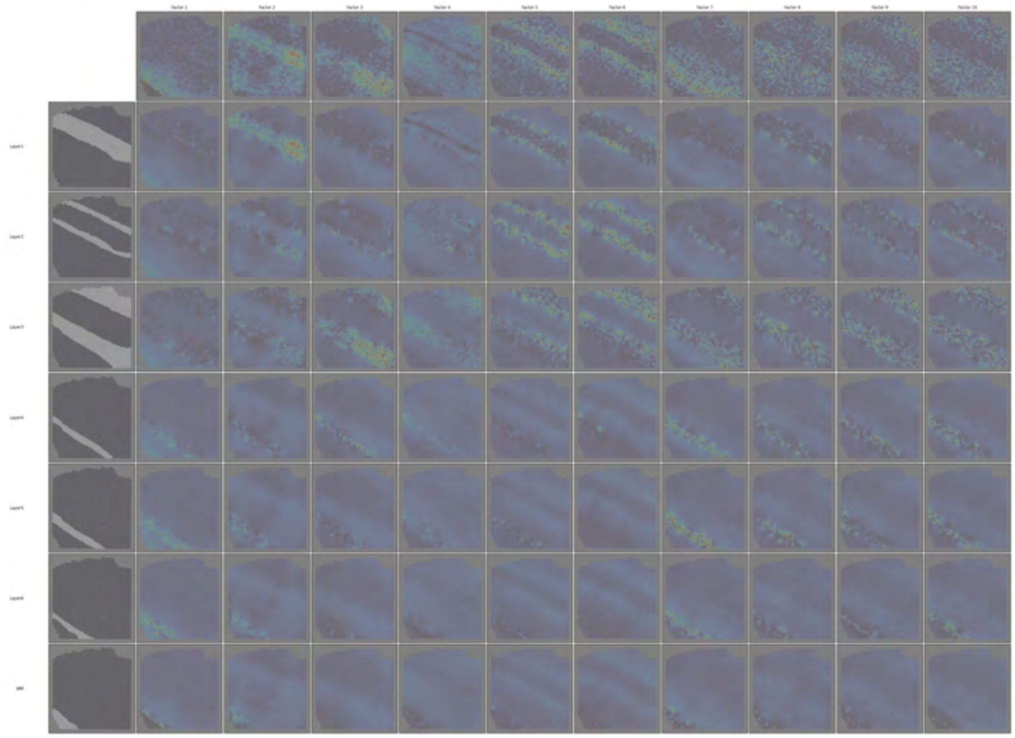


**Fig 5. MGGP-LCGP group-conditional spatial factors for DLPFC slice 151510.** Full grid of group-conditional spatial factors for Maynard et al. DLPFC slice 151510.

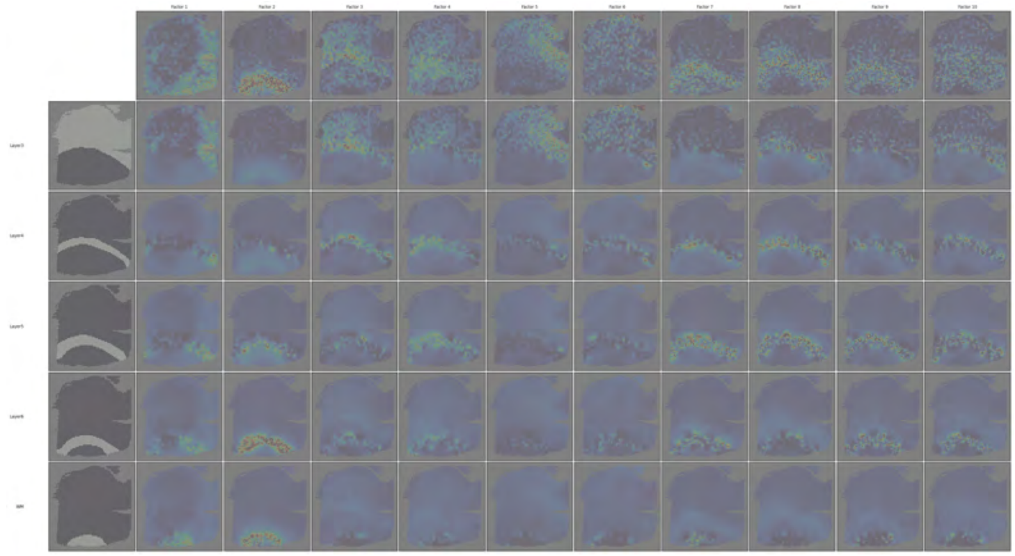

**Fig 6. MGGP-LCGP group-conditional spatial factors for DLPFC slice 151669.** Full grid of group-conditional spatial factors for Maynard et al. DLPFC slice 151669.

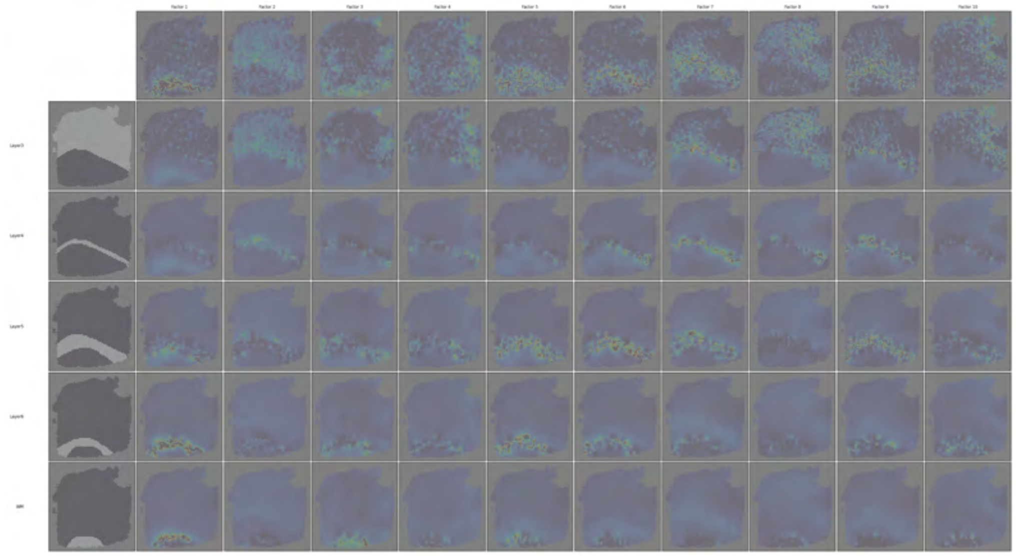

**Fig 7. MGGP-LCGP group-conditional spatial factors for DLPFC slice 151670.** Full grid of group-conditional spatial factors for Maynard et al. DLPFC slice 151670.

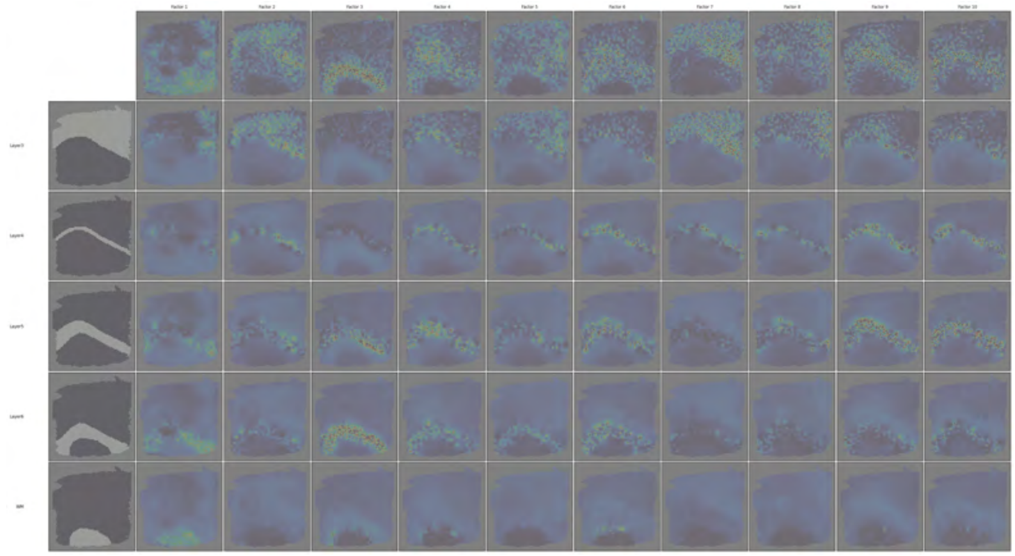

**Fig 8. MGGP-LCGP group-conditional spatial factors for DLPFC slice 151671.** Full grid of group-conditional spatial factors for Maynard et al. DLPFC slice 151671.

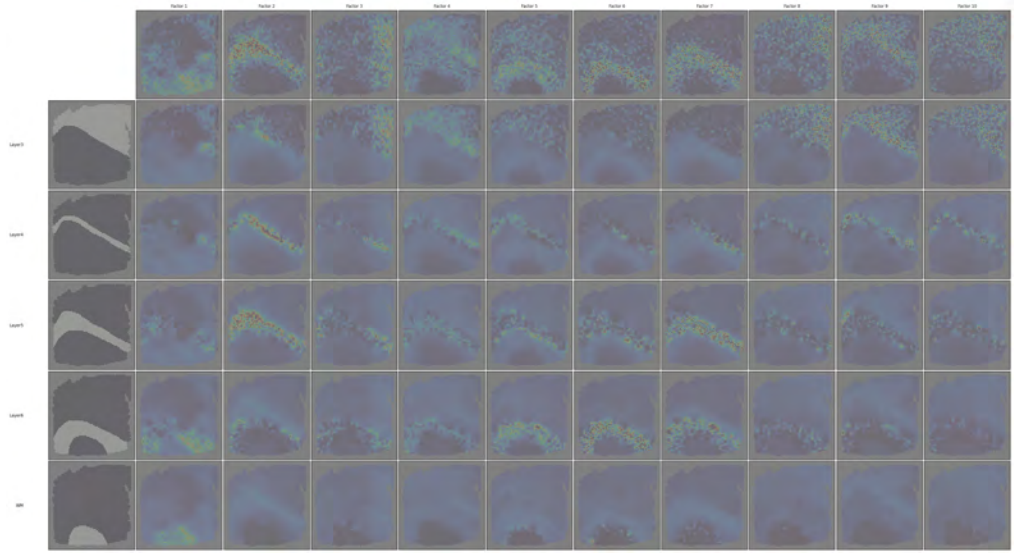

**Fig 9. MGGP-LCGP group-conditional spatial factors for DLPFC slice 151672.** Full grid of group-conditional spatial factors for Maynard et al. DLPFC slice 151672.

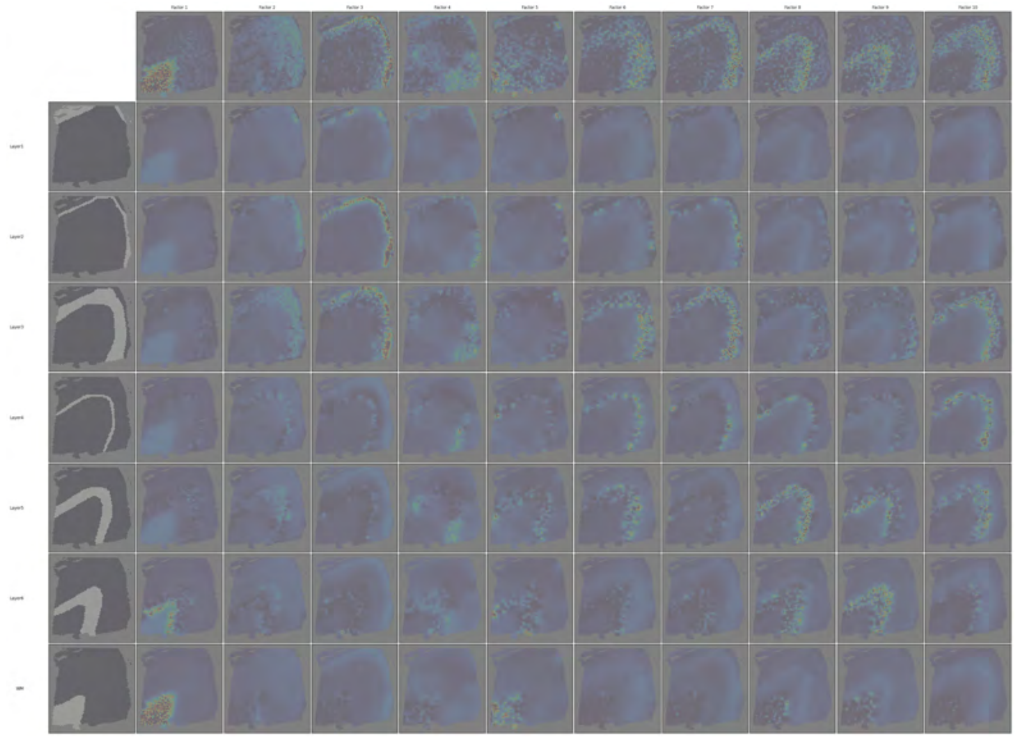


**Fig 10. MGGP-LCGP group-conditional spatial factors for DLPFC slice 151673.** Full grid of group-conditional spatial factors for Maynard et al. DLPFC slice 151673.

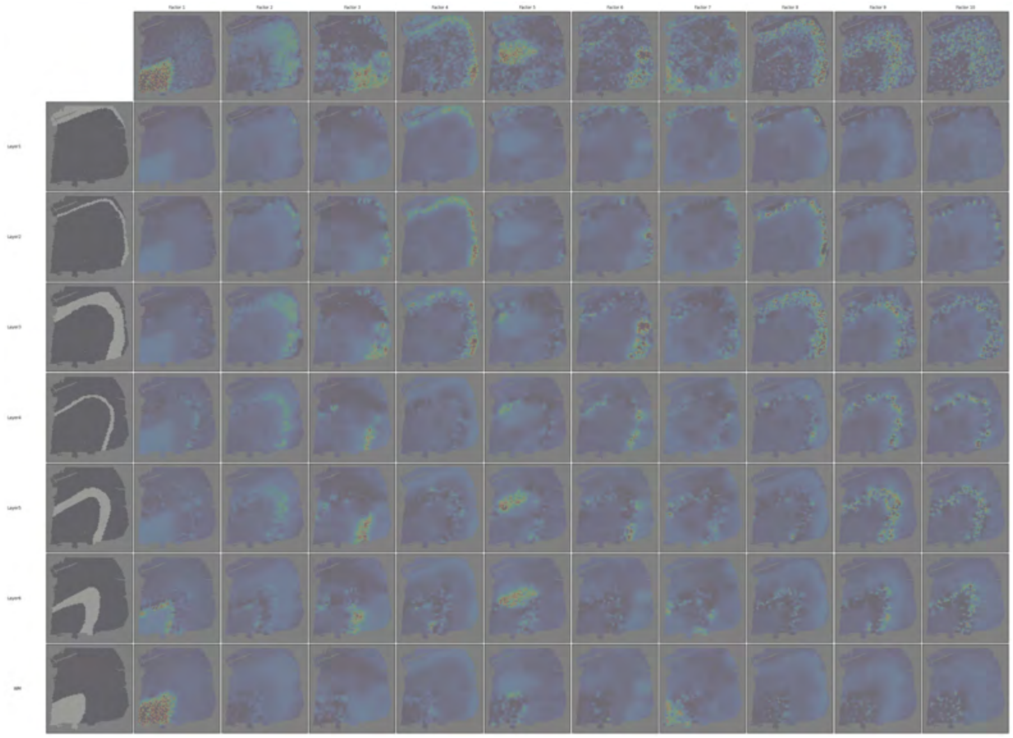

**Fig 11. MGGP-LCGP group-conditional spatial factors for DLPFC slice 151674.** Full grid of group-conditional spatial factors for Maynard et al. DLPFC slice 151674.

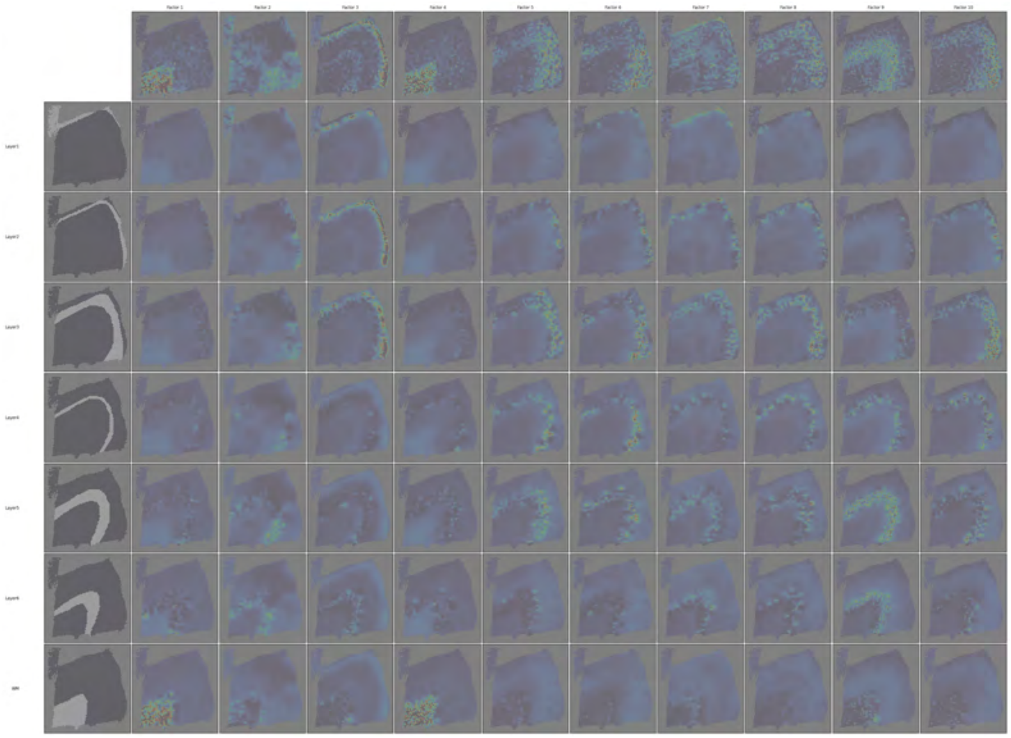

**Fig 12. MGGP-LCGP group-conditional spatial factors for DLPFC slice 151675.** Full grid of group-conditional spatial factors for Maynard et al. DLPFC slice 151675.

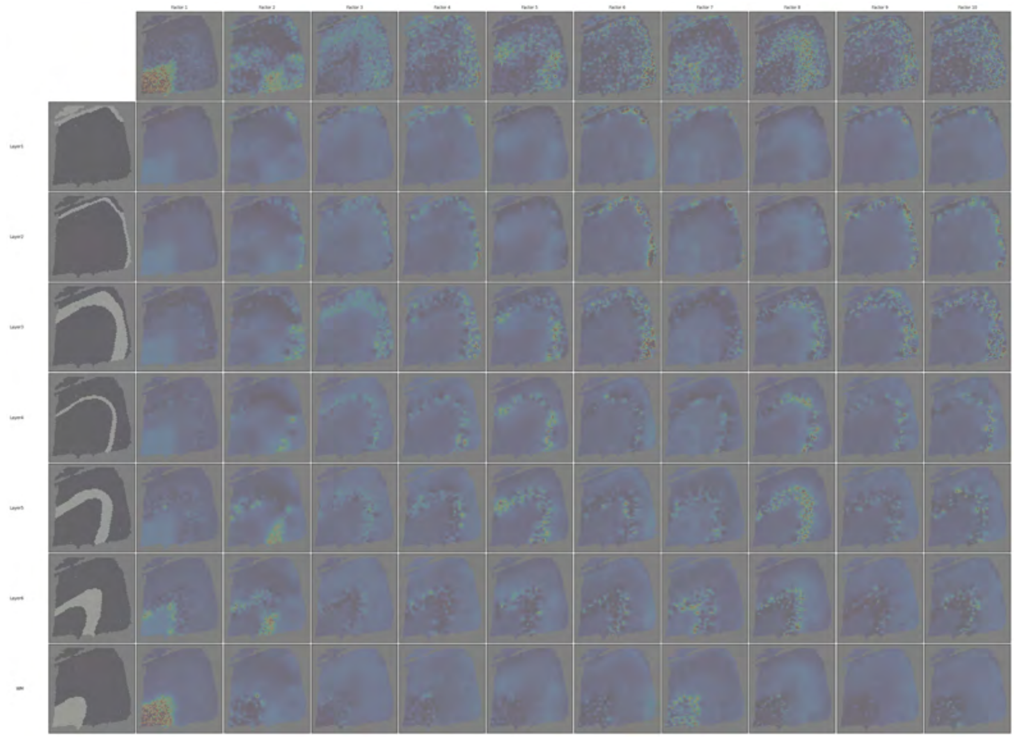

**Fig 13. MGGP-LCGP group-conditional spatial factors for DLPFC slice 151676.** Full grid of group-conditional spatial factors for Maynard et al. DLPFC slice 151676.

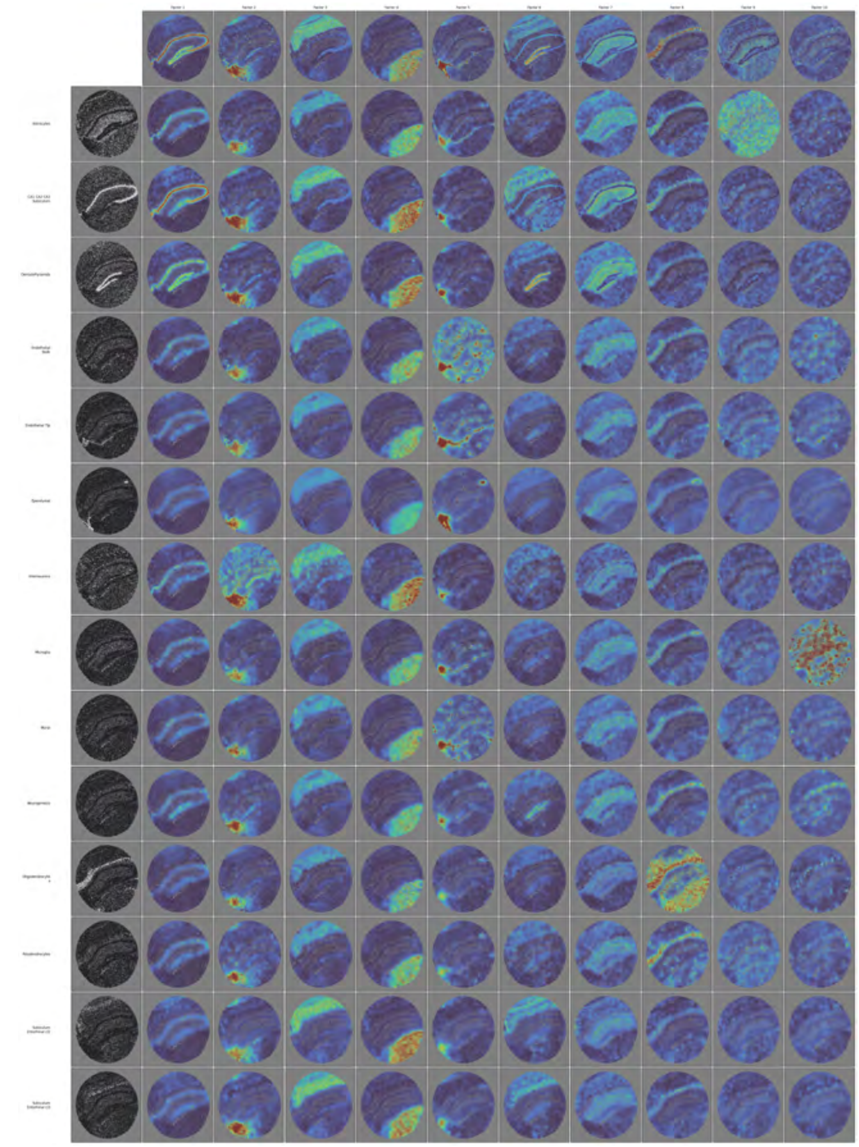

**Fig 14. MGGP-LCGP group-conditional spatial factors for Slide-seqV2 mouse hippocampus.** Full grid of all $L = 10$ smNSF spatial factors conditioned on each of the 14 annotated groups (10 cell types and 4 hippocampal subfield regions) in the Slide-seqV2 hippocampus dataset (41,783 spatial barcodes). Complements the curated subset shown in Fig 2 and the consolidated Slide-seq main-text panel.

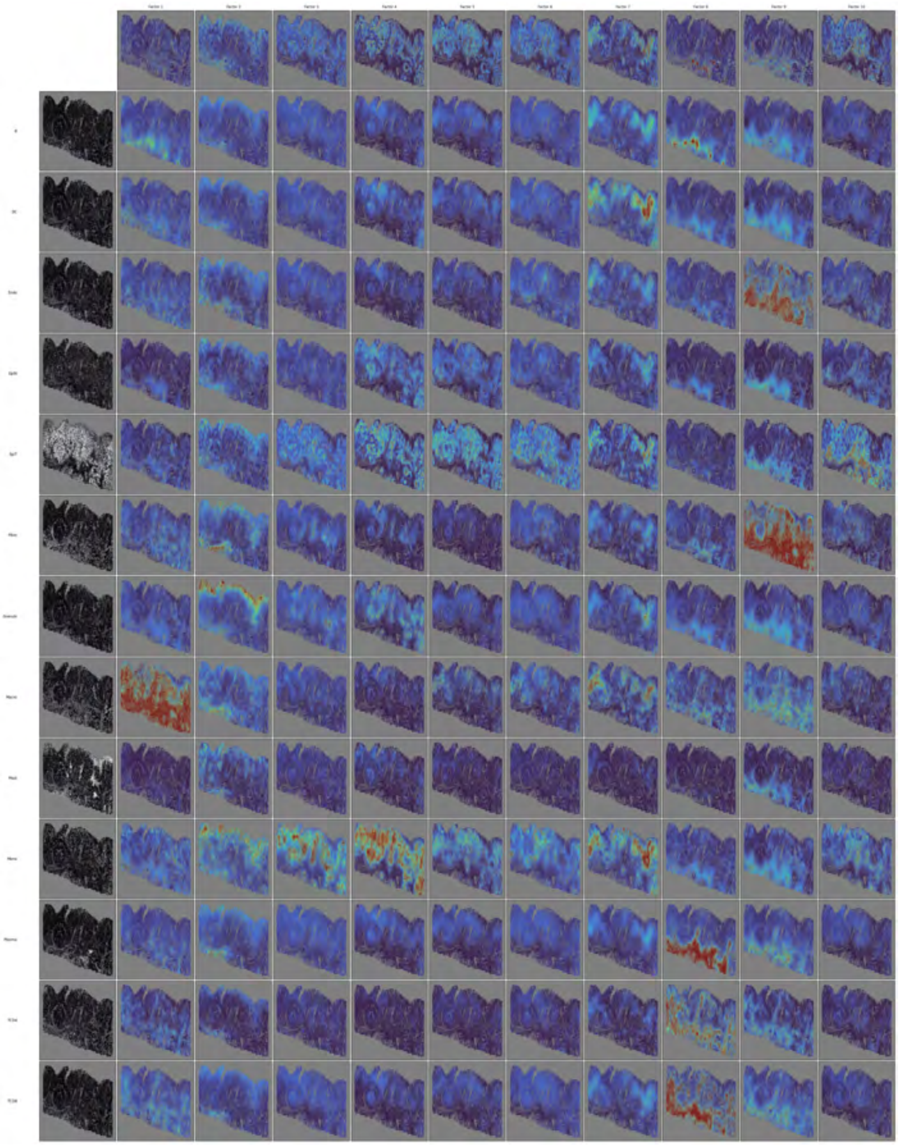

**Fig 15. MGGP-LCGP cell-type-conditional spatial factors for HuColonCa-FFPE MERFISH human colorectal cancer.** Full grid of cell-type-conditional spatial factors for the HuColonCa-FFPE MERFISH human colorectal cancer dataset (137,693 cells after 6× subsample, 13 cell-type groups collapsed from ∼50 fine annotations). Complements the curated subset shown in the colon main-text panel.

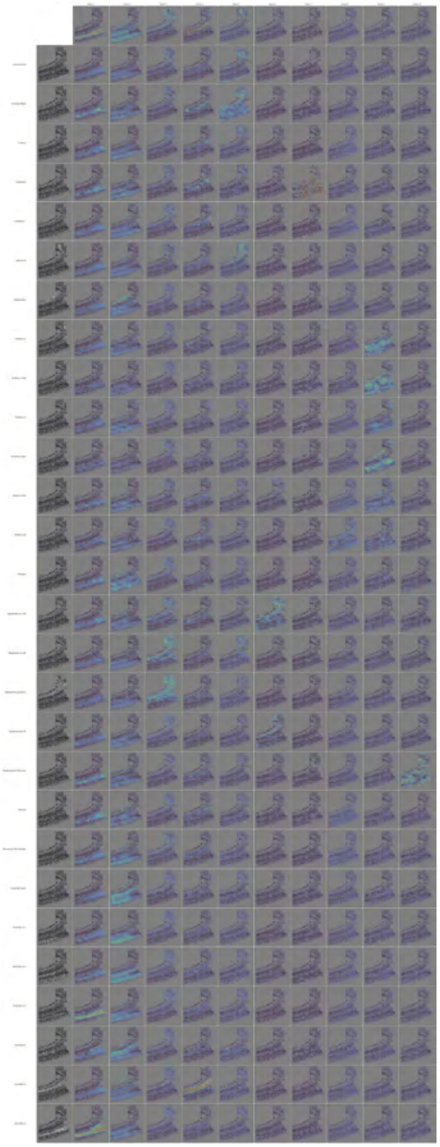

**Fig 16. MGGP-LCGP cell-type-conditional spatial factors for osmFISH mouse somatosensory cortex.** Full grid of cell-type-conditional spatial factors for the osmFISH mouse cortex dataset (4,727 cells, 33 genes, 28 cell types). Complements the curated subset shown in the osmFISH main-text panel.

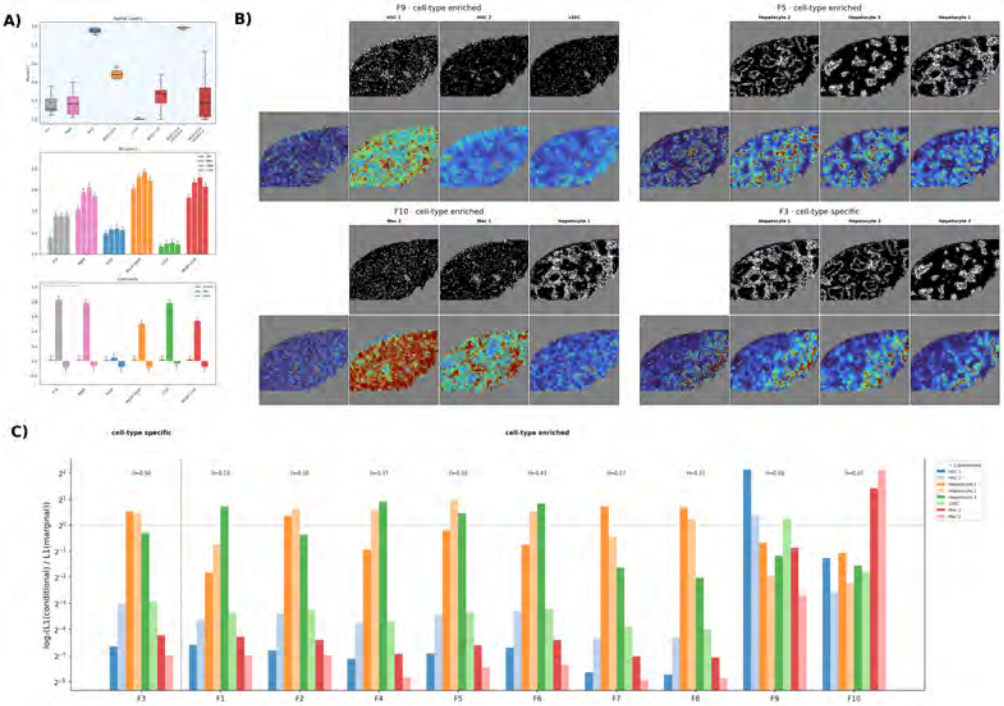


**Fig 17. smNSF analysis of MERFISH human liver — healthy sample AM042.** Per-sample panel for the healthy AM042 liver sample.

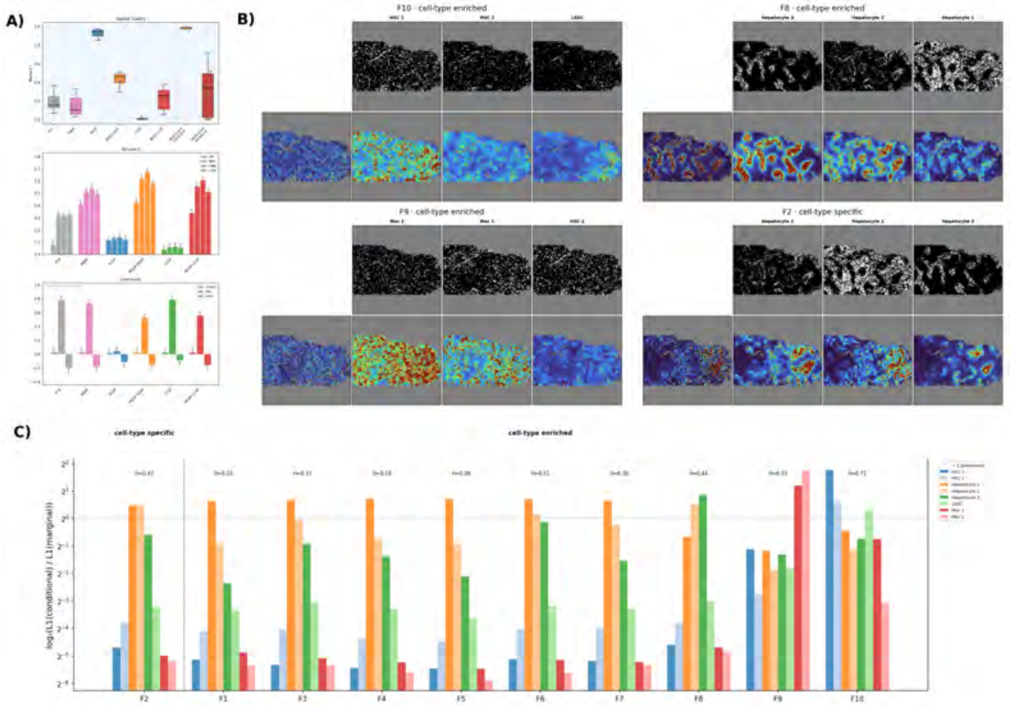


**Fig 18. smNSF analysis of MERFISH human liver — healthy sample AM061.** Per-sample panel for the healthy AM061 liver sample.

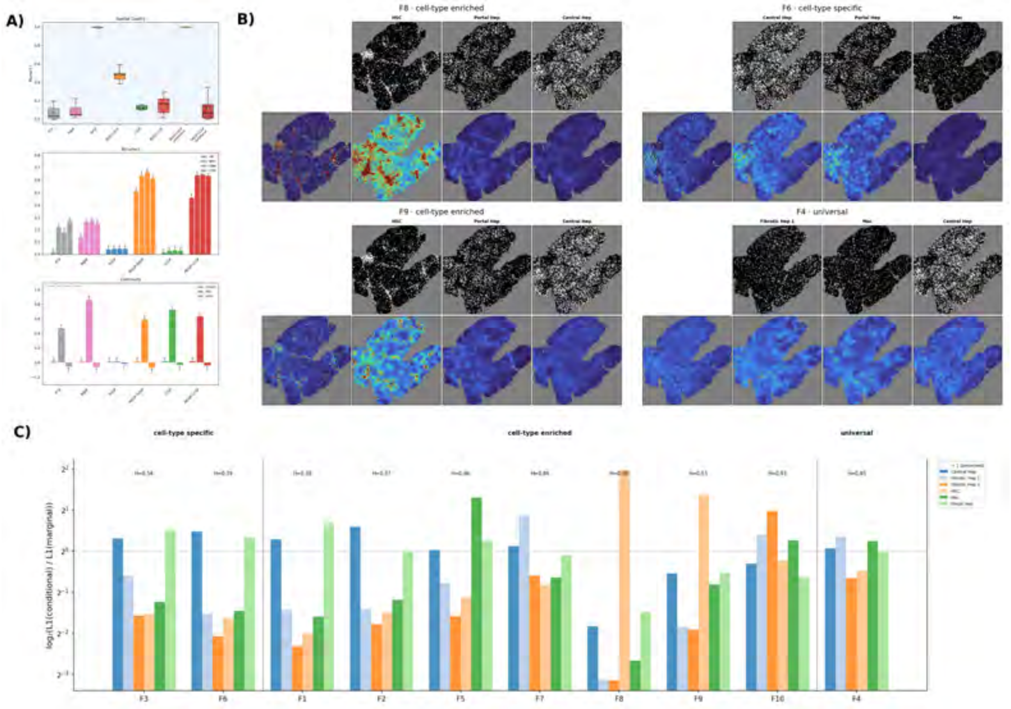


**Fig 19. smNSF analysis of MERFISH human liver — diseased sample AM031.** Per-sample panel for the diseased AM031 liver sample.

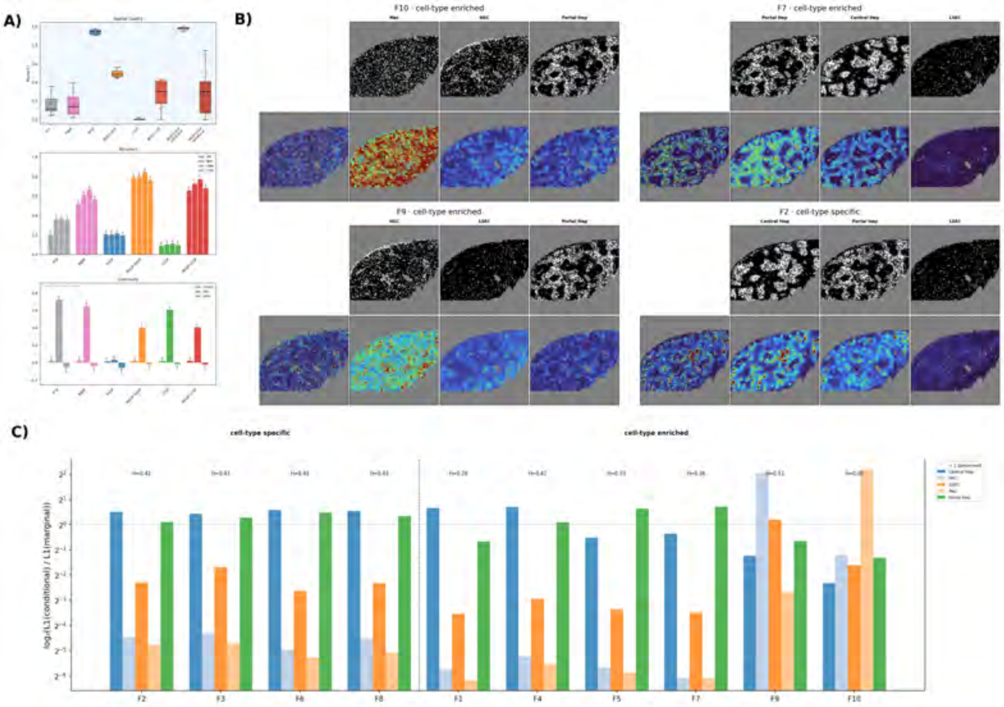


**Fig 20. smNSF analysis of MERFISH human liver — diseased sample AM042.** Per-sample panel for the diseased AM042 liver sample.

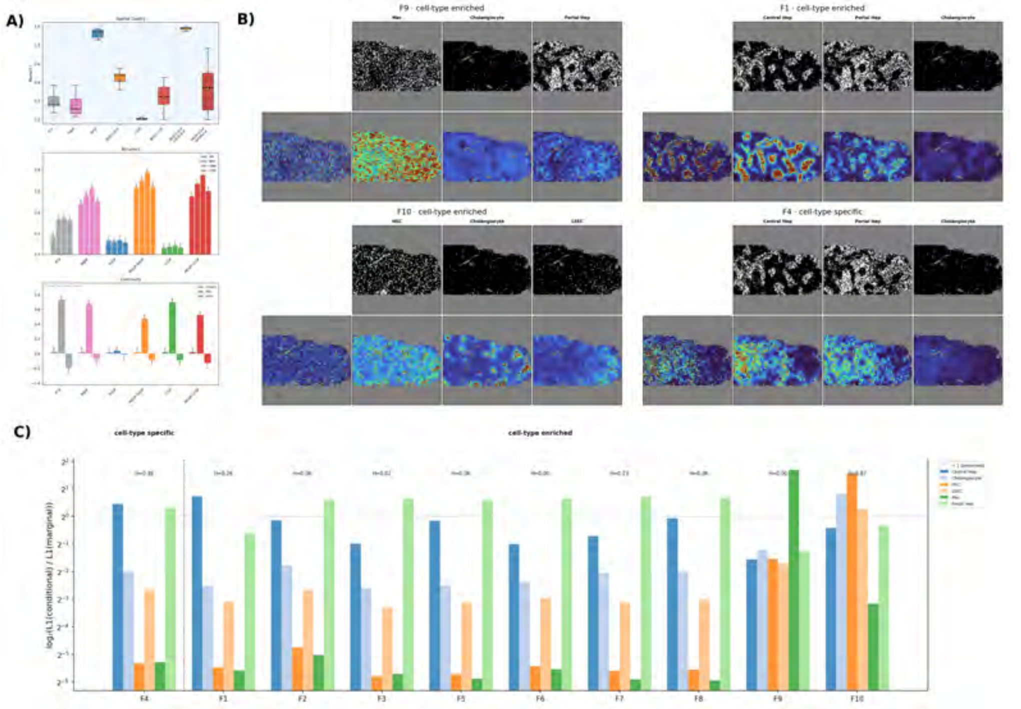


**Fig 21. smNSF analysis of MERFISH human liver — diseased sample AM061.** Per-sample panel for the diseased AM061 liver sample.

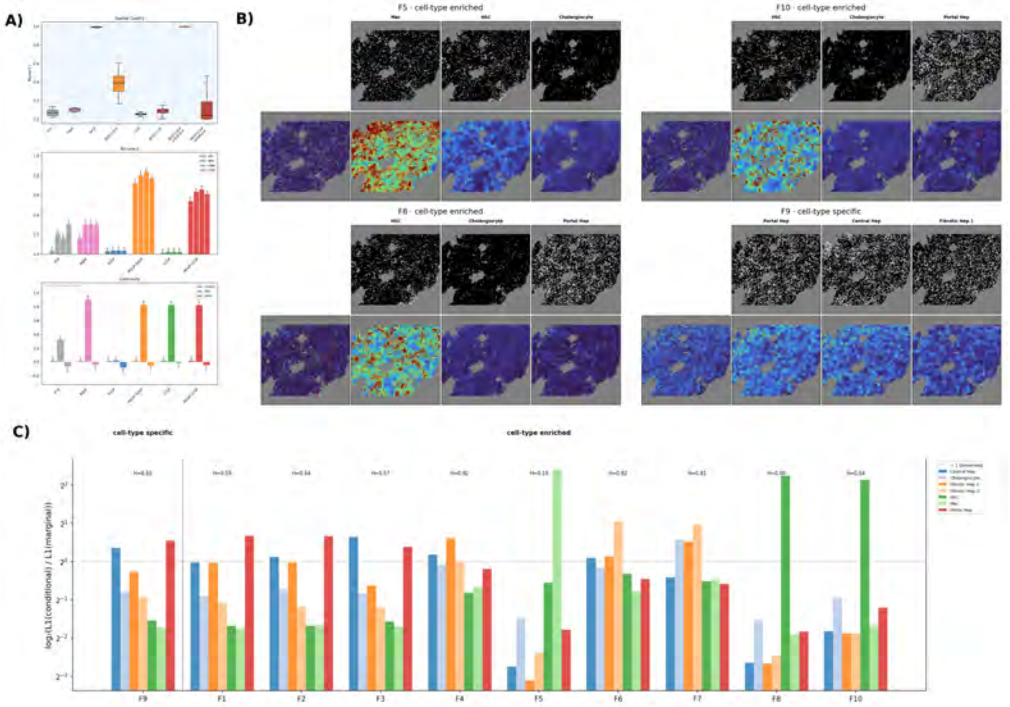

**Fig 22. smNSF analysis of MERFISH human liver — diseased sample AM062.** Per-sample panel for the diseased AM062 liver sample.

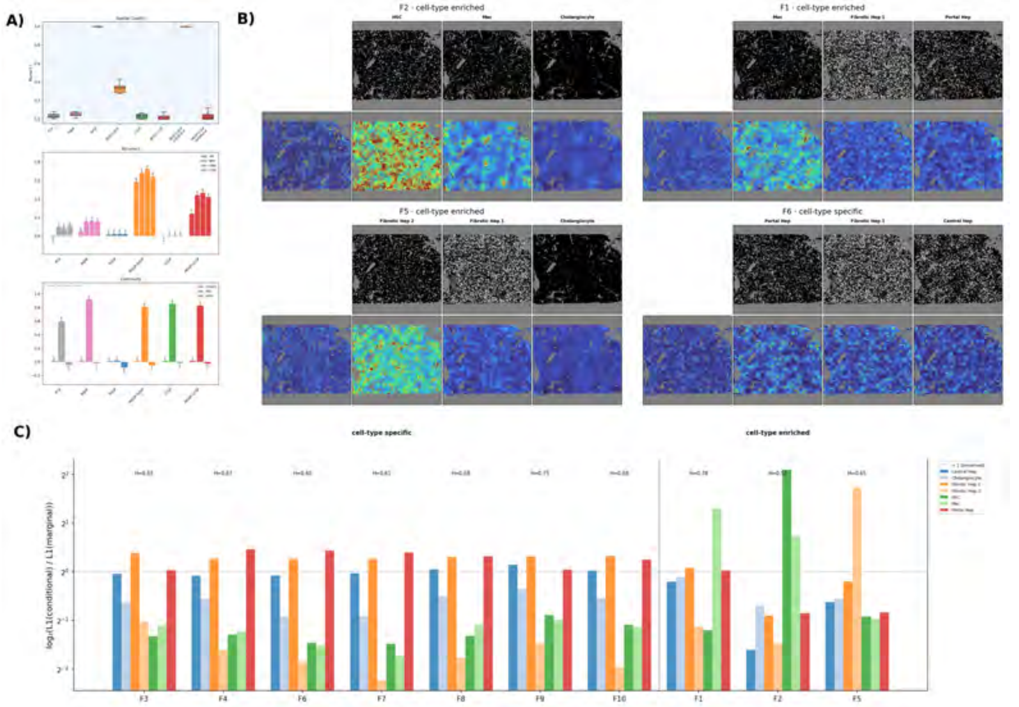

**Fig 23. smNSF analysis of MERFISH human liver — diseased sample AM072.** Per-sample panel for the diseased AM072 liver sample.

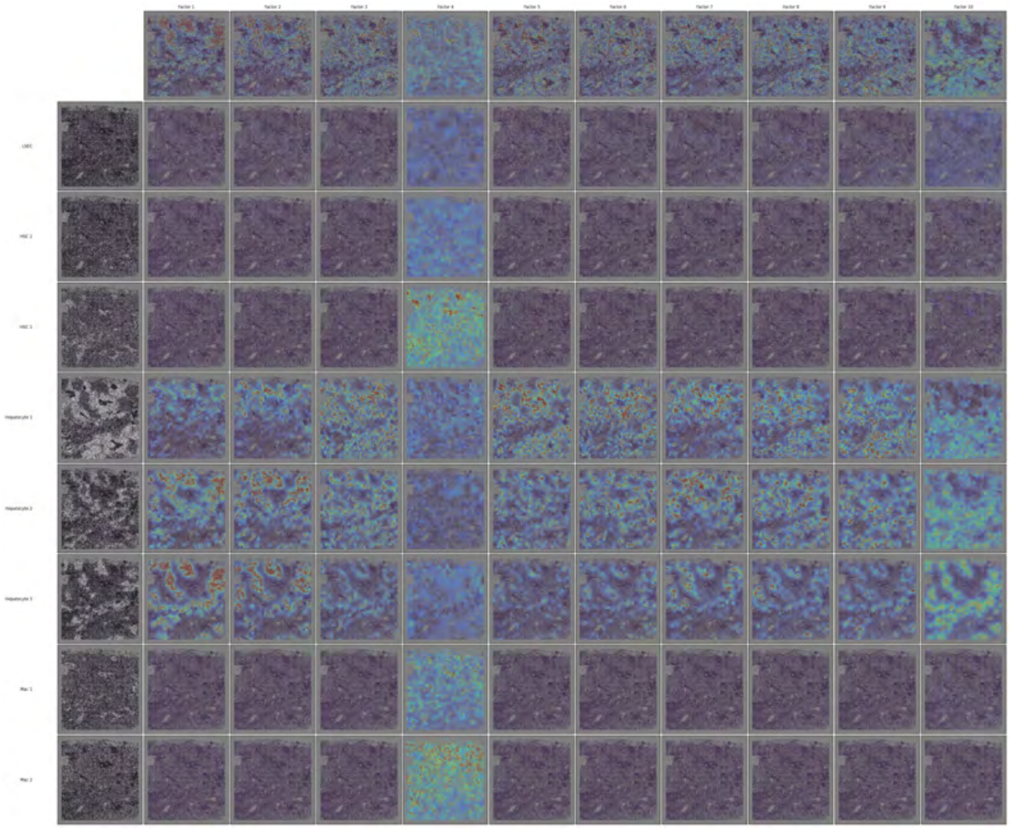

**Fig 24. MGGP-LCGP cell-type-conditional spatial factors for MERFISH human liver — healthy sample AM048.** Full grid of cell-type-conditional spatial factors for the healthy AM048 liver sample.

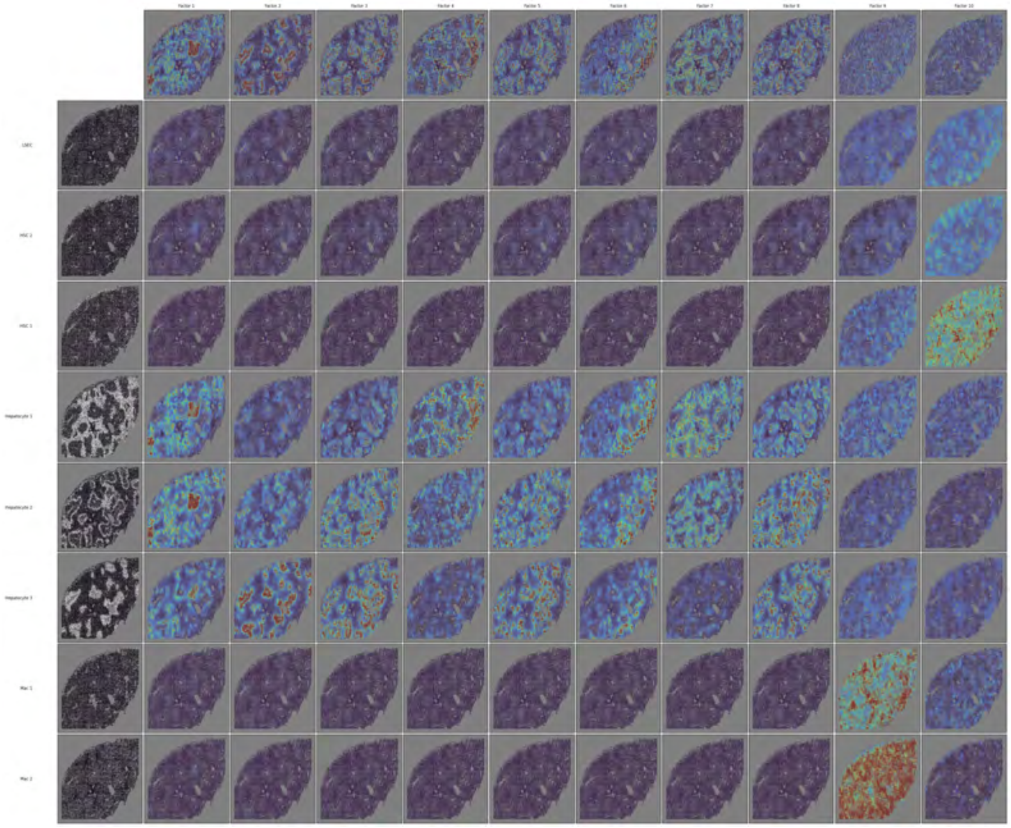

**Fig 25. MGGP-LCGP cell-type-conditional spatial factors for MERFISH human liver — healthy sample AM042.** Full grid of cell-type-conditional spatial factors for the healthy AM042 liver sample.

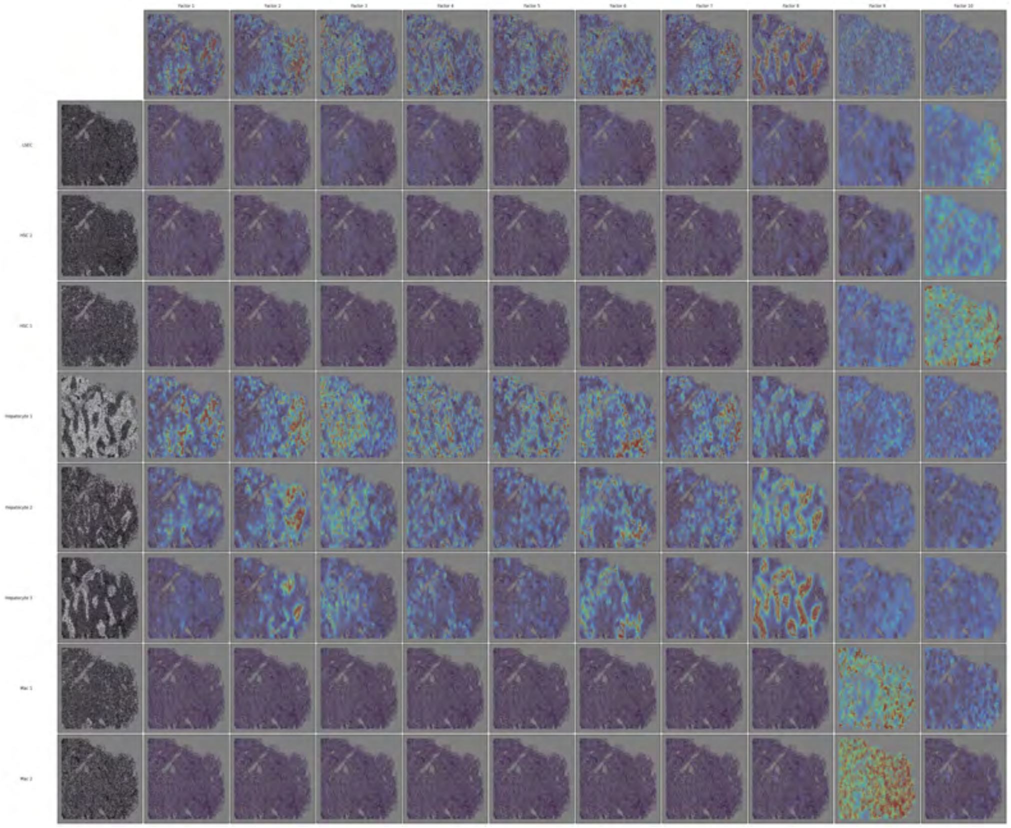

**Fig 26. MGGP-LCGP cell-type-conditional spatial factors for MERFISH human liver — healthy sample AM061.** Full grid of cell-type-conditional spatial factors for the healthy AM061 liver sample.

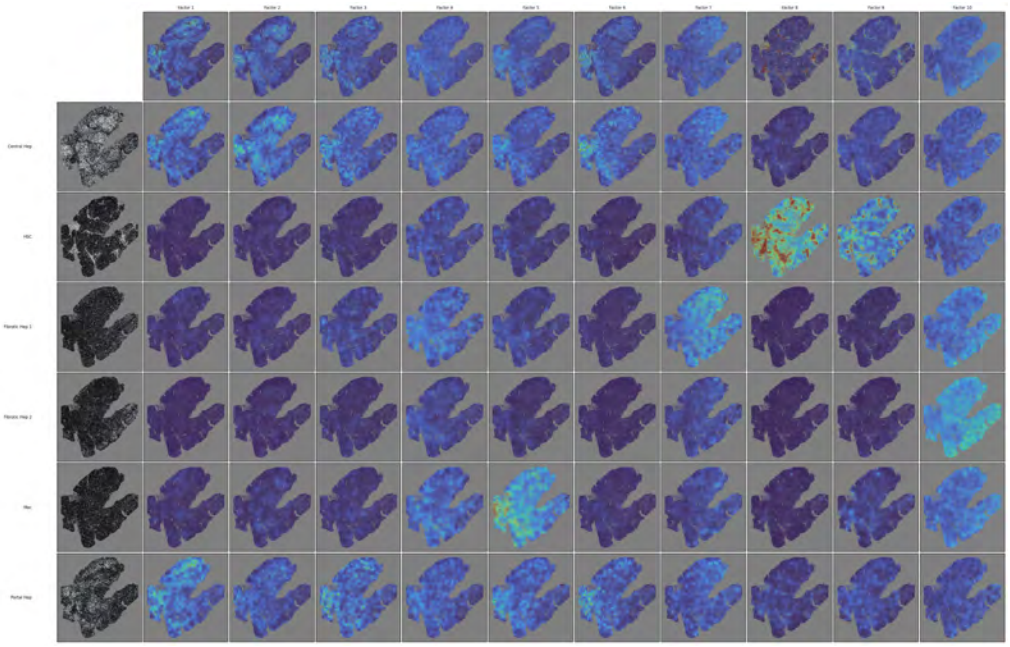

**Fig 27. MGGP-LCGP cell-type-conditional spatial factors for MERFISH human liver — diseased sample AM031.** Full grid of cell-type-conditional spatial factors for the diseased AM031 liver sample.

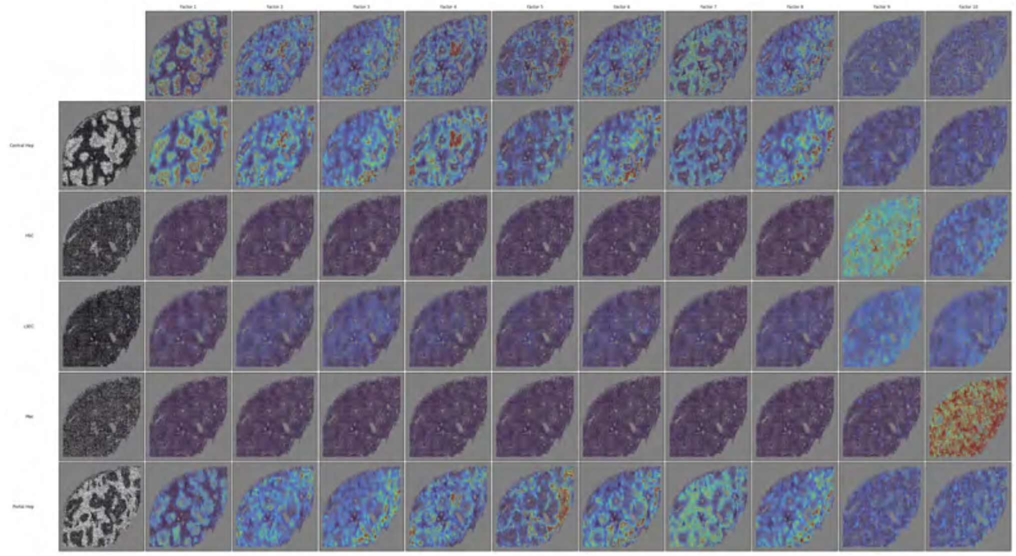

**Fig 28. MGGP-LCGP cell-type-conditional spatial factors for MERFISH human liver — diseased sample AM042.** Full grid of cell-type-conditional spatial factors for the diseased AM042 liver sample.

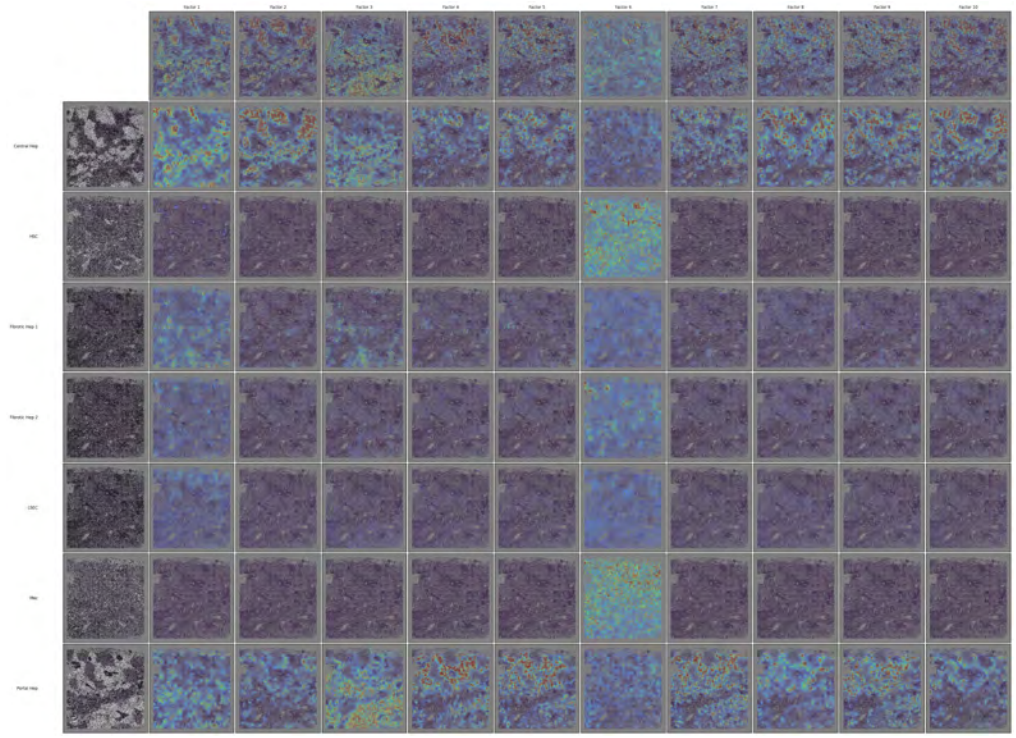

**Fig 29. MGGP-LCGP cell-type-conditional spatial factors for MERFISH human liver — diseased sample AM048.** Full grid of cell-type-conditional spatial factors for the diseased AM048 liver sample.

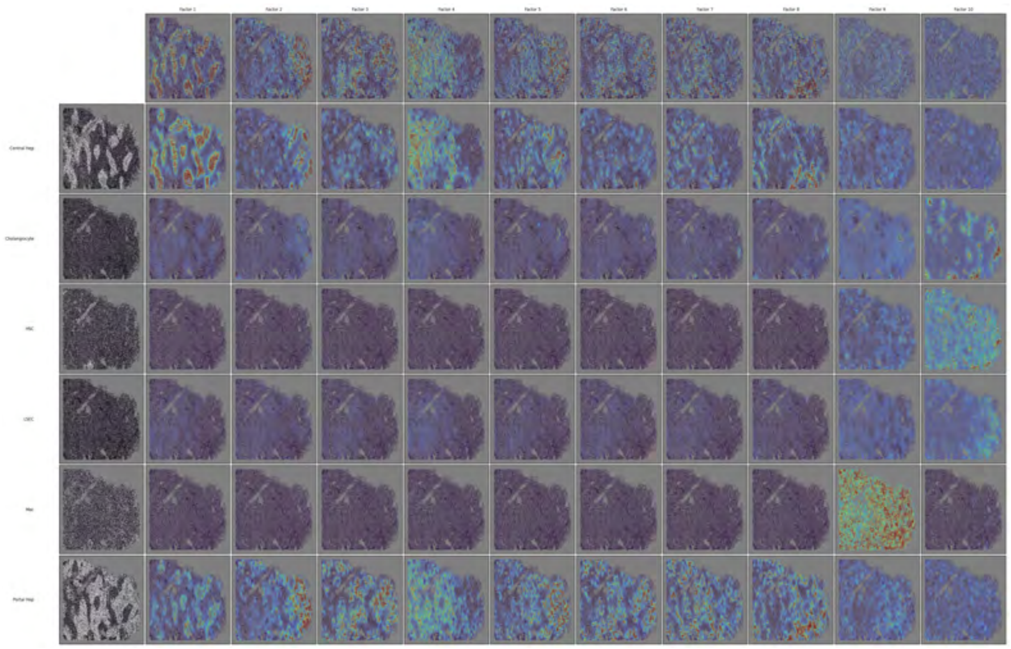


**Fig 30. MGGP-LCGP cell-type-conditional spatial factors for MERFISH human liver — diseased sample AM061.** Full grid of cell-type-conditional spatial factors for the diseased AM061 liver sample.

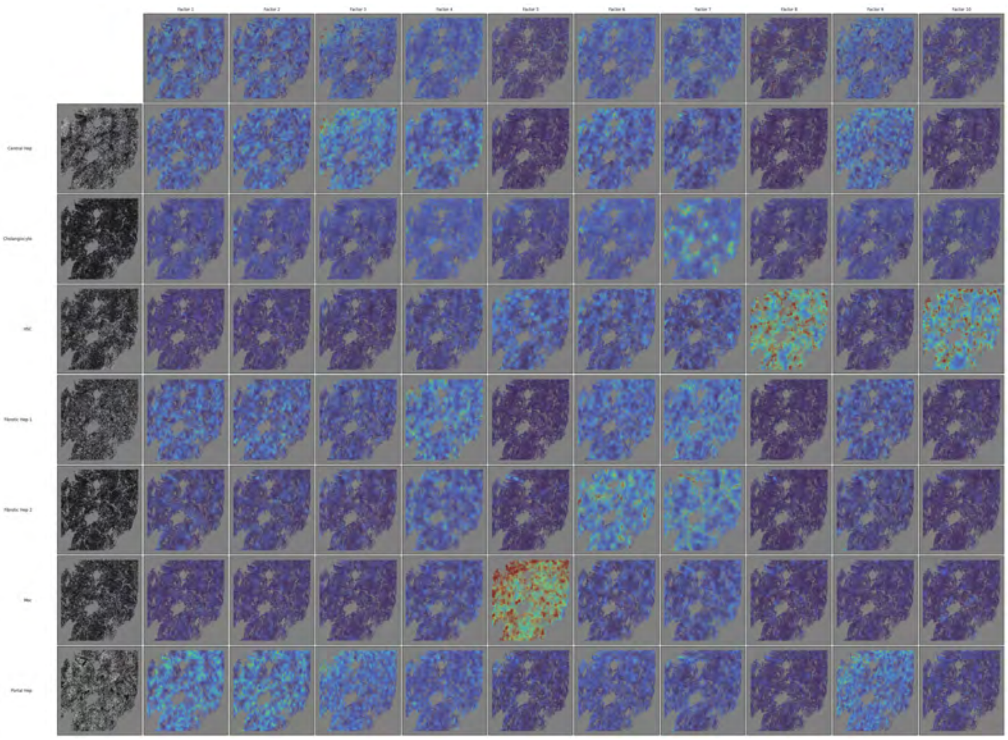

**Fig 31. MGGP-LCGP cell-type-conditional spatial factors for MERFISH human liver — diseased sample AM062.** Full grid of cell-type-conditional spatial factors for the diseased AM062 liver sample.

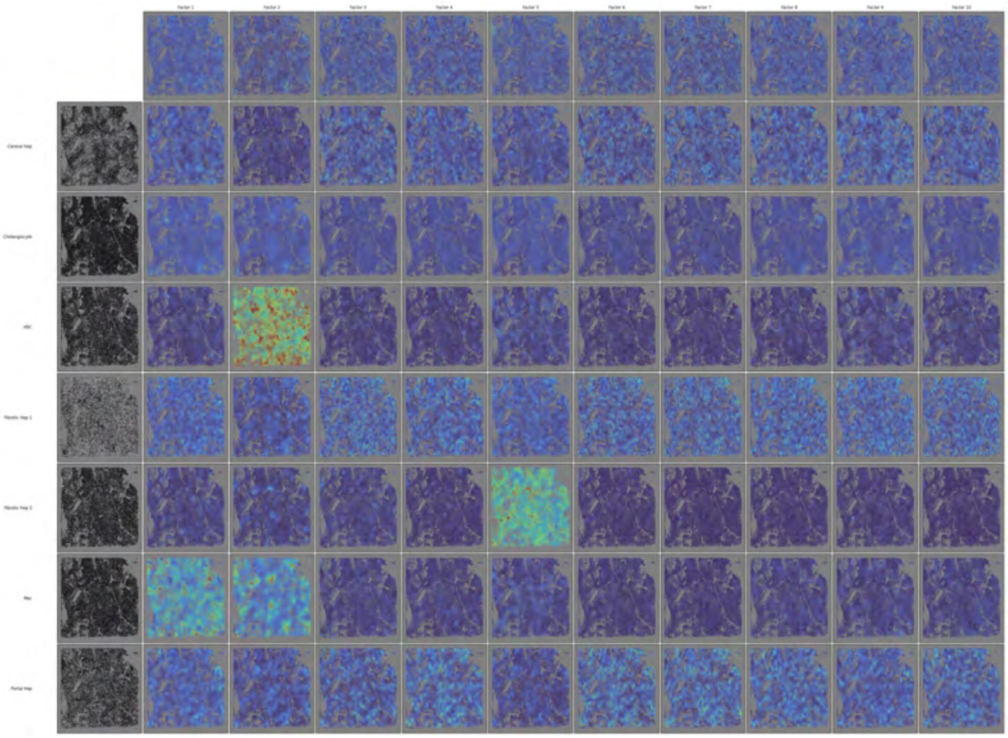

**Fig 32. MGGP-LCGP cell-type-conditional spatial factors for MERFISH human liver — diseased sample AM072.** Full grid of cell-type-conditional spatial factors for the diseased AM072 liver sample.

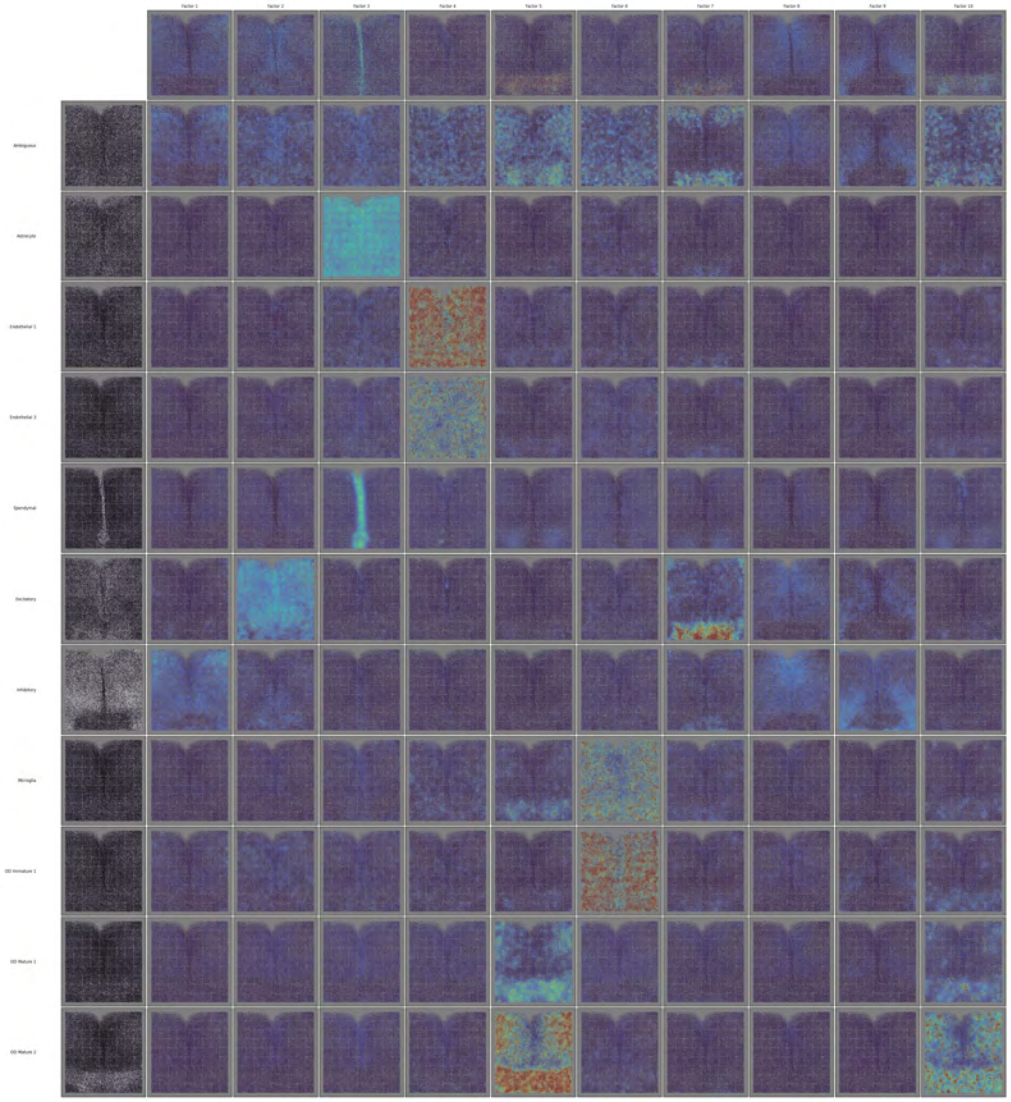


**Fig 33. MGGP-LCGP cell-type-conditional spatial factors for MERFISH mouse hypothalamus.** Full grid of cell-type-conditional spatial factors for the MERFISH hypothalamus dataset (Moffitt2018; 71,939 cells, 161 genes, 11 groups).

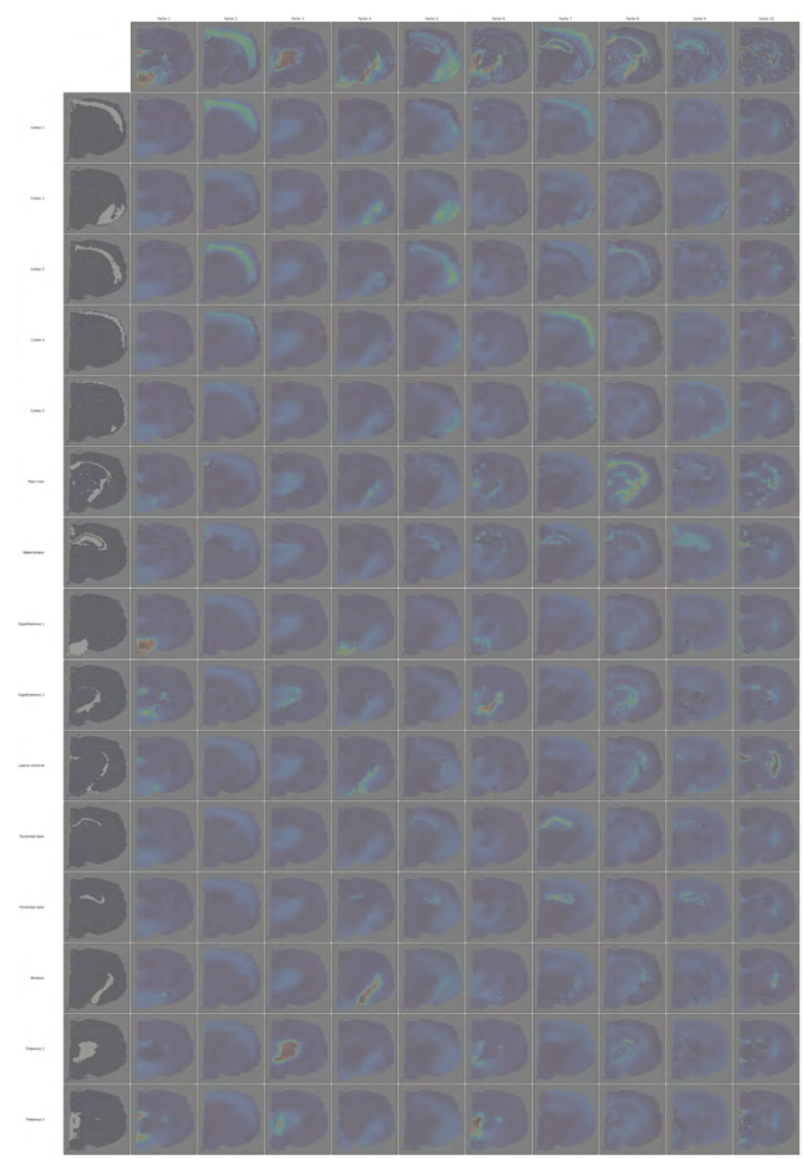

**Fig 34. MGGP-LCGP group-conditional spatial factors for 10x Visium Adult Mouse Brain (Coronal Section 1).** Full grid of group-conditional spatial factors for the 10x Visium mouse brain slice (2,688 spots, 16,944 genes, 15 brain region groups). Complements the curated subset shown in the Visium main-text panel.

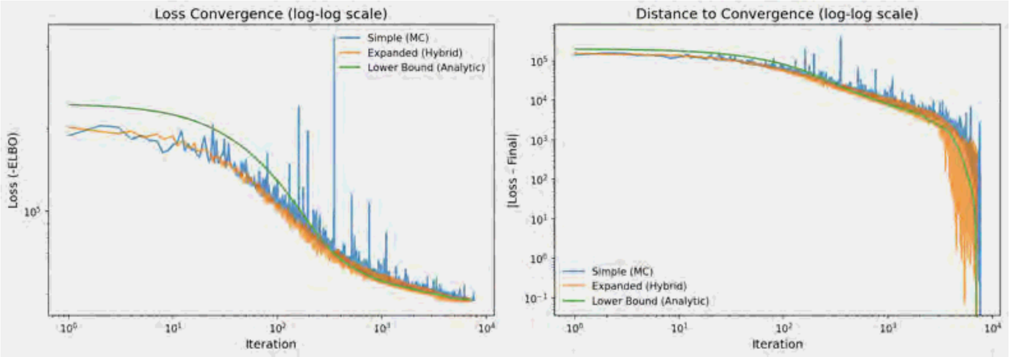


**Fig 35. Convergence of the three multiplicative-update methods on a synthetic nonnegative-matrix benchmark.** Side-by-side comparison of the Jensen lower bound (Method 1), the expanded/hybrid estimator (Method 2), and full Monte Carlo (Method 3) when factorizing a random nonnegative matrix into $L = 10$ components. The expanded estimator is substantially more stable than full Monte Carlo while attaining a lower final loss than the analytic lower bound, motivating its use as the default in all spatial experiments.

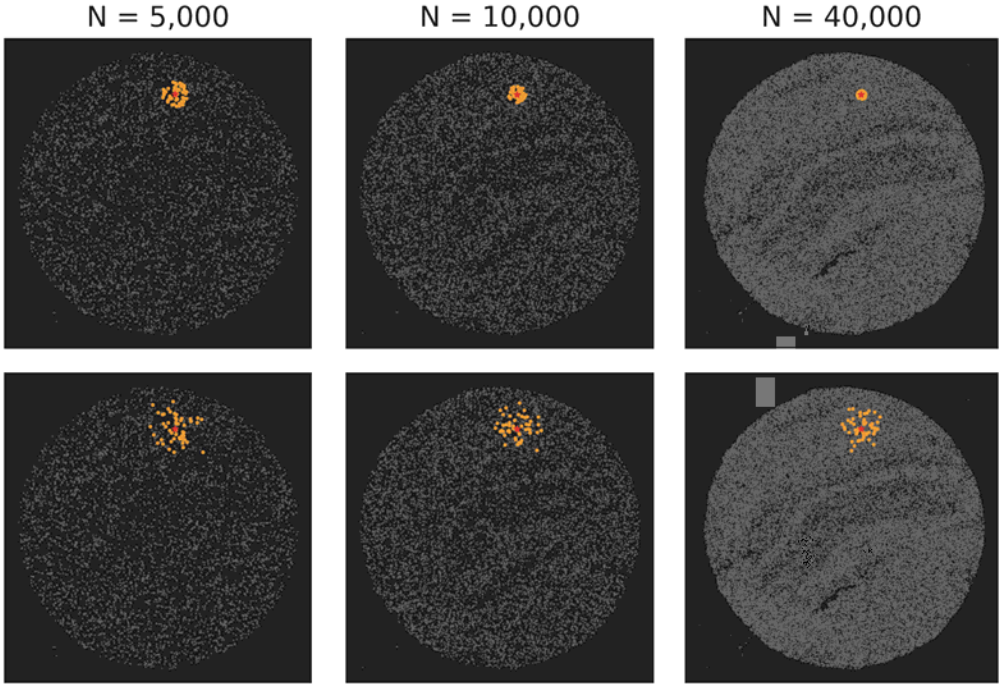


**Fig 36. Kernel-conditioned neighbor selection remains stable as data density grows.** Each panel shows the $K$=50 neighbors (orange) of the same query point in subsamples of the Slide-seqV2 hippocampus dataset at $N \in \{5{,}000,\ 10{,}000,\ 40{,}000\}$ cells. **Top row:** baseline Euclidean KNN—the neighborhood radius collapses with increasing $N$. **Bottom row:** LCGP with kernel-conditioned neighbor selection (RBF kernel, lengthscale $\ell$=8.0)—the neighborhood extent stays well-behaved across all densities.

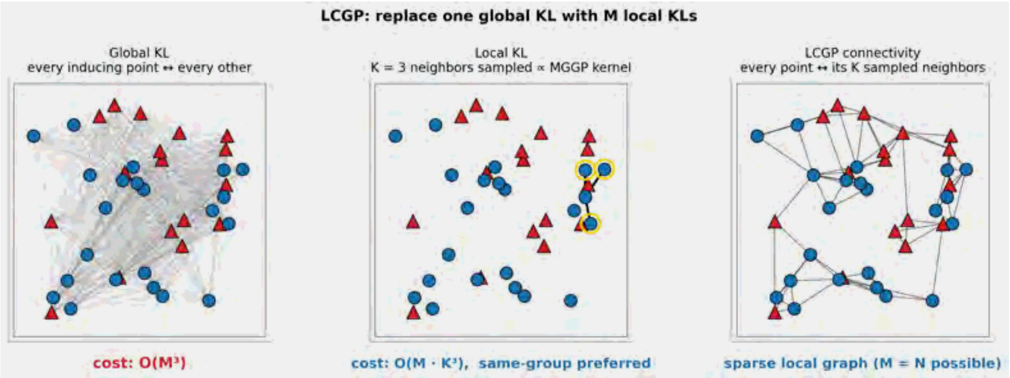


**Fig 37. LCGP replaces the global KL with $M$ local KLs.** *Left:* the exact MGGP-SVGP KL connects every inducing point to every other and costs $\mathcal{O}(M^3)$. *Middle:* LCGP samples a neighbor set $n(j)$ of size $K$ for each inducing point $j$ with probability proportional to the MGGP kernel $k\big((z_j, c_j), (z_i, c_i)\big)$; same-group neighbors (blue circles) are preferred but cross-group neighbors (red triangles) enter the set whenever the kernel ranks them highly. *Right:* repeating this sampling at every inducing point yields the LCGP connectivity graph—a sparse graph whose local KL costs $\mathcal{O}(MK^3)$, making $M = N$ inducing points tractable for the KL term.

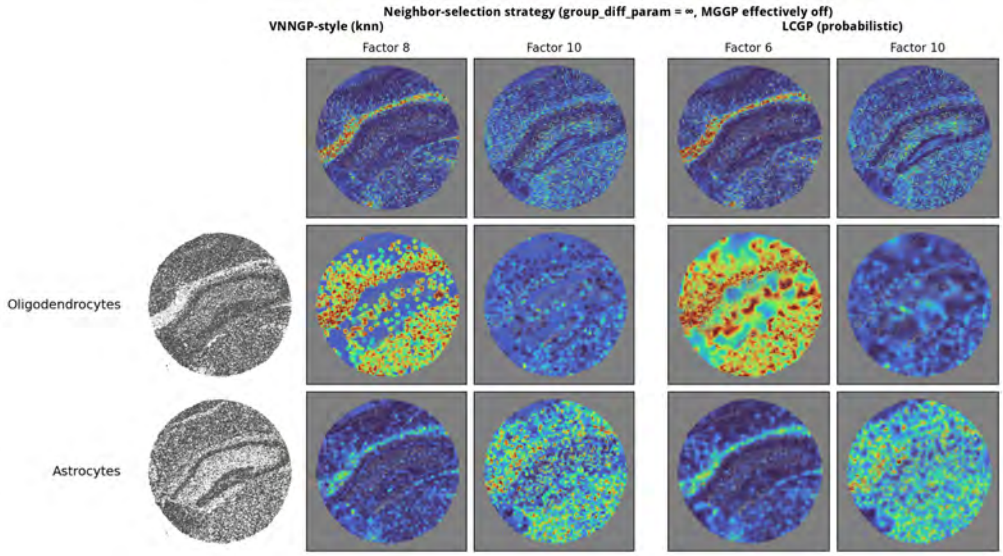


**Fig 38. Probabilistic vs. $K$-nearest-neighbor selection in LCGP on Slide-seqV2 hippocampus.** Both blocks are MGGP-LCGP fits at $a = 10^6$ (so the multi-group coupling is effectively removed and the only remaining difference is the neighbor-selection rule). Each block shows the unconditional spatial factor map (top row) and two cell-type-conditional posteriors below; the leftmost column of each row marks which cells belong to that group. Reference factors and cell-type rows are picked from the VNNGP/KNN side (left block): the top two factors by max-over-groups $L_1$ specificity ratio, with each factor's argmax-enriched cell type as its row (Oligodendrocytes, Astrocytes). The corresponding columns of the LCGP/probabilistic block are not the same numerical factor indices but are the best Pearson matches on the per-factor gene-loading vectors, so that the same biological gene program is being compared across methods. Driving selection from the VNNGP side is the conservative framing: the baseline picks the factors and cell-type rows where it shows the strongest signal, so the LCGP block's differences (more anatomically structured conditional maps on the same rows) cannot be attributed to LCGP-favorable factor selection.
